# Pathway selection for arabinose utilization in *Pseudomonas putida* reveals a rate-yield tradeoff in muconic acid production from lignocellulosic sugars

**DOI:** 10.64898/2026.07.14.738590

**Authors:** Dowan Kim, Torrey M. Lind, Chen Ling, Bruno C. Klein, Ashley N. Merrill, Elisabeth Van Roijen, Pahola Thathiana Benavides, Alexander F. Benson, Joshua R. Elmore, Morgan A. Ingraham, Eugene Kuatsjah, Nicolette R. Meyer, Sekgetho C. Mokwatlo, Kelsey J. Ramirez, Adam M. Guss, Alissa C. Bleem, Davinia Salvachúa, Christopher W. Johnson, Gregg T. Beckham

## Abstract

Engineering heterologous utilization of substrates requires selection of catabolic pathways that balance strain performance and product biosynthesis. Here, we compare the oxidative and isomerase arabinose utilization pathways in *Pseudomonas putida* strains engineered for *cis,cis*-muconic acid production from glucose and xylose. Based on the point of entry into central carbon metabolism, we hypothesized that the oxidative arabinose pathway would enable higher productivity while the arabinose isomerase pathway would enable higher muconate yield. In both strains, additional modifications were engineered to improve muconic acid production including sugar transporter tuning, catechol 1,2-dioxygenase overexpression, a feedback-resistant DAHP synthase, and a flux-stabilizing *gltA* variant. Consistent with our hypothesis, the oxidative arabinose pathway supported faster growth and higher productivity (0.58 g/L/h), whereas the arabinose isomerase pathway improved carbon efficiency, achieving muconate yields of up to 50 C-mol% in fed-batch bioreactors. Process modeling indicates that these performance metrics can reduce the minimum selling price of muconate-derived adipic acid to $2.74/kg and greenhouse gas emissions to 1.31 kg CO_2_e/kg, approaching cost parity and reducing emissions by 86% relative to fossil carbon-derived adipic acid. Overall, this study presents a systematic comparison of sugar catabolic pathways that enabled development of strains suited for the tradeoffs between rate and yield.

## Introduction

Lignocellulosic biomass is a renewable feedstock for biomanufacturing (*1–4*), typically yielding D-glucose, D-xylose, and L-arabinose as the predominant carbohydrates (*5, 6*). While extensive engineering has enabled efficient glucose and xylose co-utilization in diverse microbes, the complete valorization of these three sugars remains limited (*7–9*). The underutilized arabinose fraction in sugar hydrolysates (~5-10 %) not only reduces overall theoretical carbon yields but also negatively impacts biorefining economics. L-Arabinose assimilation typically proceeds via two distinct routes: the oxidative pathway (e.g., in *Burkholderia ambifaria*), which directs arabinose to the tricarboxylic acid (TCA) cycle via α-ketoglutarate (*10*), or the isomerase pathway (e.g., in *Escherichia coli*), which introduces arabinose into the pentose phosphate (PP) pathway via xylulose-5-phosphate (X5P) (*11*). Because these pathways enter central carbon metabolism at different points, the oxidative pathway favors ATP/NADH production, and the isomerase pathway conserves carbon for biosynthetic precursors; therefore, these two pathways can offer distinct metabolic tradeoffs between growth, product biosynthesis rates, and product yield.

*cis,cis*-Muconic acid (hereafter muconate) is a valuable platform chemical that can be converted to incumbent polymer precursors such as adipic acid, terephthalic acid, hexamethylenediamine, and caprolactam or used directly in performance-advantaged bioproducts (*3, 12–21*). Muconate serves as an intermediate in aromatic catabolism in various microbes (*22–24*), and *Pseudomonas putida* KT2440 (*P. putida*) has been extensively engineered to produce muconate due to its metabolic versatility and robustness in utilizing carbohydrates (*14, 25–27*), lignin-derived aromatics (*16, 25, 28–31*), and plastics-derived aromatics (*31–35*). To produce muconate from glucose, *P. putida* utilizes the EDEMP cycle, a metabolic architecture that combines the Entner-Doudoroff (ED), the Embden-Meyerhof-Parnas (EMP), and the PP pathways to generate phosphoenolpyruvate (PEP) and erythrose 4-phosphate (E4P) (**Fig. 1**) (*36*). These precursors are then channeled through the shikimate pathway toward muconate, based on the pioneering work of Draths and Frost (*12, 37*). While recent efforts have enabled co-utilization of hexose and pentose sugars in *P. putida* (*38–40*), achieving high muconate productivity remains a challenge. For example, our benchmark strain, *P. putida* LC224, exhibited a muconate titer of 33.7 g/L, a productivity of 0.18 g/L/h, and a C-mol yield of 46% (*27*). This strain, however, was incapable of utilizing arabinose, the third most abundant sugar in hydrolysates derived from corn stover and many other grass-based lignocellulosic feedstocks (*6, 9, 38*).

**Figure 1.**
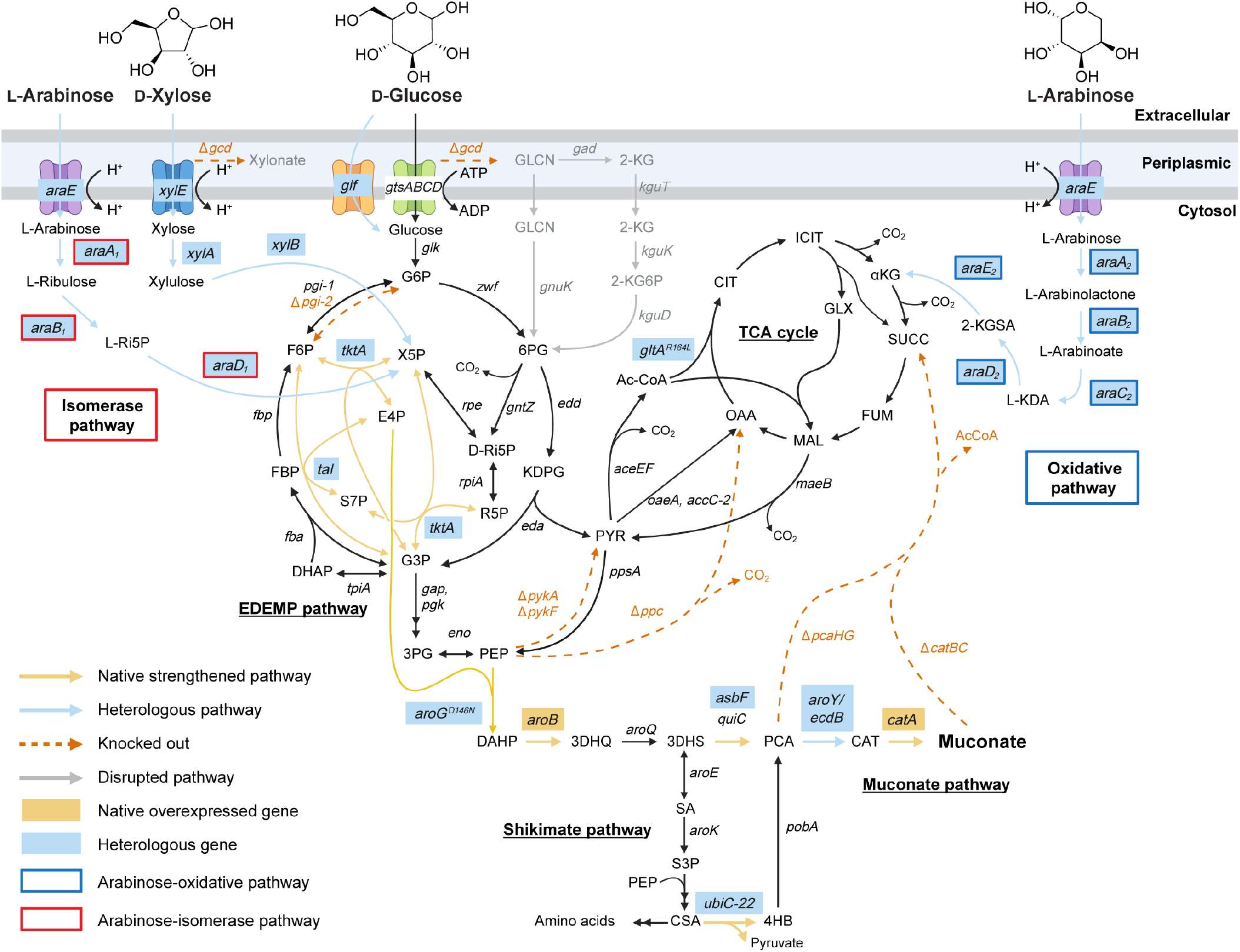
Muconate production from lignocellulosic sugars. Metabolic pathways enabling *P. putida* to assimilate sugars from lignocellulose hydrolysates (molar ratio of 63.1% glucose, 31.5% xylose, and 5.4% arabinose) (*6*) for muconate production. Two heterologous L-arabinose assimilation pathways are shown: the isomerase pathway (red box) entering the pentose phosphate pathway, and the oxidative pathway (blue box) entering the TCA cycle. Arrows represent the pathway by indicated gene products: black, native pathway; yellow, native strengthened pathway; blue, heterologous pathway; red dashed, knocked-out pathway; gray, disrupted pathway. Yellow-labeled boxes indicate native genes overexpressed and blue-labeled boxes indicate heterologous genes introduced. Abbreviations are listed in **Table S1**.

Given that muconate biosynthesis depends on the stoichiometric supply of PEP and E4P, we hypothesized that the entry point of arabinose into central metabolism could dictate production performance with distinct tradeoffs between productivity and yield (**Fig. 1**). Because the benchmark strain, *P. putida* LC224 has partially disconnected glycolysis from the TCA cycle via deletion of the pyruvate kinase genes (*pykA, pykF*), routing arabinose through the oxidative pathway into the TCA cycle is expected to enhance energy generation and biomass formation, thereby increasing the growth rate and, consequently, productivity (**Fig. 1**).

In contrast, routing arabinose through the isomerase pathway into X5P is expected to increase carbon flux toward the PP pathway and the shikimate pathway, increasing the availability of PEP and E4P and improving muconate yield at the potential expense of growth rate (**Fig. 1**). To our knowledge, while these routes have been independently implemented in various hosts (*10, 41–45*), a systematic comparison of their performance tradeoffs regarding target molecule biosynthesis has not been reported (*46, 47*).

In this study, we engineered *P. putida* to metabolize arabinose via these two distinct pathways alongside glucose and xylose to enhance muconate production (**Fig. 1**). Through rational strain engineering and adaptive laboratory evolution, we resolved several metabolic bottlenecks, slow steps in metabolism that throttle growth and/or production and often cause the accumulation of metabolic intermediates. These efforts included introducing a glucose-facilitated diffusion transporter from *Zymomonas mobilis* (*48*), overexpressing catechol 1,2-dioxygenase, integrating an additional copy of a feedback-resistant DAHP synthase (AroG^D146N^) gene (*49*), and fine-tuning TCA cycle flux via a feedback-resistant GltA^R164L^ variant (*50, 51*). Our optimized strains achieved complete co-utilization of glucose, xylose, and arabinose with industrially relevant performance. Specifically, the oxidative arabinose pathway strain reached a muconate titer of 41.3 g/L, a productivity of 0.58 g/L/h, and a yield of 48.1 C-mol % in fed-batch bioreactors, while the isomerase pathway strain achieved 38.2 g/L, 0.54 g/L/h, and 50.2 C-mol %. Finally, we performed techno-economic analysis (TEA) and life cycle assessment (LCA) to evaluate how these pathway-dependent tradeoffs influence the economic viability and environmental impact of muconate-derived adipic acid production.

## Results

### Engineering two L-arabinose pathways to enable co-utilization of glucose, xylose, and arabinose

To enable muconate production from all three carbohydrates in lignocellulosic hydrolysates, genes encoding two distinct L-arabinose utilization pathways were chromosomally integrated into strain LC224 as shown in **Fig. 1** and **Extended Data Table 1**, which was previously engineered for glucose and xylose conversion (*27*). For arabinose uptake, we first introduced genes encoding the arabinose-H^+^ symporter AraE from *E. coli*, and the oxidative pathway was subsequently constructed by integrating five genes encoding the L-arabinose dehydrogenase (AraA_2_), L-arabinolactonase (AraB_2_), L-arabinonate dehydratase (AraC_2_), 2-keto-3-deoxy-L-arabinoate dehydratase (AraD_2_), and α-ketoglutarate semialdehyde dehydrogenase (AraE_2_) into strain LC224, which we had previously optimized for conversion of glucose and xylose to muconate (*27*), generating strain LC237 (**Fig. 1, Extended Data Table 1**). In parallel, the isomerase pathway was constructed by introducing three genes encoding the L-arabinose isomerase (AraA_1_), ribulokinase (AraB_1_), and L-ribulose-5-phosphate 4-epimerase (AraD_1_) into LC224, generating strain LC357 (**Fig. 1, Extended Data Table 1**).

To test these engineered pathways, we first evaluated strains LC237 and LC357 along with their parent strain LC224 using a plate reader in M9 medium supplemented with 30 mM arabinose, 30 mM glucose + 2.5 mM arabinose, or mock hydrolysate containing 30 mM glucose + 15 mM xylose + 2.5 mM arabinose), similar sugar ratios to those found in corn stover-derived sugar hydrolysates (**Fig. S1**) (*6*). Both LC237 and LC357 grew in M9 medium with arabinose as the sole carbon and energy source, confirming the arabinose pathways were functional, as shown in **Fig. 2a**. In mixed-sugar media (**Fig. S1**), LC237 exhibited a reduced lag in growth compared to LC224, likely driven by more rapid energy generation from the arabinose-oxidative pathway, while LC357 displayed a longer lag and slower growth rate. Shake-flask experiments in mock hydrolysate corroborated the growth profiles observed in the plate reader assays. Notably, while all strains produced comparable muconate yields and titers, LC237 exhibited superior growth kinetics (**Fig. 2b-d, Fig. S2**). LC237 reached maximum growth (measured as optical density at 600 nm, OD_600_) within 24 h, whereas LC224 and LC357 required 42 h and 67 h, respectively. This rapid growth resulted in a higher productivity (calculated from the final titer point) of 0.43 mM/h for LC237, compared to 0.40 mM/h and 0.26 mM/h for LC224 and LC357, respectively. These results were an early indication that the arabinose-oxidative pathway can accelerate growth and muconate productivity.

**Figure 2.**
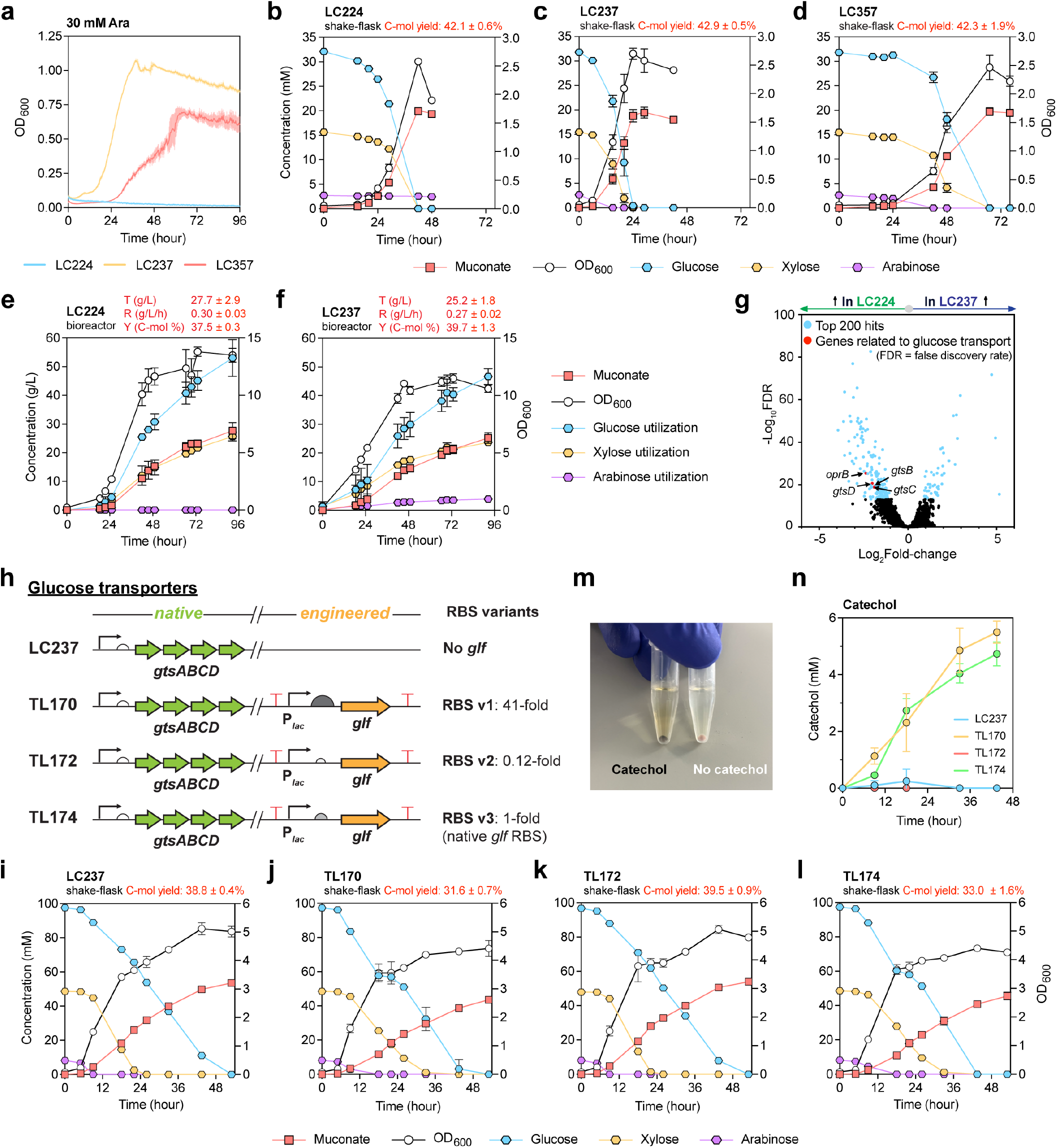
Comparison of arabinose catabolic pathways and identification of bottlenecks in glucose uptake and catechol ring-opening in the oxidative pathway strain. **a** Growth curves of LC224, LC237 (LC224 w/ arabinose-oxidative pathway) and LC357 (LC224 w/ arabinose-isomerase pathway). Strains were grown in a plate reader on M9 medium supplemented with 30 mM arabinose. **b-d** Shake-flask experiment profiles showing the bacterial growth, residual sugar concentrations, and muconate production of LC224 (**b**), LC237 (**c**), and LC357 (**d**) on M9 medium supplemented with 8 g/L mock hydrolysate (see **Fig. S2** for comparative profiles). **e-f** Fed-batch profiles from cultivations in 0.5-L bioreactors, showing the bacterial growth, total sugar utilization, and muconate production of LC224 (**e**) and LC237 (**f**). Additional bioreactor profiles, including residual sugars in bioreactors during the cultivations, yields, and productivities are provided in **Fig. S3. g** Transcriptomic analysis of LC224 and LC237 grown on 40 g/L mock hydrolysates. The volcano plot shows genes that exhibited significant differences in expression level (p < 0.005, log_2_ fold-change > |1|), with the top 200 hits in blue and glucose transport-related genes in red. **h** Schematic of glucose transport systems in engineered strains. Native glucose transporter: GtsABCD, engineered glucose facilitator: Glf. The *glf* gene is expressed under the *lac* promoter with RBS variants of different targeted translation initiation rates (TIRs), generating TL170, TL172, and TL174. **i-l** Shake-flask experiment profiles showing the bacterial growth, residual sugar concentrations, and muconate production of LC237 (**i**), TL170 (**j**), TL172 (**k**), and TL174 (**l**) on M9 medium supplemented with 25 g/L mock hydrolysate (see **Fig. S7** for comparative profiles). **m** Centrifuged cell-pellets of TL170 (*left*, catechol accumulation) and LC237 (*right*, no catechol accumulation). **n** Catechol accumulation during shake-flask cultivation of LC237, TL170, TL172, and TL174 on 25 g/L mock hydrolysate. Concentrations are reported in mM for flasks and g/L for bioreactors to accommodate low and high ranges of substrates and products, respectively. Titer: final muconate concentration (g/L), Rate: titer at the final time point/time (g/L/h), Yield (C-mol %): [(mM muconate × 6) / mM (glucose × 6 + mM xylose × 5 + mM arabinose × 5) × 100%] calculated at the final time point. Data represent the average of *n* = 3 biological replicates in **b**-**d, i**-**l, n** and *n* = 2 biological replicates in **e-h.** Error bars correspond to standard deviation in **b-d, i**-**l, n** and absolute error between duplicates in **e-h**. Numerical data are provided in a **Source Data File**.

Given the superior growth and production kinetics observed with incorporation of the oxidative pathway, we prioritized evaluation of strain LC237 in 0.5-L fed-batch bioreactors compared to the parent strain, LC224 (**Fig. 2b-d**). Mock hydrolysate was used for the batch and feeding phases (16.8 g/L glucose, 7 g/L xylose, and 1.2 g/L of arabinose for batch and 336 g/L of glucose, 140 g/L of xylose, 23 g/L of arabinose for feeding) (*6*). All the experiments with mock hydrolysate were conducted with this composition hereafter. Notably, LC237 showed faster initial growth than LC224 until 40 h, but its final growth and muconate titer were ultimately lower, reaching 25.2 g/L muconate compared to 27.7 g/L in LC224 (**Fig. 2e-f, Fig. S3**). This decline in performance correlated with markedly reduced glucose consumption after 40 h, suggesting a potential metabolic bottleneck in glucose metabolism in LC237 under bioreactor conditions. We next sought to address this bottleneck.

### Identifying the glucose uptake bottleneck

To investigate cellular mechanisms for the reduced glucose consumption rates in LC237 during conversion of mock hydrolysates in **Fig. S3**, we performed transcriptomic profiling of LC237 and the parent strain LC224 at mid-log phase in shake flasks. Among the 5,570 transcripts identified, 556 genes were upregulated and 441 genes were downregulated by at least two-fold (|log2 fold-change| > 1) in LC237 relative to LC224 (**Fig. 2g**, see **Supplementary File** for details). Notably, the glucose transporter operon *gtsABCD* (PP_1015-1018) and the outer membrane porin *oprB-I* (PP_1019) were among the most strongly downregulated genes in LC237. The gene *oprB-I* encodes a porin involved in glucose uptake across the outer membrane, while *gtsABCD* encodes a high-affinity inner membrane glucose transporter complex (*52*). Given their essential roles in glucose uptake, we reasoned that impaired glucose transport might represent a bottleneck limiting muconate production.

To further investigate the impact of arabinose-oxidative pathway on overall sugar uptake, we also examined the importance of the newly introduced arabinose-H^+^ symporter, AraE, on the uptake of arabinose and other sugars. Because AraE uses the proton motive force (PMF), high levels of AraE activity may reduce available PMF, potentially impairing ATP generation under energy-limited conditions (*53*) (**Supplementary Note 1**). Thus, we posited that its expression might interfere with the uptake of other sugars. To test this, we deleted *araE* from LC237, generating TL015. Notably, this *araE* deletion strain exhibited faster growth on xylose than LC237 in plate-reader assays (**Fig. S4**), confirming that AraE activity inhibits utilization of xylose. Interestingly, AraE deletion delayed but did not prevent growth on arabinose (**Fig. S4**), suggesting possible transport promiscuity by native or heterologous transporters (*42*). We also found that downregulating *araE* expression slightly improved glucose consumption in the shake-flask experiment, also confirming our hypothesis, but it simultaneously impaired xylose and arabinose consumption (**Fig. S5**). Together, while AraE impairs glucose and xylose utilization, its removal or attenuation cannot fully resolve the sugar uptake bottleneck.

### Harnessing the Glf glucose facilitator to resolve the glucose uptake bottleneck

Given the reduced glucose metabolism and likely cellular burden associated with AraE, we reasoned that overexpressing the genes encoding the native ATP-dependent glucose transporter (GtsABCD) might place additional demand on a strain already operating under a constrained energy budget. We therefore evaluated use of the glucose facilitator (Glf) from *Zymomonas mobilis*, which mediates passive transport via facilitated diffusion (*48, 54*), reasoning that this approach could offer a more energy-efficient means of enabling glucose uptake. We first designed constructs for the constitutive expression of *glf* in strain LC237 using the *tac* promoter (P_*tac*_), which drives strong, constitutive expression in *P. putida (55, 56)*. Despite multiple attempts, genomic integration of *glf* under P_*tac*_ was unsuccessful, suggesting that this level of *glf* expression may be toxic.

We therefore tuned *glf* expression by using a medium-strength promoter (P_*lac*_) with variable ribosome binding sites (RBSs). Using the Salis RBS Calculator (*57*), three RBS variants (v1: 41-fold, v2: 0.12-fold, v3: 1-fold from native *glf* RBS, **Extended Data Table 1**) were designed and integrated with the *glf* gene into LC237, resulting in strains TL170, TL172, and TL174, respectively (**Fig. 2h**). When evaluated on mock hydrolysate using a plate reader, TL170 and TL174 improved growth rates and reduced lag times compared to LC237, whereas TL172 showed no significant improvement (**Fig. S6**). Notably, TL170 and TL174 cultures also exhibited a pronounced dark brown color, which was absent in cultures of LC237 and TL172, consistent with the accumulation of catechol in the medium, a metabolic pathway intermediate directly upstream of muconate. We posited that this accumulation, coupled with the improved growth, resulted from enahnced glucose utilization in TL170 and TL174.

To determine if catechol oxidation to muconate was the new bottleneck, we next conducted shake-flask experiments using M9 medium supplemented with mock hydrolysate (25 g/L total sugars) matching the sugar loading used in the bioreactor conditions in the batch phase. Compared to LC237, strains TL170 and TL174 showed markedly improved glucose utilization, with nearly complete glucose depletion by 44 h and enhanced consumption within the first 18 h, as shown in **Fig. 2i-l** and **Fig. S7**. As observed in the plate reader assays, cultures of TL170 and TL174 exhibited the same brown coloration by 24 h, whereas LC237 and TL172 cultures did not (**Fig. 2m** and **Fig. S8**). HPLC analysis confirmed catechol accumulation to 5.5 mM in TL170 and 4.7 mM in TL174, while LC237 peaked at 0.7 mM before catechol disappeared and TL172 showed no measurable catechol accumulation (**Fig. 2n**). These findings suggest that *glf* expression increased glucose transport and metabolic flux into the shikimate pathway, and this resulted in catechol accumulation as a new bottleneck. Given the toxicity of catechol in metabolism (*16*), we therefore reasoned that increasing the expression of *catA1*, encoding one of two native catechol 1,2-dioxygenases in *P. putida*, could relieve this constraint (*58, 59*).

### Reducing catechol accumulation by increasing CatA expression

TL170 exhibited the fastest growth rate and rapid glucose uptake among the engineered strains, so we selected it for further engineering to mitigate catechol accumulation (**Fig. 3a**). Since *catA1* was already expressed using the strong *tac* promoter, we opted to improve its translation efficiency through RBS engineering. Of the designed RBS variants (predicted to be about 10-fold and 100-fold higher translation initiation rates (TIRs) relative to the native RBS), only the 10-fold variant was successfully integrated, yielding strain TL207 (**Extended Data Table 1**). To assess whether this RBS modification increased CatA1 protein abundance and, subsequently, activity, we performed specific activity measurements on crude cell extracts of the different engineered *P. putida* strains using oxygraphy, as shown in **Fig. 3b**. Because one mole of oxygen is consumed per mole of catechol converted to muconate via CatA-catalyzed ring cleavage, oxygen consumption served as a proxy for CatA activity. However, as crude extracts may contain additional oxygen-consuming enzymes, oxygen uptake should be interpreted as an indirect estimate of CatA activity rather than a direct kinetic measurement. Strain TL207 exhibited a ~7-fold increase in oxygen consumption compared to both LC237 and TL170, suggesting a substantial elevation in CatA1 enzyme abundance. In a plate reader assay, TL207 showed no visible dark brown coloration in M9 medium, while maintaining growth rates comparable to TL170 (**Fig. S6**). In shake-flask experiments on mock hydrolysate, TL207 consumed glucose more rapidly than LC237 with similar growth and muconate production (**Fig. S9**). Compared to TL170, TL207 achieved a higher muconate titer and rate, without catechol accumulation, as shown in **Fig. 3c** and **Fig. S9**. These results indicate that RBS engineering of *catA1* effectively alleviated catechol accumulation while retaining the benefits of the previous engineering interventions to improve glucose uptake.

**Figure 3.**
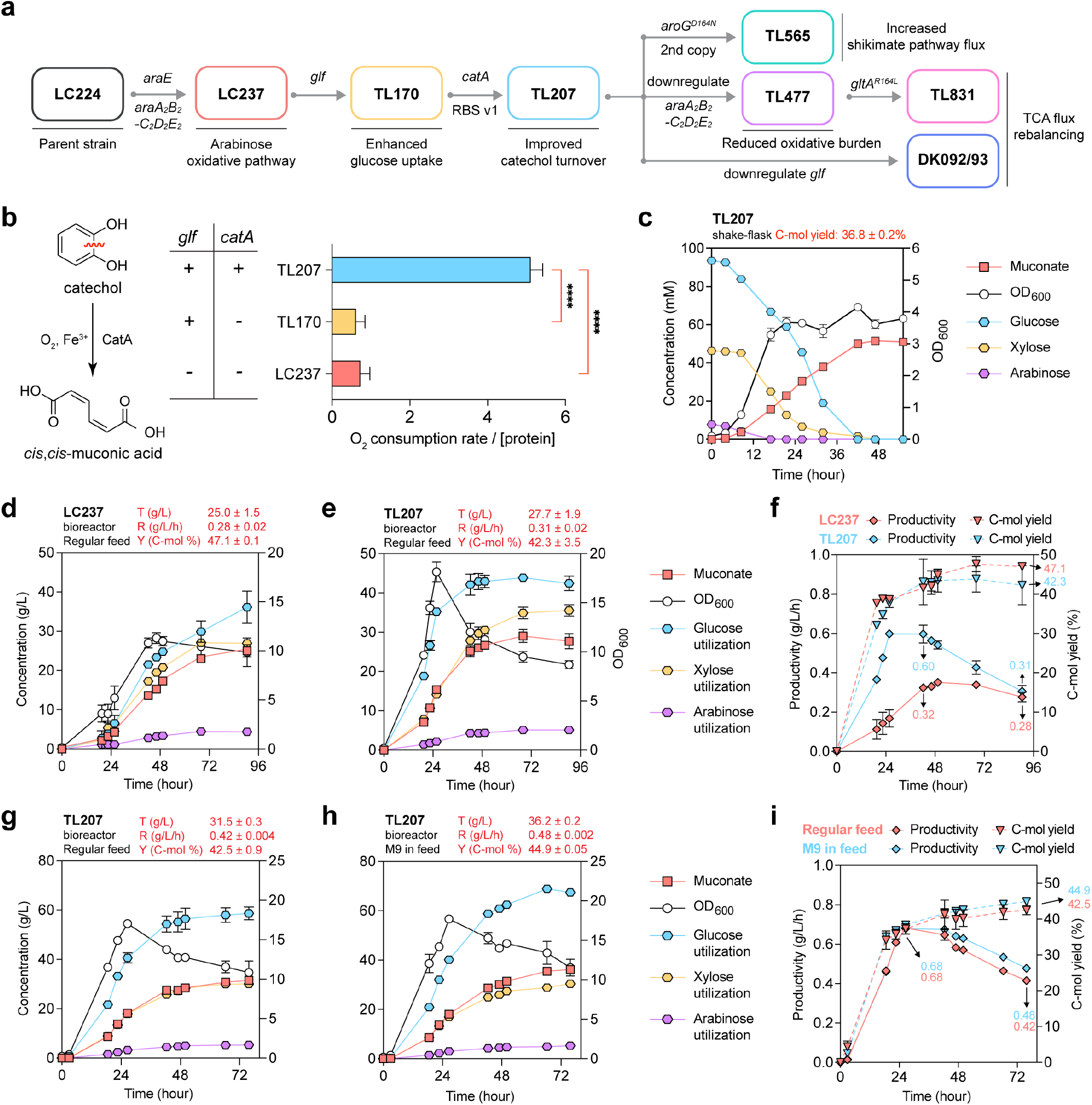
Engineering strategies to relieve glucose uptake and catechol ring-opening bottlenecks in the oxidative pathway strain. **a** Stepwise strain engineering strategies for arabinose-oxidative pathway optimization. Genetic modifications (above arrows) and their functional impacts (below or above strain names) are shown for each derivative strain starting from the parent strain LC224. Details of engineering after TL207 are shown in **Extended Data Figure 1** and **Extended Data Figure 2. b** Oxygen consumption assays using catechol as the substrate, comparing LC237, TL170, and TL207. Chemical reaction (top) and measured rates (bottom) are shown. Introduction of *glf* and overexpression of *catA1* on engineered strains are indicated (right). Statistical significance was determined by ordinary one-way ANOVA followed by Dunnett’s multiple comparisons test (****, p-value < 0.0001). **c** Shake-flask experiment profiles showing the bacterial growth, residual sugar concentrations, and muconate production of TL207 on M9 medium supplemented with 25 g/L mock hydrolysate (see **Fig. S9** for comparative profiles with LC237 and TL170). **d**-**e** Fed-batch bioreactor evaluation of LC237 (**d**) and TL207 (**e**) in 0.5-L scale with ~ 96 h of cultivation. Bacterial growth (OD_600_), muconate production and total sugar utilization of LC237 and TL207 are shown. Additional bioreactor profiles, including residual sugars in bioreactors during the cultivations are provided in **Fig. S10. f** Comparison of muconate productivity and carbon yield of LC237 and TL207. **g**-**h** Fed-batch bioreactor evaluation of TL207 using the regular feed (**g**) and M9-supplemented feed (**h**) with ~72 h of cultivation. Bacterial growth (OD_600_), muconate production, and total sugar utilization are shown. Additional bioreactor profiles, including residual sugars in bioreactors during the cultivations are provided in **Fig. S11. i** Comparison of muconate productivity and carbon yield of TL207 with media optimization. Concentrations are reported in mM for flasks and g/L for bioreactors to accommodate low and high ranges of substrates and products, respectively. Titer: final muconate concentration (g/L), Rate: titer at the final time point/time (g/L/h), Yield (C-mol %): [(mM muconate × 6) / mM (glucose × 6 + mM xylose × 5 + mM arabinose × 5) × 100%] calculated at the final time point. Data represent the average of *n* = 3 biological replicates in **b**-**c**, and *n* = 2 biological replicates in **d**-**i.** Error bars correspond to standard deviation in **b**-**c**, and absolute error between duplicates in **e**-**i**. Numerical data are provided in a **Source Data File**.

Finally, evaluation in fed-batch bioreactors demonstrated the advantage of the TL207 strain compared to LC237. Specifically, TL207 outperformed LC237 in growth rate, sugar consumption rate, and overall muconate titer and productivity (**Fig. 3d-f**). TL207 achieved a muconate titer of 25 g/L within only 42 h, resulting in a productivity of 0.60 g/L/h, nearly double that of LC237 (0.32 g/L/h). Notably, while TL207 was faster, LC237 exhibited a higher muconate C-mol yield of 47.1%, compared to 42.3% in TL207, over the course of cultivation as shown in **Fig. 3f** (see **Fig. S10** for sugar profiles). Based on the overall increase in sugar utilization in TL207 (which leads to further dilution of the batch media components), we evaluated whether nutrient supplementation in the feed could improve the strain performance. For that purpose, we tested our regular feed (sugars dissolved in deionized H_2_O) as well as feed dissolved in M9 medium. These cultivations (**Fig. 3g-h**) were conducted for 72 h rather than 96 h, as sugar utilization in TL207 ceased after 72 h (**Fig. 3e**). Supplementation with M9 medium enhanced the glucose consumption rate, leading to higher titer (36.2 g/L), rate (0.48 g/L/h), and C-mol yield (44.9 %) compared to regular feed (**Fig. 3g-i**), suggesting that nutrient limitation in TL207 may have occurred. Therefore, all subsequent bioreactor experiments were conducted using M9-supplemented feeds.

Despite the improved muconate production in strain TL207, the accumulation of metabolic intermediates such as pyruvate, lactate, and acetate was observed during the late-exponential growth phase (**Extended Data Figure 1, Fig. S12**). This accumulation typically occurs when the carbon flux toward pyruvate exceeds the processing capacity of the TCA cycle (*54, 60, 61*). Efforts to mitigate this overflow by attenuating carbon flux into the TCA cycle through feedback inhibition-resistant citrate synthase (*gltA*^*R164L*^) (*50*) (TL477 and TL831) or downregulating glucose uptake (DK092 and DK093) resulted in only modest reductions in accumulation and, in some cases, compromised overall glucose utilization rates (**Fig. 3a, Extended Data Figure 1**, and **Figs. S12**-S**14**; see **Supplementary Note 2** for a detailed discussion).

Lastly, to increase carbon flux into the shikimate pathway, we introduced an additional copy of a feedback-resistant DAHP synthase (AroG^D146N^) gene (*49*) into the oxidative strain TL207, generating strain TL565. This strategy was motivated by previous results showing that additional expression of *aroG*^*D146N*^ enhances precursor supply toward aromatic pathway-derived products (*14*). In shake-flask experiments using mock hydrolysate, TL565 showed similar growth, sugar consumption, and muconate production compared to TL207 (**Extended Data Figure 2**). In fed-batch bioreactor cultivation, however, TL565 exhibited faster growth and enhanced initial muconate production, suggesting that the additional AroG^D146N^ increased carbon flux toward the shikimate pathway. Specifically, TL565 reached a muconate titer of 40.9 g/L within 43 h, corresponding to a maximum productivity of 1.02 g/L/h, whereas TL207 produced only 32.6 g/L with a maximum productivity of 0.86 g/L/h during the same period (**Extended Data Figure 2**). However, despite its accelerated growth, TL565 also directed more carbon towards microbial biomass than TL207 (**Fig. S15**), leading to a substantially lower C-mol yield (41.6% versus 49.2% for TL207). We speculate that the enhanced consumption of PEP and E4P by the additional AroG^D146N^ relieved a potential upstream flux bottleneck in the ED pathway. However, this shift in metabolic flux may have disrupted the optimal balance between product formation and cellular growth, directing more carbon toward the TCA cycle for biomass generation rather than muconate production, thereby reducing the muconate yield. Given that the product yield is a primary metric for industrial bioprocess development when productivities are comparable, we selected TL207 as the optimal strain among the arabinose-oxidative strains.

Taken together, our sequential efforts to overcome the metabolic bottlenecks of the arabinose-oxidative pathway either by engineering overflow metabolism or by reinforcing precursor supply did not yield substantial improvements in overall process efficiency. These findings suggest that the arabinose-oxidative pathway may have approached its practical ceiling. Therefore, further improvements in muconate production would likely require a shift toward alternative strategies, such as the arabinose-isomerase pathway or more sophisticated bioprocess optimization (*vide infra*).

### Translating engineering strategies from the arabinose-oxidative pathway to the arabinose-isomerase pathway strains

Having established design strategies for the arabinose-oxidative pathway, we next applied them to the arabinose-isomerase strain LC357 (**Fig. 4a** and **Extended Data Table 1**). Unlike the oxidative strain LC237, LC357 exhibited a prolonged lag phase (**Figs. S1**-**S2**). This difference may stem from the metabolic convergence point of the isomerase pathway; since both arabinose and xylose metabolism converge on the non-oxidative pentose-phosphate pathway, excessive flux can trigger sugar-phosphate stress (*38, 62*). In this isomerase background, such stress likely impairs sugar uptake, ultimately hindering initial growth.

**Figure 4.**
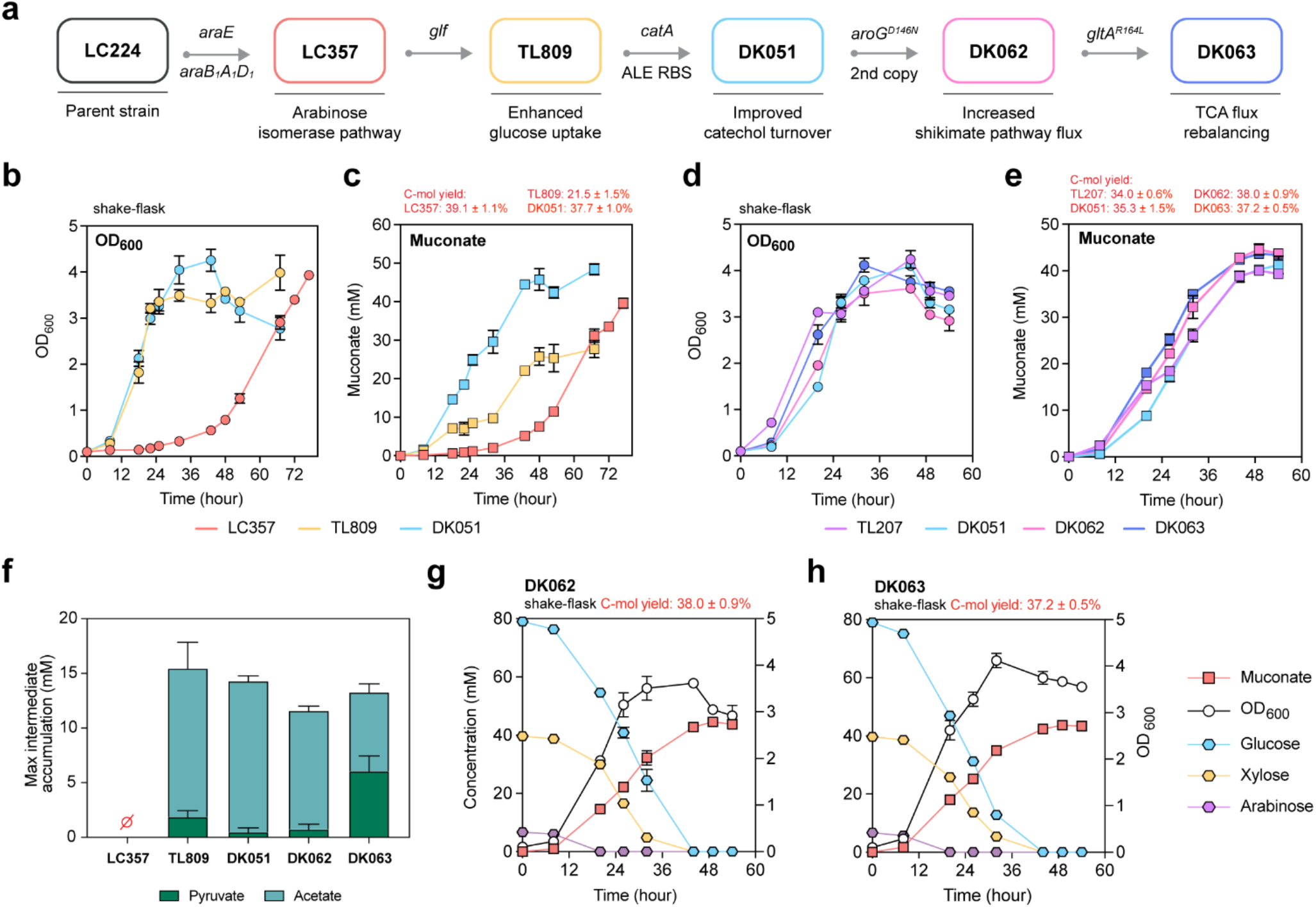
Engineering strategies to improve performance in the arabinose-isomerase strain. **a** Stepwise strain engineering strategies for arabinose-isomerase pathway optimization. Genetic modifications (above arrows) and their functional impacts (below strain names) are shown for each derivative strain starting from the parent strain LC224. **b-c** Growth (**b**) and muconate production (**c**) in arabinose-isomerase strains LC357, TL809, and DK051 with applied engineering strategies. Application of successive engineering strategies led to gradual improvements in both growth and muconate production (see **Fig. S17** for additional profiles). **d-e** Growth (**d**) and muconate production (**e**) in arabinose-isomerase strains DK051, DK062, and DK063 compared with the engineered arabinose-oxidative strain TL207. Introduction of stabilizing strategies from oxidative pathway engineering (TL207) into the isomerase background enabled DK062 and DK063 to achieve similar growth and muconate production performance (see **Fig. S18** for additional profiles). **f** Maximum net accumulation of metabolic intermediates observed in strains LC357, TL809, DK051, DK062, and DK063 during shake-flask experiments in the 24-48 h interval. Lactate was not detected. LC357 showed no accumulation. **g-h** Shake-flask experiment profiles showing the bacterial growth, residual sugar concentrations, and muconate production of DK062 (**g**) and DK063 (**h**) on M9 medium supplemented with 25 g/L mock hydrolysate. For **d-h**, see **Fig. S18** for detailed profiles. Maximum intermediate accumulation was calculated as the sum of pyruvate, lactate, and acetate concentrations (mM). C-mol yield was calculated as [(mM muconate × 6) / mM (glucose × 6 + mM xylose × 5 + mM arabinose × 5) × 100%]. Data represent the average of *n* = 3 biological replicates. Error bars correspond to standard deviation. Numerical data are provided in a **Source Data File**.

Therefore, to accelerate glucose uptake, we first introduced a high-expression *glf* cassette (RBSv1, 41-fold TIR) into LC357, yielding strain TL809. The introduction of the *glf* cassette facilitated the consumption of all three sugars during plate cultivation compared to LC357, resulting in improved growth in mock hydrolysate (**Fig. S16**). Similarly, in shake-flask experiments, TL809 exhibited enhanced sugar consumption (**Fig. S17**), leading to faster growth than LC357 (**Fig. 4b**).

However, while TL809 showed superior growth and sugar consumption profiles, its final muconate titer was lower than that of LC357 (**Fig. 4c**) This was reflected in a substantial reduction in the C-mol muconate yield, which fell from 39.1% in LC357 to 21.5% in TL809 (**Fig. 4c**). Furthermore, TL809 cultures showed a brown coloration during shake-flask experiments and confirmed catechol accumulation from HPLC analysis (**Fig. S17**). These results demonstrate that while *glf* introduction effectively resolves the growth lag by accelerating sugar uptake, a downstream bottleneck at the catechol 1,2-dioxygenase step limits high yield muconate production in the isomerase background, mirroring the phenotype of oxidative pathway strains.

To resolve this, we replaced the native *catA* RBS with a stronger ALE-derived variant (*59*), generating strain DK051. This modification eliminated catechol accumulation, accelerated sugar uptake, and increased muconate yield and growth as shown in **Fig. 4b-c**. Additionally, to further drive flux through the shikimate pathway, we introduced an additional copy of the gene encoding AroG^D146N^ (*49*) as we had in the oxidative strain. This modification resulted in strain DK062, which achieved the highest C-mol muconate yield (38% within 54 h) among all isomerase strains (**Fig. 4d-e**). Finally, we introduced the *gltA*^R164L^ mutation to generate strain DK063, aiming to minimize overflow metabolism as previously explored in the oxidative lineage. In this background, acetate accumulation was reduced relative to DK062, but pyruvate accumulation was greater, resulting in slightly higher overall intermediate accumulation in DK063 (**Fig. 4f**).

To evaluate our original hypothesis regarding the rate and yield tradeoffs between the two assimilation pathways, we compared these optimized isomerase pathway strains, DK062 and DK063, to the top-performing oxidative pathway strain, TL207. In shake-flask experiments, DK062 and DK063 displayed higher final muconate titers and C-mol yields, whereas TL207 exhibited the fastest growth kinetics (**Fig. 5a-b, Fig. S18**). Specifically, while TL207 reached 39.3 mM titer with 34% C-mol yield within 54 h, DK062 and DK063 reached 43.7 mM (38% yield) and 43.4 mM (37% yield), respectively, in the same period. These results underscore that while the oxidative pathway offers faster growth rates, the isomerase pathway can achieve higher final yields once bottlenecks in transport, catechol accumulation, and precursor supply are addressed, thereby validating our original hypothesis.

**Figure 5.**
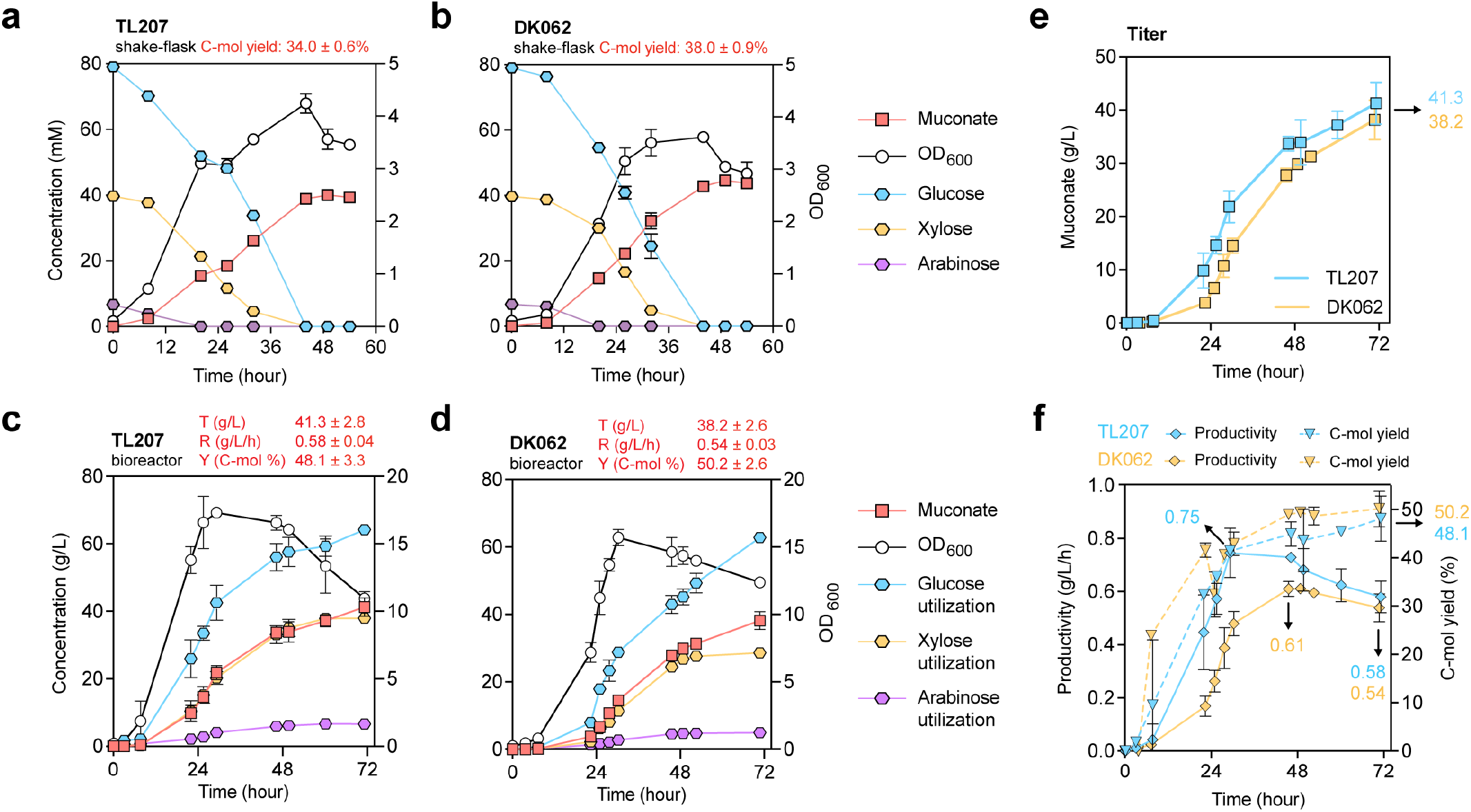
Evaluation of oxidative and isomerase strains in 0.5-L bioreactors in fed-batch mode. **a-b** Shake-flask experiment profiles showing the bacterial growth, residual sugar concentrations, and muconate production of TL207 (**a**) and DK062 (**b**) on M9 medium supplemented with 25 g/L mock hydrolysate (see **Fig. S18** for detailed profiles). **c**-**d** Evaluation of TL207 (**c**) and DK062 (**d**) in 0.5-L bioreactors in fed-batch model. Cultivation profiles show bacterial growth (OD_600_), sugar consumption, and muconate production. Additional bioreactor profiles, including residual sugars in bioreactors during the cultivations and catabolic intermediates are provided in **Fig. S19. e**-**f** Comparison of muconate titer (**e**), rate, and carbon yield (**f**) of TL207 and DK062 with their final titers, maximum and final productivities, and final C-mol yields presented. For comparative profiles with TL207, DK062, and DK063, see **Fig. S20.** Concentrations are reported in mM for flasks and g/L for bioreactors to accommodate low and high ranges of substrates and products, respectively. Titer: final muconate concentration (g/L), Rate: titer at the final time point/time (g/L/h), Yield (C-mol %): [(mM muconate × 6) / mM (glucose × 6 + mM xylose × 5 + mM arabinose × 5) × 100%] calculated at the final time point. Data represent the average of *n* = 3 biological replicates in **a**-**b**, and *n* = 2 biological replicates in **c**-**f**. Error bars correspond to standard deviation in **a**-**b**, and absolute error between duplicates in **c**-**f**. Numerical data are provided in a **Source Data File**.

To determine whether the distinct performance differences observed in shake-flask experiments translate to bioreactors, we evaluated the oxidative strain TL207 and the isomerase strains DK062 and DK063 in 0.5-L fed-batch bioreactors on mock hydrolysate (**Fig. 5c-d, Fig. S19**). The performance metrics for all three strains are summarized in **Fig. 5e-f** and **Fig. S20**. TL207 achieved a final muconate titer of 41.3 g/L with an overall productivity of 0.58 g/L/h (maximum productivity of 0.75 g/L/h) and a C-mol yield of 48.1%. In comparison, the isomerase strains DK062 and DK063 reached slightly lower final titers of 38.2 g/L and 39.2 g/L, respectively, but achieved higher C-mol yields of 50.2% and 49.6%. While their maximum productivities (0.61 g/L/h for DK062 and 0.65 g/L/h for DK063) were lower than that of TL207, they maintained competitive final productivities of 0.54 g/L/h and 0.55 g/L/h.

As shown in **Fig. S19**, TL207 exhibited faster sugar consumption than the isomerase strains. While DK062 and DK063 reached similar levels of glucose utilization by the end of the fermentation, they utilized less total pentose sugar than TL207, suggesting that the oxidative pathway facilitates more efficient co-utilization of xylose and arabinose. Notably, acetate accumulation was observed in all three strains, with DK063 showing levels nearly identical to that of DK062 (**Fig. S19**). Given the similar muconate TRY and catabolic intermediate profiles (**Figs. S19**-**20**), these results suggest that the introduction of *gltA*^R164L^ into the isomerase strain exhibited only a minor effect on muconate production and the mitigation of intermediate accumulation, consistent with the observations in the oxidative strain.

These findings indicate that the arabinose-oxidative pathway enabled rapid carbon flux directly into the TCA cycle, favoring cell biomass accumulation and high production rates. In contrast, routing arabinose through the isomerase pathway into the non-oxidative PP pathway improved carbon efficiency with respect to muconate production. TL207 and DK062 substantially outperformed the parental strain LC224 (**Fig. 2e, Fig. 5e-f**). Thus, with the addition of arabinose catabolic pathways and the Glf glucose transporter, as well as increased expression catechol dioxygenase, the engineered strains achieved on average a 44% increase in titer, an 87% increase in productivity, and a 31% increase in yield relative to LC224. Together, these results illustrate a fundamental tradeoff between rate and yield that emerged from alternative arabinose assimilation pathways based on where they integrate into central carbon metabolism, providing a strategic framework for pathway selection.

### Techno-economic analysis and life cycle assessment on engineered strains

Based on the performance improvements enabled by arabinose utilization and pathway engineering, we evaluated the economic and environmental impacts of our improved strains through process modeling, TEA, and LCA (**Fig. 6**). Detailed methods for the TEA and LCA are provided in the Methods section, **Supplementary Note 4**, and **Fig. S21**. Despite arabinose being a relatively minor sugar in lignocellulosic hydrolysate, the engineered strain DK062 achieved the lowest estimated minimum selling price (MSP) for adipic acid at $2.74/kg, representing a 26% reduction compared to the parental strain LC224, which lacks arabinose utilization capability. The results in **Fig. 6b** demonstrate that integrating arabinose utilization with pathway engineering can bring the MSP of bio-based adipic acid closer to the $1.00-2.50/kg market range of its conventionally sourced fossil counterpart (*63*). As shown in **Fig. 6c**, the two largest drivers of the total production cost were feedstock costs (50%) and capital charges (30%). Since these two cost components are directly impacted by bioconversion yield and rate, respectively, the overall reduction in MSP for DK062 relative to the parental strain is driven by concurrent improvements in both yield and rate. Furthermore, when compared to fast-growing TL207, TL207 exhibited a higher productivity (0.58 g/L/h) than DK062 (0.54 g/L/h), but it yielded a higher MSP at $2.81/kg. This highlights that under a feedstock-dominated cost structure, even a slightly lower C-mol yield (48.1% for TL207 versus 50.2% for DK062) outweighs marginal gains in productivity. Additional economic results are showcased in **Fig. S22**. Notably, the MSPs shown in **Fig. 6b-c** correspond to the TRY metrics at the final timepoint of each cultivation. If the analysis is evaluated under maximum rate conditions, strain DK063 would yield the lowest MSP at $2.63/kg with 31.7 g/L titer,51.1 % C-mol yield, and 0.65 g/L/h productivity (**Fig. S23**). Although this operating point features a lower final titer than the final timepoint (39.2 g/L), the reduced fermentation time minimizes capital charges per batch. This suggests that prioritizing higher initial rates can provide a faster biorefinery turnover.

**Figure 6.**
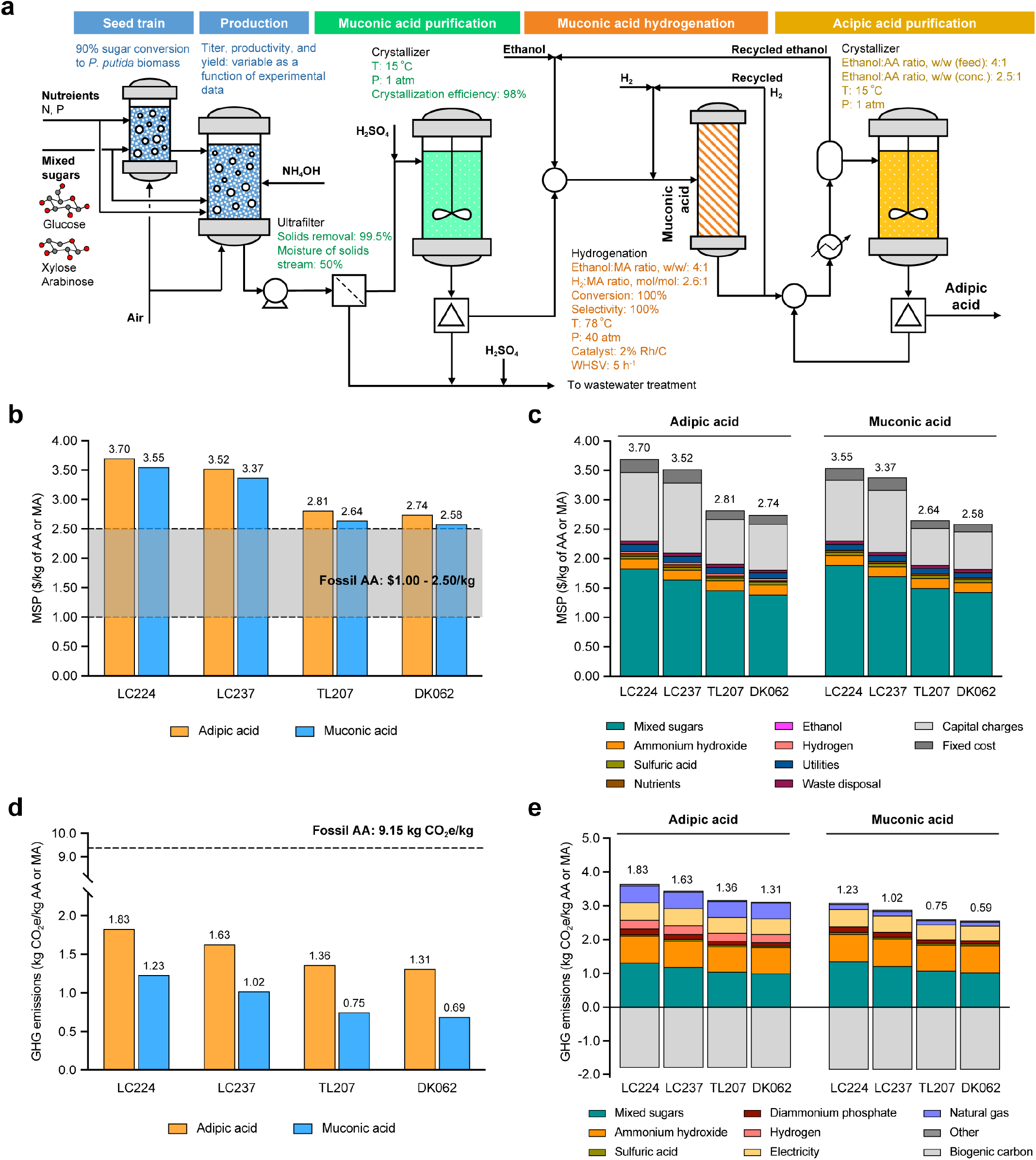
Process model, techno-economic analysis, and life cycle assessment for muconic acid (MA) and adipic acid (AA) production from lignocellulosic sugars. **a** Process flow diagram of MA and AA production from mixed sugars. The process includes a seed train, production in *P. putida*, MA purification, hydrogenation, and AA purification. Key process conditions and simulation parameters are highlighted as color-coded. Mixed sugars contain glucose, xylose, and arabinose derived from corn stover (see **Fig. S21** for detailed process flow diagram). **b** Minimum selling prices (MSPs) for MA (blue) and AA (orange) produced from mixed sugars using bioprocess metrics from a benchmark study by Ling *et al*. (2022) (LC224) (*27*), and those determined in this study (LC237, TL207, and DK062). Total MSP values are compared against the fossil-derived AA price range ($1.00-$2.50/kg). Bioprocess metrics are based on the titer, rate, and yield achieved at the final timepoint of the bioreactor cultivations. **c** Detailed MSP cost breakdown into key contributors, including mixed sugars, ammonium hydroxide, sulfuric acid, nutrients, ethanol, hydrogen, utilities, waste disposal, capital charges, and fixed cost, in MA and AA production. The sum of costs from all contributors is corresponding to the final MSP value in **b. d** Greenhouse gas (GHG) emissions for MA and AA for the strains described in **b.** Total GHG emissions are compared against the fossil-derived AA benchmark (9.15 kg CO_2_e/kg). **e** Detailed GHG emissions breakdown into key contributors, including mixed sugars, ammonium hydroxide, sulfuric acid, diammonium phosphate, hydrogen, electricity, natural gas, biogenic carbon, and others in MA and AA production. Negative values represent the credit from biogenic carbon. The sum of GHG emissions from all contributors is corresponding to the final GHG emissions value in **d**. MSPs and GHG emissions calculated from maximum rate are presented in **Fig. S23** for comparison. Numerical data are provided in a **Source Data File**.

From an LCA standpoint, strain DK062 enabled an 86% reduction in greenhouse gas (GHG) emissions compared to fossil-derived adipic acid (**Fig. 6d**), emphasizing the environmental benefits of this biomanufacturing approach. Compared to the parent strain LC224, the engineered strains TL207 and DK062 resulted in an average 27% reduction in GHG emissions, confirming that expanding the substrate to arabinose reduces this environmental impact. Consistent with the TEA trends, evaluating the process’s maximum rate conditions showed that strain DK063 would achieve the lowest GHG emission at 1.28 kg CO_2_e/kg (**Fig. S23**), again highlighting the potential benefits of a strategic bioprocess termination once peak productivity is reached. As shown in **Fig. 6e**, mixed sugar feedstock (32-36%) and ammonium hydroxide (22-25%) are the two primary GHG emission contributors. Specially for the latter, its substantial contributions of ammonium hydroxide are due to the energy-intensive Haber-Bosch process required for ammonia production (*64*). If the current grid electricity is replaced by low-carbon sources (e.g., nuclear, wind, or solar), the overall GHG emissions could be further reduced by 29-35%, which would lower the GHG emissions of DK062 from 1.31 to 0.85 kg CO_2_e/kg (**Figs. S24**). While all results discussed up to this point have been generated with the R&D GREET version 2025 (*65*), we also conducted LCA using Brightway version 3.6.6 (*66*) to assess GHG emissions among strains (main results shown in **Figs. S25**-**S26**). In addition, our engineered strains reduced fossil resource use by 47% and cumulative energy demand by 37% on average compared to the fossil-derived adipic acid production, further emphasizing the environmental benefits (**Fig. S26**).

## Discussion

Efficient valorization of lignocellulosic feedstocks requires a strategic understanding of how different carbon assimilation routes influence host physiology and product formation. In this work, we engineered *P. putida* into a robust glucose, xylose, and arabinose co-utilizing strain, achieving muconate production at industrially relevant metrics. While prior efforts in *E. coli (42), Bacillus subtilis* (*43*), and *Saccharomyces cerevisiae* (*41*) primarily focused on isomerase pathways for pentose co-utilization, our findings reveal that the oxidative and isomerase pathways for L-arabinose are not merely alternative entry points for carbon, but rather, they enable fundamental metabolic tradeoffs between rate and yield for products derived from central carbon metabolism that precedes the TCA cycle. The oxidative pathway (TL207) routs arabinose into the TCA cycle, providing the energy (ATP/NADH) required for biomass production and high productivity (0.58 g/L/h), whereas the isomerase pathway (DK062) directs arabinose into the non-oxidative PP pathway to enhance precursor supply for the shikimate pathway, achieving a C-mol muconate yield of 50%.

Interestingly, these flux-shaping effects were triggered by arabinose present at relatively low concentrations compared to other sugars. The early depletion of arabinose relative to other carbon sources in bioreactors suggests that its assimilation may serve as a primary metabolic driver, establishing the high energy- or precursor-rich environments for muconate production before the other carbon sources are fully utilized. We similarly predict that these pathways will have different trade-offs for molecules made from the TCA cycle, with further differences for products derived from TCA cycle intermediates upstream and downstream of α-ketoglutarate, where the oxidative arabinose pathway enters.

A key insight from this study is the context-dependent impact of introducing an additional copy of *aroG*^*D146N*^ across two distinct arabinose pathways. In the isomerase background, reinforcing the shikimate pathway with an additional copy of feedback-resistant DAHP synthase effectively increased muconate production. In contrast, the same strategy in the oxidative background disrupted the optimal metabolic balance and failed to improve overall carbon efficiency. We posit that this divergence stems from how the two pathways interface with central carbon metabolism; while the isomerase pathway channels arabinose through the PP pathway, where an abundant supply of PEP and E4P naturally supports enhanced flux through shikimate pathway, the oxidative pathway directs arabinose toward the TCA cycle. In this context, limited supply of PEP and E4P with enhanced demand imposed by additional AroG^D146N^ may have triggered an increased overall carbon uptake and flux. However, this increased flux was not exclusively directed toward muconate production but was largely distributed toward biomass generation, thereby compromising the muconate yield. It should also be noted that mutations in *aroG*^*D146N*^ can improve growth at the expense of muconate production (*27*) and having two copies could prevent such mutations from being advantageous and overtaking a population in a process at scale. Our findings suggest that the integration of heterologous pathways is not always a “plug-and-play” process but must be contextually optimized relative to host metabolism and flux distribution (**Extended Data Fig. 3**).

Although we have overcome productivity barriers that have long hindered muconate production, the accumulation of acetate and pyruvate suggests a fundamental metabolic bottleneck occurring when catabolism becomes saturated due to accelerated sugar uptake (*54*). ALE of a strain with the *glf* integrated could provide a strategic solution to balance sugar influx with the internal metabolic capacity. Furthermore, we acknowledge that the transition to real corn stover hydrolysate and industrial scales typically reduces strain performance (*67*). While our engineered strains performed well in mock hydrolysate at the bench scale, future efforts will focus on understanding and engineering molecular mechanisms to enhance microbial performance for the conversion of true lignocellulosic streams at larger scales. From a process development point of view, implementing dynamic feeding control via real-time metabolite monitoring to prevent intermediate accumulations and developing *in situ* product recovery will be essential to mitigate product inhibition and maximize TRY performance at scale, which will be pursued in future work (*68, 69*). Overall, these results indicate that only modest improvements to the strain or bioprocess could bring the MSP into the cost range of fossil-based adipic acid (**Fig. 6b**), paving the way for industrially viable production of muconic acid and adipic acid.

**Extended Data Table 1.**
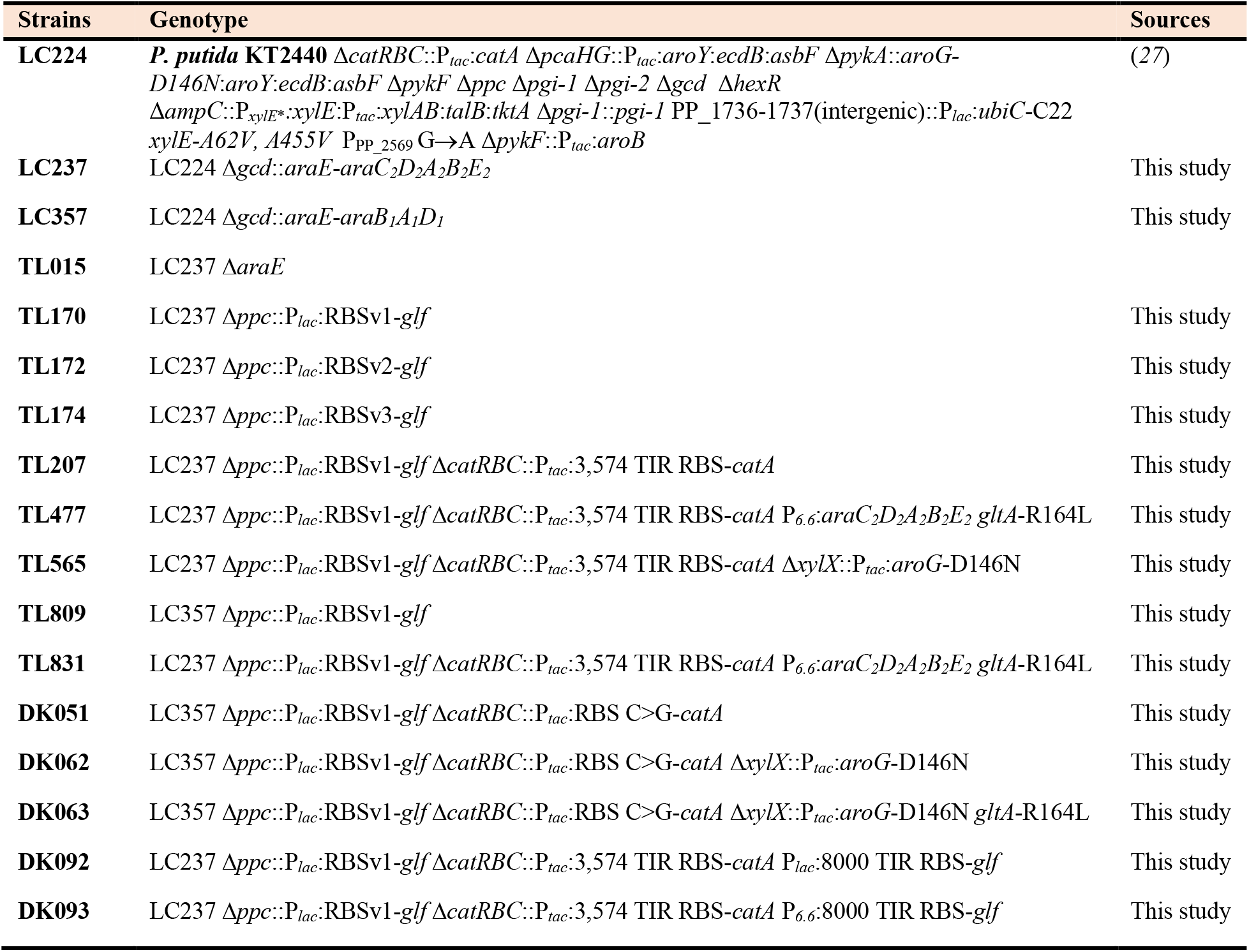
Genotypes of strains used in this study.

**Extended Data Figure 1.**
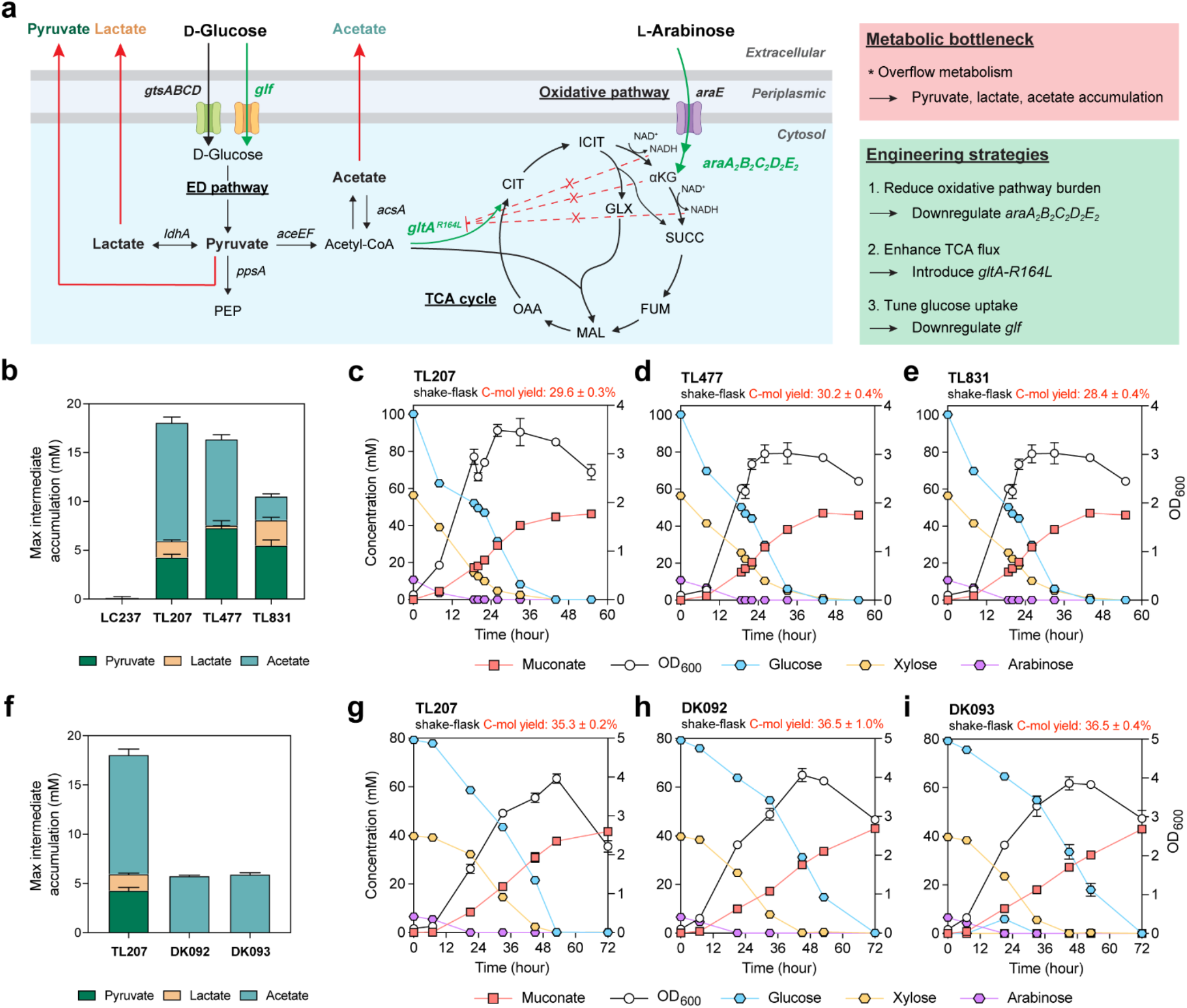
Engineering strategies to alleviate overflow metabolism in TL207. **a** Schematic of glucose and arabinose utilization pathway highlighting the accumulation of overflow metabolites (pyruvate, lactate, acetate) and the corresponding engineering strategies. Three approaches include: (i) reducing metabolic burden from the arabinose-oxidative operon (downregulation of *araA*_*2*_*B*_*2*_*C*_*2*_*D*_*2*_*E*_*2*_), (ii) enhancing TCA cycle flux by introducing a feedback-resistant citrate synthase (*gltA*^R164L^), and (iii) tuning glucose uptake by downregulating *glf*. **b** Maximum net accumulation of metabolic intermediates observed in strains TL207, TL477, and TL831 during shake-flask experiments in the 24-48 h interval. **c-e** Shake-flask experiment profiles showing the bacterial growth, residual sugar concentrations, and muconate production of TL207 (**c**), TL477 (**d**), and TL831 (**e**), on M9 medium supplemented with 25 g/L mock hydrolysate (see **Fig. S12** for additional profiles). **f** Maximum net accumulation of metabolic intermediates observed in strains TL207, DK092, and DK093 during shake-flask experiments in the 24-48 h interval. **g-i** Shake-flask experiment profiles showing the bacterial growth, residual sugar concentrations, and muconate production of TL207 (**g**), DK092 (**h**), and DK093 (**i**), on M9 medium supplemented with 25 g/L mock hydrolysate (see **Fig. S14** for additional profiles). Maximum intermediate accumulation was calculated as the sum of pyruvate, lactate, and acetate concentrations (mM). C-mol yield was calculated as [(mM muconate × 6) / mM (glucose × 6 + mM xylose × 5 + mM arabinose × 5) × 100%]. Data represent the average of *n* = 3 biological replicates in **b**-**i**. Error bars correspond to standard deviation. Numerical data are provided in a **Source Data File**.

**Extended Data Figure 2.**
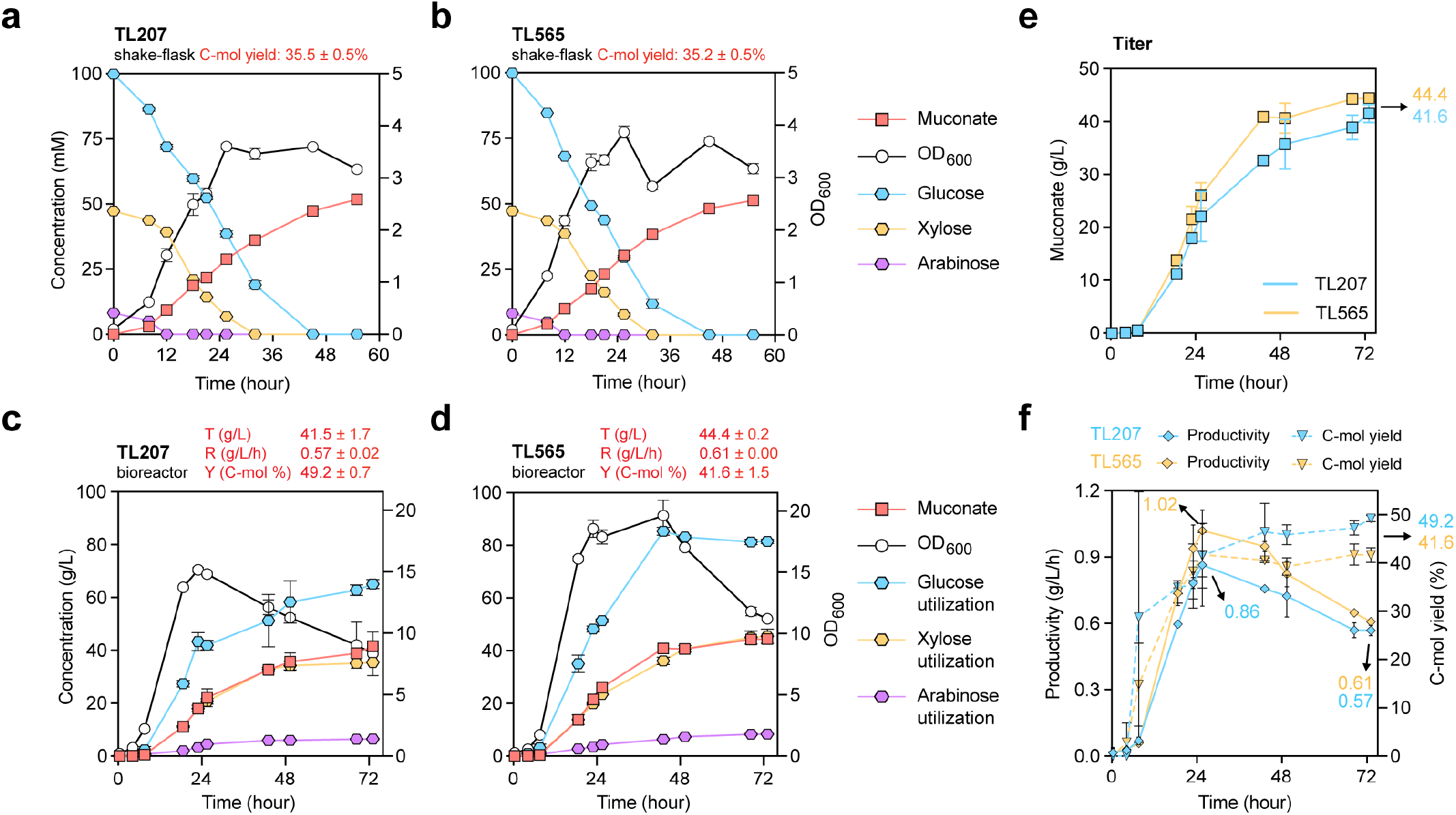
Engineering effect of additional *aroG*^*D164N*^ integration into TL207. **a**-**b** Shake-flask experiment profiles showing the bacterial growth, residual sugar concentrations, and muconate production of TL207 (**a**) and TL565 (**b**) on M9 medium supplemented with 25 g/L mock hydrolysate. **c**-**d** Evaluation of TL207 (**c**) and TL565 (**d**) in 0.5-L bioreactors in fed-batch model. Cultivation profiles show bacterial growth (OD_600_), sugar consumption, and muconate production. Additional bioreactor profiles, including residual sugars in bioreactors during the cultivations and catabolic intermediates are provided in **Fig. S15. e**-**f** Comparison of muconate titer (**e**), rate, and carbon yield (**f**) of TL207 and TL565 with their final titers, maximum and final productivities, and final C-mol yields presented. Concentrations are reported in mM for flasks and g/L for bioreactors to accommodate low and high ranges of substrates and products, respectively. Titer: final muconate concentration (g/L), Rate: titer at the final time point/time (g/L/h), Yield (C-mol %): [(mM muconate × 6) / mM (glucose × 6 + mM xylose × 5 + mM arabinose × 5) × 100%] calculated at the final time point. Data represent the average of *n* = 3 biological replicates in **a**-**b**, and *n* = 2 biological replicates in **c**-**f**. Error bars correspond to standard deviation in **a**-**b**, and absolute error between duplicates in **c**-**f**. Numerical data are provided in a **Source Data File**.

**Extended Data Figure 3.**
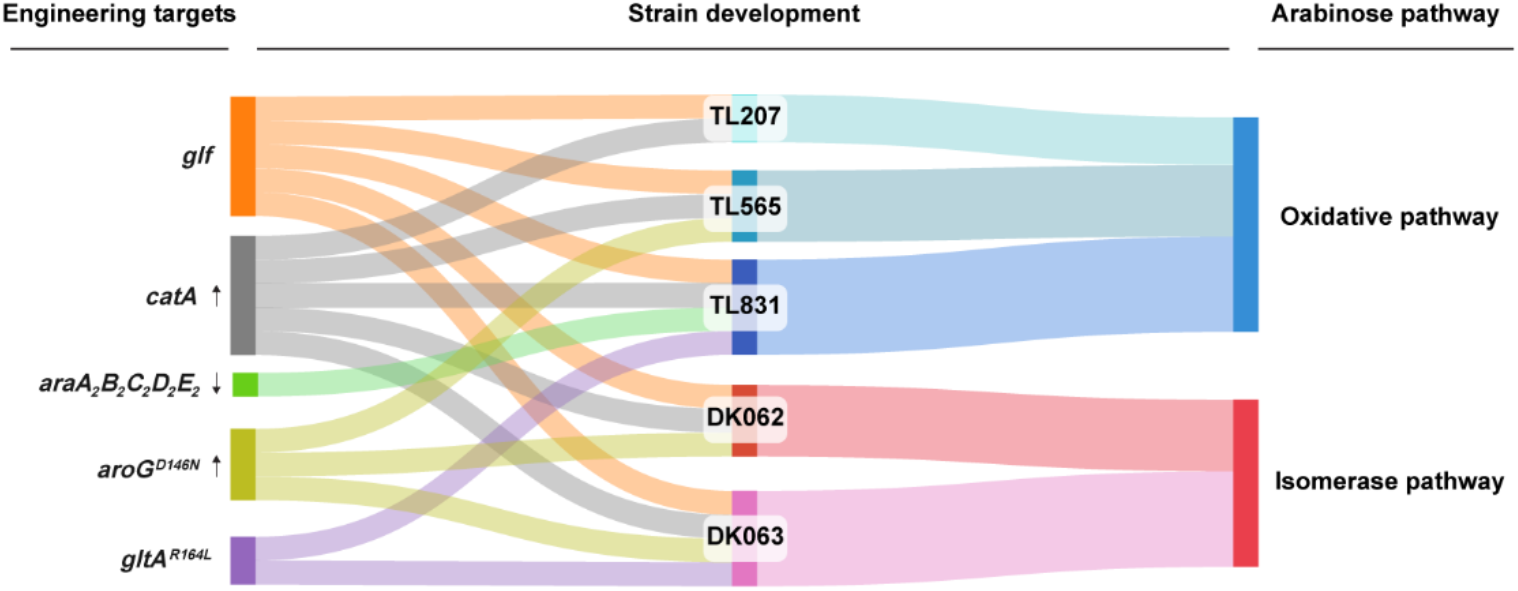
Schematic overview of genetic engineering strategies and strain lineages across metabolic pathways. Sankey diagram illustrating the relationship between specific genetic engineering targets, engineered strains, and their corresponding arabinose pathways. Arabinose-oxidative pathway derived strains (TL207, TL565, TL831) and isomerase pathway derived strains (DK062, and DK063) are shown. The connecting bands indicate the inclusion of specific engineering targets with each lineage, demonstrating the shared engineering features between the different approaches. The diagram was designed using SanKeyMATIC tool.

## Materials and methods

### Plasmid construction

All plasmids used to perform gene replacements in this study were constructed in the backbone plasmid pK18mobSacB (Genbank FJ437239, ATCC 87097) (*70*), or its derivatives, pK18msB (GenBank OK423783, Addgene Plasmid 177839) (*27*), and pK18msBI (pK18msB with *lacI*) which have the pMB1 replicon originated from *E. coli* (*70*), and do not replicate in *P. putida* (*71*). They also contain a kanamycin resistance gene as a selectable marker and a *sacB* gene encoding the levansucrase that converts sucrose into levans, which is lethal to many gram-negative bacteria including *P. putida*, as a counter-selectable marker. Each plasmid also has two fragments homologous to the upstream and downstream sequences flanking the target site on the genome to enable recombination of the plasmids into and out of the genome.

For plasmid cloning, fragments and backbone vectors were amplified from genomic DNA of *E. coli* or *P. putida* by polymerase chain reactions (PCR) using Q5^®^ High-Fidelity 2X Master Mix (New England Biolabs). Some fragments were also synthesized from Twist Bioscience. The PCR products of vector backbone, and fragments derived from plasmids with same antibiotic resistance gene, were subsequent digested by adding DpnI (New England Biolabs) enzyme into the PCR products, digestion to remove the cell-derived plasmid template. We typically add 1 μL DpnI into 50 μL Q5^®^ PCR products without adding digesting buffer, assuming that DpnI remains active in the Q5^®^ PCR system. The PCR products with DpnI were subsequent incubated at 37°C for 30 min and then purified using the DNA purification kit (Thermo Scientific™ K0831). Subsequently, NEBuilder HiFi Assembly Master Mix (New England Biolabs) were used to assemble the purified fragments into the backbone vector. The assembled products were then transformed into chemically competent NEB^®^ 5-alpha F’*I*^*q*^ *E. coli* (New England Biolabs) cells according to the manufacturer’s instructions. The tight control of expression by *lacI*^*q*^ allows for the cloning of potentially toxic genes. The transformed cells were plated on Lysogeny Broth (LB) agar (Lennox) plates supplemented with 50 μg/mL kanamycin (Sigma-Aldrich) and incubated overnight at 37°C to allow for the development of single colonies. Colony PCR was conducted to pre-screen single colonies for the desired band size using agarose gel electrophoresis. Colonies with the desired band were inoculated into liquid LB (Miller’s) culture and incubated at 37°C with shaking at 225 rpm. Plasmids were then purified from the liquid culture and sent for Nanopore sequencing (Plasmidsaurus) to confirm the correct plasmid sequence. All plasmids used in this study are listed and described in detail in **Table S4**.

### Strain construction

Gene replacements, deletions, and insertions were accomplished by incorporating the relevant plasmids into the genome through homologous recombination. Specifically, plasmids were transformed into relevant *P. putida* strains via either electroporation or conjugation.

For electroporation, *P. putida* strains were inoculated from a glycerol stock into 14-mL round bottom Falcon^®^ tubes containing 5 mL LB medium and incubated overnight at 30 °C with shaking at 225 rpm. Electrocompetent cells were prepared by washing the cells with 300mM sucrose three times (*72*). Washed cells were mixed with 300-500 ng of plasmid and electroporated in 0.1 cm cuvette using a Gene Pulser Xcell (Bio-Rad) with the setting 1.6 kV, 25 μF, 200 ohms (*73*).

For conjugation, plasmids were transferred into to the recipient *P. putida* strains from donor *E. coli* WM6026 cells (*74*). *E. coli* WM6026 is an auxotrophic strain whose growth relies on exogenously supplemented diaminopimelic acid (DAP), and its *lacI*^*q*^ genotype ensures tight control of the gene expression on the plasmid, avoiding mutation in the plasmid during the conjugation process. Electrocompetent *E. coli* WM6026 cells were prepared by washing with ice-cold 10% glycerol and H_2_O, then stored at −80 °C. Upon electroporation, frozen cells in Eppendorf tubes were thawed on ice for 10 minutes. Subsequently, 300-500 ng of the relevant plasmid was added and electroporated using parameters similar to those for *P. putida* (*vide supra*). After electroporation, 950 μL of SOC outgrowth medium (New England Biolabs) was added. The cell cultures in Eppendorf tubes were taped to the incubator surface and incubated at 37 °C with shaking at 225 rpm for 1 hour to recover. After recovery, cells were plated on LB agar plates supplemented with 50 μg/mL kanamycin and 300 μM DAP and incubated overnight at 37 °C. The plasmid-containing donor *E. coli* WM6026 strain was then inoculated into LB medium supplemented with 50 μg/mL kanamycin and 300 μM DAP and cultivated in a shaking incubator at 37 °C, 225 rpm overnight; the recipient *P. putida* strain was inoculated into LB medium and cultivated in a shaking incubator overnight at 30 °C, 225 rpm. 1 mL of both donor and recipient cells from the overnight cultures were centrifuged at 5000 *g* for 2 min followed by a double wash in LB medium to remove any antibiotic. The resuspended pellets were then combined in a microcentrifuge tube, mixed, and centrifuged again. The mixed cell pellet was then resuspended in 50 µL LB medium and plated onto a LB plate with kanamycin and DAP at low concentration: 10 μg/mL kanamycin to inhibit cell growth of untransformed *P. putida*, and 30 μM DAP to support *E. coli* WM6026 growth. The plate was incubated at 30 °C for 6-10 h to allow for conjugation, after which a scratch of cells was resuspended in 200 µL LB medium and plated onto a LB plate supplemented with 50 μg/mL kanamycin (to select for *P. putida* with kanamycin resistance gene) and no DAP (to select against the *E. coli* WM6026 donor). The plate was incubated at 30 °C overnight, and single colonies were re-streaked to a new LB plate with 50 μg/mL kanamycin to purify *P. putida* cells containing the kanamycin resistance gene and ensure no carryover of *E. coli* WM6026 donor cells.

As the plasmids in our study cannot replicate in *P. putida*, the kanamycin resistance gene must be integrated into the genome through homologous recombination to confer resistance. Following initial selection on LB plates supplemented with 50 μg/mL kanamycin, colonies were restreaked onto fresh kanamycin-supplemented LB plates to purify the integration strains. Subsequently, sucrose-sensitive SacB counterselection was performed by streaking the purified cells on YT (Yeast-Tryptone) agar plates (10 g/L tryptone, 5 g/L yeast extract, and 18 g/L Bacto agar) with 25 % (w/v) of sucrose (*73*). Diagnostic colony PCR using MyTaq™ HS Red Mix (Bioline) was used to identify correct mutants based on product size or presence. Once construction is confirmed through colony PCR, single colonies were re-streaked to a LB plate without kanamycin to get the isolates. All strains and construction details are listed and described in **Tables S2-S3**.

### Shake-flask and plate reader experiments

Seed cultures were inoculated from glycerol stocks into 14-mL round bottom Falcon^®^ tubes containing 5 mL LB medium and grown overnight at 30 °C with a shaking speed of 225 rpm. After overnight incubation, cultures were transferred into 125-mL baffled shake-flasks containing 10 mL LB medium to an initial OD_600_ of 0.2. The flasks with second seed cultures were then incubated at 30 °C and 225 rpm for approximately 4 h to reach an OD_600_ of ~2. Note that the final OD_600_ of different strains may vary due to differing growth lags. Following this growth phase, the cultures were pelleted by centrifugation at 4000 *g* and washed twice with M9 salts (6.78 g/L Na_2_HPO_4_, 3 g/L KH_2_PO_4_, 0.5 g/L NaCl, 1 g/L NH_4_Cl). For shake-flask experiments, washed cells were inoculated to 125-mL baffled shake-flask containing 25 mL modified M9 minimal medium (M9 salts plus 2 mM MgSO_4_, 100 µM CaCl_2_, 18 µM FeSO_4_) supplemented with glucose, xylose and arabinose, to an initial OD_600_ of 0.1. The molar ratio of glucose, xylose, and arabinose in mixed substrates was 2:1:0.17, reflecting the typical composition of corn stover hydrolysates (*6*). Samples were taken periodically throughout the cultivation to measure bacterial growth (OD_600_). Cells were pelleted and the supernatant was passed through a 0.2-µm filter into a glass HPLC vial fitted with a rubber septum for metabolite analysis. The pH values of flasks were monitored at each sampling time point using a mini pH meter (HORIBA LAQUAtwin pH-33), and when necessary, 1 N NaOH was used to adjust the pH to 7.

For plate reader experiments, initial growth measurements were characterized using Bioscreen C Pro analyzers (Growth Curves USA) with 300 µL cultures set to an initial OD600 of 0.1. Overnight strain cultivations and downstream washing procedures were conducted identically to those described for the shake-flask experiments. For each strain, cells were aliquoted in triplicate with un-inoculated wells included as controls. Shake-flask and plate reader data were plotted and analyzed using GraphPad Prism version 8.4.2. Carbon yields were calculated at the time point where maximum muconate concentration was detected.

### Seed train preparation prior bioreactor cultivations

To test the different strains in bioreactors, independent seed cultures for each bioreactor were prepared from glycerol stocks in 250 mL baffled flasks containing 50 mL of LB Miller. These seed cultures were incubated at 30 °C and 225 rpm for ~16h before pelleting the cell biomass through centrifugation at 5,000 *g* for 8 min. This pellet was then re-suspended in LB Miller to inoculate 250 mL baffled shake-flasks (containing 50 mL of LB Miller) at a target optical density at 600 nm (OD_600_) of 0.2 (measured with the NanoDrop One, Thermo Scientific). These seed cultures were incubated at 30 °C and 225 rpm for ~4 h before pelleting the cell biomass through centrifugation at 5,000 *g* for 8 min and resuspending it in 5 mL of M9 medium (13.56 g/L Na_2_HPO_4_, 6 g/L KH_2_PO_4_, 1 g/L NaCl, and 2.25 g/L (NH_4_)_2_SO_4_). Each seed was used to inoculate bioreactors to a target OD_600_ of 0.2.

### Bioreactor cultivations in fed-batch mode

The bioreactors (0.5-L, BioStat-Q Plus, Sartorius Stedium) contained an initial batch volume of 250 mL. The medium in the batch phase contained 13.56 g/L Na_2_HPO_4_, 6 g/L KH_2_PO_4_, 1 g/L NaCl, and 2.25 g/L (NH_4_)_2_SO_4_, 0.24 g/L MgSO_4_ x 7 H_2_O, 11 mg/L CaCl_2_ x 2 H_2_O, and 2.73 mg/L FeSO_4_ x 7 H_2_O as well as 25 g/L of sugars (mimicking sugar proportions in corn stover-derived hydrolysate: 16.8 g/L glucose, 7 g/L xylose, and 1.2 g/L of arabinose). Prior to inoculation, three drops of Antifoam 204 (Sigma) were added to each reactor. The bioreactors were maintained at 30°C and pH 7 (controlled by 4N H_2_SO_4_ and 8N NH_4_OH) and air was sparged at 300 mL/min. The initial agitation speed and dissolved oxygen (DO) of the batch phase was 350 rpm and 100%, respectively. Once DO dropped to 30%, an automatic agitation cascade was employed to maintain it at this level. After ~10 h of batch phase, the fed-batch phase was initiated. The feeding medium utilized to evaluate LC237, LC224, and TL207 consisted of 336 g/L of glucose, 140 g/L of xylose, 23 g/L of arabinose, and 4 mL/L of Antifoam 204. As shown in **Fig. 3g-h**, TL207 was also evaluated with M9-supplemented feed medium (336 g/L of glucose, 140 g/L of xylose, 23 g/L of arabinose, 4 mL/L of Antifoam 204, as well as 13.56 g/L Na_2_HPO_4_, 6 g/L KH_2_PO_4_, 1 g/L NaCl, and 2.25 g/L (NH_4_)_2_SO_4_, 0.24 g/L MgSO_4_ x 7 H_2_O, 11 mg/L CaCl_2_ x 2 H_2_O, and 2.73 mg/L FeSO_4_ x 7 H_2_O). After demonstrating the efficacy of the M9-supplemented medium, it was subsequently used for all remaining strains tested in bioreactors. The feeding medium consisted of 336 g/L of glucose, 140 g/L of xylose, 23 g/L of arabinose, 4 mL/L of Antifoam 204, as well as 13.56 g/L Na_2_HPO_4_, 6 g/L KH_2_PO_4_, 1 g/L NaCl, and 2.25 g/L (NH_4_)_2_SO_4_, 0.24 g/L MgSO_4_ x 7 H_2_O, 11 mg/L CaCl_2_ x 2 H_2_O, and 2.73 mg/L FeSO_4_ x 7 H_2_O.

External pumps (Watson Marlow 120U/DV analogue control variable speed pump) were used to deliver the feed, using I/S 13 MasterFlex tubing with a section of I/14 Pharmed tubing positioned in the pump head to improve feed accuracy. The feeding rates were adjusted at each sampling point to attempt to maintain a glucose concentration between 10 g/L-30 g/L. Glucose was measured overtime by an offline sugar analyzer (YSI 2900 Series). Cultivations were terminated when agitation reached minimum values (350 rpm) and glucose utilization rates became negligible. Samples were taken periodically from the bioreactors to evaluate bacterial growth (OD_600_), residual glucose in the bioreactor, and to analyze extracellular metabolites (xylose, glucose, arabinose, lactate, acetate, pyruvate, protocatechuate, catechol, muconic acid).

### Calculation of bacterial performance metrics

Muconate titers (g/L) represent the total sum of both isomers analyzed over time. Muconate productivities were calculated as the muconate titer (g/L) divided by the corresponding each time point (h). Muconate C-mol yield were calculated as the C-mol generated from muconate divided by the C-mol utilized from glucose, xylose, and arabinose: [(mM muconate × 6) / mM (glucose × 6 + mM xylose × 5 + mM arabinose × 5) × 100%]. For bioreactor cultivations, yields were corrected for dilution due to feeding and base addition for pH control.

### Transcriptomics analysis

Shake-flask experiments in triplicates were conducted in 25mL M9 with 40 g/L mock hydrolysate, for the samples of transcriptomics analysis. All samples were taken when the OD_600_ reached between 1/8 to 1/4 of maximum OD_600_, which we defined as mid-log phase. Samples were taken and centrifuged in a pre-chilled centrifuge at 4000 *g* for 10 minutes. Supernatants were discarded and cell pellets were frozen on liquid nitrogen and transferred to a −80 °C freezer. After library preparation and sequencing by Genewiz, the RNA-seq data were analyzed using a combination of Geneious Prime software and KBase platform. In Geneious Prime, genome annotation was optimized by exclusively retaining ORF features to prevent locus tag conflicts – heterologous genes received new locus tags, and all properties except locus_tag were systematically removed from CDS annotations through batch editing. For KBase analysis, a narrative was created where noninterleaved paired-end sequencing reads and the curated genome in .gb format were imported via URL upload, with critical adjustments to the genome file header ensuring KBase compatibility. Reads were consolidated into an RNA-seq sample set and aligned to the annotated genome using Bowtie2. Resulting alignments underwent transcript assembly and quantification via StringTie, enabling subsequent differential expression analysis with DESeq2 (*75*) (see **Supplementary File**, provided separately).

### Oxygraph

To evaluate the collective contribution of the engineered promoter and RBS on *catA1*, oxygen consumption was measured in engineered strain lysates. The strains were grown in LB medium to mid-log phase, harvested by centrifugation, washed with M9 media, and lysed chemically using B-PER (Thermo Fisher) according to the manufacturer’s protocol. Lysates were clarified by centrifugation (16,000 x g, 15 min) and filtration through 0.45 µm filter. The protein concentration of each lysate was determined using a BCA protein assay kit (Pierce, Thermo Fisher) using bovine serum albumin (BSA) as the standard. Oxygen consumption was monitored using a Clark-type electrode (Oxygraph, Hansatech) with 50% saturated KCL as the electrolyte, and this served as a rough proxy for CatA1 activity. The system was calibrated by a two-point method with air-saturated buffer and sodium dithionite according to the manufacturer’s protocol. The reaction was performed at 25 °C following the addition of the 10 µL lysate into 1 mL of air-saturated 25 mM HEPES, 50 mM NaCl pH 7.5 and 200 µM catechol (< 0.1 % (v/v) final DMSO). The assay was initiated by the addition of 10 μL of the cleared lysate to a total of 1 mL assay volume. The observed reading was corrected with the background readings prior to the addition of the lysate in biological triplicates.

### Quantitation of muconic acid isomers and aromatic byproducts

Muconic acids (*cis,cis*-muconic acid and *cis,trans*-muconic acid), along with aromatic compounds, were quantified using an Agilent 1290 Infinity II series ultra-high-performance liquid chromatography system equipped with a diode array detector (DAD) and separation was achieved using reverse-phase chromatography on a Waters Acquity UPLC BEH column (*76*).

### Analysis of sugars and co-products

Sugars and co-products from biological cultivation experiments were analyzed by high-performance liquid chromatography (HPLC) (*77*), with minor modifications. To enable quantification of pyruvic acid and address the partial co-elution of pyruvic acid and xylose, the column temperature was increased from 55 °C to 80 °C to achieve sufficient separation. Additionally, a diode array detector (DAD) was employed at a quantitative wavelength of 210 nm for pyruvic acid analysis, as xylose exhibits minimal to no absorbance at this wavelength.

### Techno-economic analysis and life cycle assessment

Dedicated process simulations for the synthesis of muconic acid and adipic acid from a corn stover hydrolysate were developed using Aspen Plus V14 (AspenTech). The production of the mixed sugars stream considers the processing of 2,000 dry tons of corn stover per day via Deacetylation and Mechanical Refining (DMR), following the assumptions detailed in Mokwatlo *et al*. (*63*). The bioconversion of sugars to muconic acid and catalytic upgrading to adipic acid was modeled both with process engineering parameters (detailed in Johnson *et al*. (*14*) and refined as needed) and with experimental data discussed herein. Further details are presented in the **Tables S6**-**S7**.

Material and energy balances obtained from process simulations were utilized to estimate the capital expenditures (CAPEX) and operational expenses (OPEX) tied to the chemical plants. The economic performance of the production processes was benchmarked using the minimum selling price (MSP) of either muconic acid or adipic acid. This metric was obtained using discounted cash flows, which are solved to yield the sales price (i.e., MSP) required to achieve a net present value of zero over a 30-year lifetime. Additional details concerning process simulation and TEA results are shown in **Supplementary Note 3** and **Tables S7**-**S10**.

For LCA, a consistent methodology as presented in Mokwatlo *et al*. (*63*) was considered based on mass and energy balances generated by process simulations. Additional information concerning the supply chain of chemical and energy for the production of adipic and muconic acids were retrieved from the R&D GREET version 2025 (*65*). The analysis is on a cradle-to-gate basis and the functional unit considered is one kg of adipic or muconic acid produced from DMR sugars. The main metrics in the analysis are the GHG emissions (kg CO_2_e/kg of product), fossil energy consumption (MJ/kg of product), and water consumption (L/kg of product), and further compared to conventional adipic acid production. To ensure a robust and comprehensive evaluation, LCA modeling was additionally on the same life cycle inventories (LCIs) using the Brightway version 3.6.6, utilizing background datasets from the ecoinvent v3.11 database (*66*). Additional details regarding these LCA results are shown in **Supplementary Note 4**, with the main LCIs detailed in **Tables S11**-**S14**.

## Supporting information

RNA Sequence

## Acknowledgements

This work was authored in part by Alliance for Energy Innovation, LLC, the manager and operator of the National Laboratory of the Rockies (NLR) for the U.S. Department of Energy (DOE) under Contract No. DE-AC36-08GO28308. Funding was provided by the U.S. Department of Energy Office of Alternative Fuels and Feedstocks Office (AFFO) under the DOE Agile BioFoundry. We acknowledge the support of Gayle Bentley, Dan Fishman, and Lisa Guay at AFFO. We thank Keven Dooley and Tracy Hodges at NLR for helpful discussions. This work was authored in part by Oak Ridge National Laboratory, which is managed by UT-Battelle, LLC, for the U.S. Department of Energy (DOE) under contract DE-AC05-00OR22725. The views expressed in the article do not necessarily represent the views of the DOE or the U.S. Government. The U.S. Government retains and the publisher, by accepting the article for publication, acknowledges that the U.S. Government retains a nonexclusive, paid-up, irrevocable, worldwide license to publish or reproduce the published form of this work, or allow others to do so, for U.S. Government purposes.

## Data availability

The data supporting the findings of this study are available within the article and its Supplementary Information. RNA-seq data are provided as a separate Supplementary File. The sequences of all plasmids constructed in this study can be found under the GenBank accession numbers PZ618242-PZ618254 (https://www.ncbi.nlm.nih.gov/genbank), and the Addgene plasmid numbers 260362-260374 (https://www.addgene.org). The genome details are available upon request. Source data are provided with this paper.

