## Supplementary Information for "Pathway selection for arabinose utilization in *Pseudomonas putida* reveals a rate-yield tradeoff in muconic acid production from lignocellulosic sugars"

**Supplementary note 1 |** Energy limited conditions from glucose metabolism in engineered strain

**Supplementary note 2 |** Investigation of metabolic intermediates accumulation due to overflow metabolism in TL207

**Supplementary note 3 |** Process simulation and techno-economic analysis

**Supplementary note 4 |** Life cycle assessment from Brightway framework utilizing ecoinvent v3.11 database

**Figure S1** | Growth curves of LC224, LC237, and LC357 in plate cultivation

**Figure S2** | Metabolic profiles of LC224, LC237, and LC357 in shake-flask cultivation

**Figure S3** | Metabolic profiles of LC224 and LC237 in fed-batch bioreactor cultivation

**Figure S4** | Effect of AraE deletion on LC224 – growth curves of LC224, LC237, and TL015 in plate cultivation

**Figure S5** | Metabolic profiles of LC237 and DK037 in shake-flask cultivation

**Figure S6** | Growth curves of LC237, TL170, TL172, TL174, and TL207 in plate cultivation

**Figure S7** | Metabolic profiles of LC237, TL170, TL172 and TL174 in shake-flask cultivation

**Figure S8** | Phenotypic observation of dark brown coloration in shake-flask cultivation

**Figure S9** | Metabolic profiles of LC237, TL170, and TL207 in shake-flask cultivation

**Figure S10** | Metabolic profiles of LC237 and TL207 in fed-batch bioreactor cultivation

**Figure S11** | Metabolic profiles of TL207 during medium optimization in fed-batch bioreactor cultivation

**Figure S12** | Metabolic profiles of TL207, TL477, and TL831 in shake-flask cultivation

**Figure S13** | Metabolic profiles of TL207 and TL831 in fed-batch bioreactor cultivation

**Figure S14** | Metabolic profiles of TL207, DK092, and DK093 in shake-flask cultivation

**Figure S15** | Metabolic profiles of TL207 and TL565 in fed-batch bioreactor cultivation

**Figure S16** | Growth curves of LC357 and TL809 in plate cultivation

**Figure S17** | Metabolic profiles of LC357, TL809, and DK051 in shake-flask cultivation

**Figure S18** | Metabolic profiles of TL207, DK051, DK062, and DK063 in shake-flask cultivation

**Figure S19** | Metabolic profiles of TL207, DK062, and DK063 in fed-batch bioreactor cultivation

**Figure S20** | Strain performance comparison of TL207, DK062, and DK063 in fed-batch bioreactor cultivation

**Figure S21** | Main operations required for the bioconversion of mixed sugars to muconic acid and posterior catalytic upgrading to adipic acid

**Figure S22 |** Expanded techno-economic analysis and life cycle assessment for adipic acid (AA) and muconic acid (MA) production

**Figure S23 |** Techno-economic analysis and life cycle assessment for adipic acid (AA) and muconic acid (MA) production from TRY achieved at maximum rate

**Figure S24 |** Sensitivity analysis of low-emission electricity sources on adipic acid (AA) and muconic acid (MA) production from DMR sugars

**Figure S25** | Life cycle assessment of adipic acid (AA) and muconic acid (MA) production from TRY achieved at final and maximum rate (Database: Brightway using ecoinvent v3.11)

**Figure S26** | Life cycle assessment of adipic acid (AA) production across various environmental impact categories (Database: Brightway using ecoinvent v3.11)

### Supplementary note 1 | Energy limited conditions from glucose metabolism in engineered strain

*Pseudomonas putida* (*P. putida*) naturally lacks 6-phosphofructo-1-kinase (Pfk) and relies almost exclusively on the Entner–Doudoroff (ED) pathway for glucose catabolism, yielding only one net ATP per glucose molecule (*1*). To balance fluxes, *P. putida* utilizes the Embden–Meyerhof–Parnas (EMP) pathway in the gluconeogenic direction, creating a cyclic EDEMP pathway. In our engineered strains, we deleted *pgi-2*, which encodes the predominant glucose-6-phosphate isomerase to disrupt the gluconeogenic recycling phase of the EDEMP cycle and redirect carbon toward the shikimate pathway for muconate production (*2*). Although a secondary isomerase (*pgi-1*) remains, this deletion restricts gluconeogenic recycling, thereby pushing the host into an energy-limited state.

By modifying the EDEMP cycle, the engineered strains may possess minimal energetic buffering capacity. In this energetically constrained background, high activity of the proton-coupled AraE transporter rapidly depletes the proton motive force (PMF). This PMF drain subsequently starves the ATP synthase, thereby impairing overall ATP generation (*3*). Ultimately, this energetic depletion broadly interferes with energy-dependent active transport systems, including those required for glucose metabolism.

### Supplementary note 2 | Investigation of metabolic intermediates accumulation due to overflow metabolism in TL207

Among the strains evaluated, TL207 exhibited improved sugar consumption and muconate production; however, we also observed an increase in the transient accumulation of acetate, lactate, and pyruvate s during the late-exponential growth phase (**Extended Data Figure 1**). Accumulation of these intermediates are indicative of overflow metabolism, which typically occurs when the carbon flux toward pyruvate exceeds the processing capacity of the TCA cycle (*4-6*). We therefore posited that overflow metabolism could be mitigated by attenuating carbon flux into the TCA cycle and/or tuning the expression of *glf* to avoid excessive glucose influx. To evaluate these hypotheses, we first decreased the promoter strength of the arabinose-oxidative pathway to reduce flux toward the TCA cycle. We then added a mutant of the citrate synthase (*gltA*$^{R164L}$; strain TL831) (*7*), which was previously reported to promote the entry of acetyl-CoA into the TCA cycle by resisting feedback inhibition from high levels of NADH or ATP. These modifications resulted in a modest reduction of intermediate accumulation, with ~ 40 % decrease in total overflow metabolites in both shake-flask (**Fig. S12**) and bioreactor experiments (**Fig. S13**).

In parallel, we slightly downregulated *glf* expression in TL207 to balance the rate of glucose transport with the metabolic capacity of the cell and prevent carbon overflow (strains DK092 and DK093; **Extended Data Table 1**). As expected, this approach reduced glucose uptake rates, thereby lowering overflow accumulation as shown in **Extended Data Figure 1**. Although DK092 and DK093 exhibited comparable muconate yields to TL207, their overall glucose utilization was slower (**Extended Data Figure 1**, **Fig. S14**), which could potentially compromise productivity in bioreactor operations. Collectively, while overflow metabolism can be partially mitigated by adjusting TCA cycle flux or decreasing glucose uptake, these strategies alone may be insufficient to improve muconate yield.

### Supplementary note 3 | Process simulation and techno-economic analysis

Production of muconic acid through the bioconversion of mixed sugars is carried out in aerobic vessels at specific productivities, yields, and titers, as determined experimentally for the individual points in the main text of the manuscript. Reactions for biomass growth and product formation are shown in Table TEA1 for the three sugar species in corn stover hydrolysate: glucose, xylose, and arabinose. The reactions specified for xylose and arabinose in **Table S6** are derived from the equations starting with glucose by multiplying all non-sugar compounds by a factor of 5/6. Losses of sugars due to contamination (Glucose to biomass, **Table S6**) are fixed at 3% for all cases. Further details are presented in the **Table S7-S10**.

### Supplementary note 4 | Life cycle assessment from Brightway framework utilizing ecoinvent v3.11 database

Besides LCA methodology from R&D GREET version 2025 used in main manuscript, a parallel LCA methodology from Brightway version 3.6.6 (*8*) was also employed to estimate usual environmental impact categories (such as GHG emissions) and additional ones (**Figs. S25-S26**). Cradle-to-gate life cycle assessments were conducted for each muconic acid and adipic acid production pathway to determine the environmental impacts in comparison to fossil-based adipic acid production. The scope of the assessment includes the collection, transportation, and conversion of biomass feedstocks into DMR mixed sugars, the production of muconic acid, and final conversion to adipic acid. The functional unit of analysis is 1 kg of adipic acid or muconic acid. Life cycle inventory data for the collection/harvesting and transportation of corn stover is derived from the ecoinvent v3.11 inventory for corn production (*8*). The impacts for corn stover are determined using economic allocation, assuming 15% of impacts are attributed to corn stover, and the remainder to corn grain (*9*). In addition, phosphorus, nitrogen and potassium fertilizer are added to compensate for the removal of nutrients from the soil due to corn stover harvesting (*9*). The life cycle inventory data for muconic and adipic acid are based on the process models developed herein, with background data from ecoinvent v3.11 database (*8*). The ReCiPe 2016 Midpoint Hierarchist impact assessment method (*10*) was used to assess 13 impact categories including: acidification (kg $SO_2$ eq), ecotoxicity (kg 1,4-DCB eq), freshwater eutrophication (kg P eq), marine eutrophication (kg N eq), fossil resource use (kg oil eq), human toxicity (kg 1,4-DCB eq), ionizing radiation (kg Co-60 eq), land use ($m^2$a crop eq), mineral resource use (kg Cu eq), ozone depletion (kg CFC-11 eq), particulates formation (kg PM2.5 eq), photo-oxidant formation (kg NOx eq), and water use ($m^3$). To analyze greenhouse gas (GHG) emissions, the Intergovernmental Panel on Climate Change (IPCC) 2021 GWP characterization method was used (*11*).

From an LCA standpoint, all the product routes for adipic acid resulted in a 73-80% reduction in GHG emissions compared to fossil-derived adipic acid. These results include biogenic carbon credits, or negative emissions, associated with the biomass-derived glucose source. Without the biogenic carbon credit, assuming it is treated as net-neutral throughout the products lifecycle, a 53-61% reduction in GHG emissions compared to the fossil-scenario is observed. 37-45% of the remaining GHG emissions for adipic acid are associated with the production of mixed sugars, followed by ammonium hydroxide, contributing 19-22%. Electricity and heat requirements for adipic acid together contribute an additional 21-24% to total GHG emissions. In addition to reducing GHG emissions, this pathway results in a 24% reduction in cumulative energy demand, on average, compared to fossil-derived adipic acid. However, this bio-based pathway has 15 times higher marine eutrophication impacts (nitrogen emissions), and 8-9 times higher land and water use, on average, compared to fossil-derived adipic acid (as detailed in **Tables S11-15** and **Figs. S25-S26**). This is largely due to the impacts of agriculture associated with corn stover harvesting. Therefore, other bio-based feedstocks with less land, water and fertilizer requirements may have the potential to reduce the magnitude of these trade-offs.

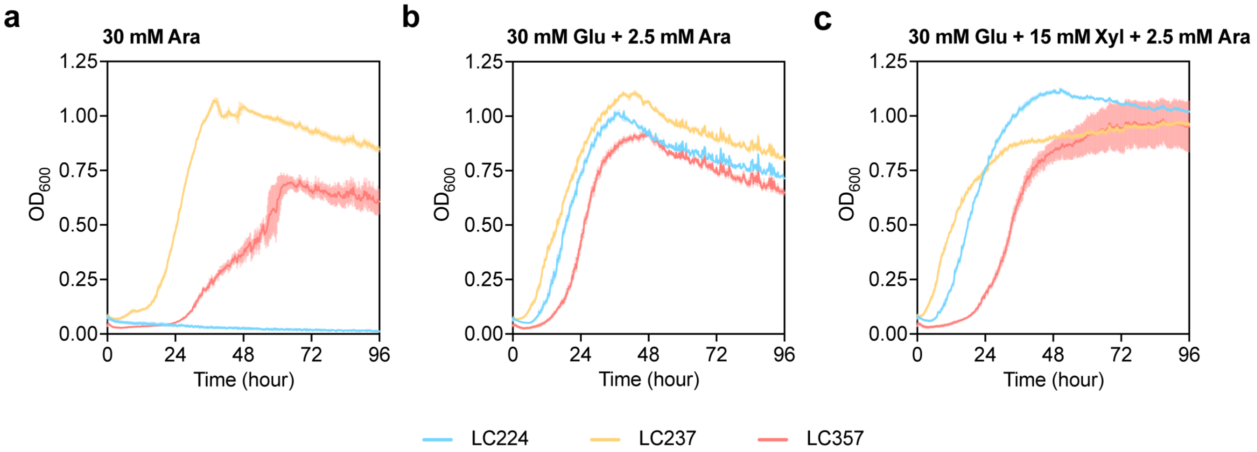


**Figure S1 | Growth curves of LC224, LC237, and LC357 in plate cultivation.** Growth curves of LC224, LC237, and LC357 in plate cultivation for comparison. Strains were grown on M9 medium supplemented with 30 mM arabinose (**a**), 30 mM glucose and 2.5 mM arabinose (**b**), and 30 mM glucose, 15 mM xylose, and 2.5 mM arabinose (**c**). **a** is adapted from **Fig. 2a** for comparison. Data represent the average of $n$ = 3 biological replicates. Error bars correspond to standard deviation. Numerical data are provided in a **Source Data File**.

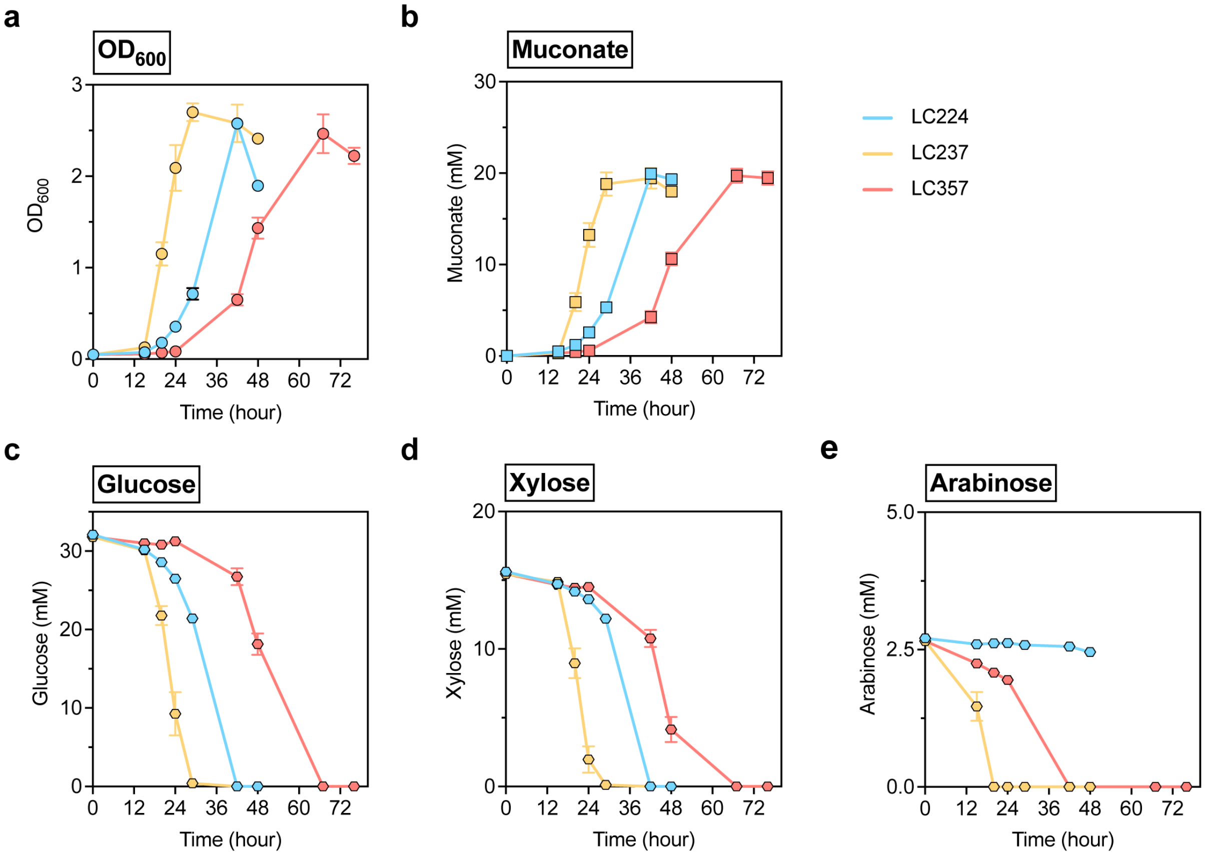


**Figure S2 | Metabolic profiles of LC224, LC237, and LC357 in shake-flask cultivation.** Metabolic profiles of LC224, LC237, and LC357 in shake-flask cultivation for comparison. Profiles show the bacterial growth ($OD_{600}$) (**a**), muconate production (**b**), and residual concentrations of glucose (**c**), xylose (**d**), and arabinose (**e**) on M9 medium supplemented with 8 g/L mock hydrolysate. Data represent the average of $n$ = 3 biological replicates. Error bars correspond to standard deviation. Numerical data are provided in a **Source Data File**.

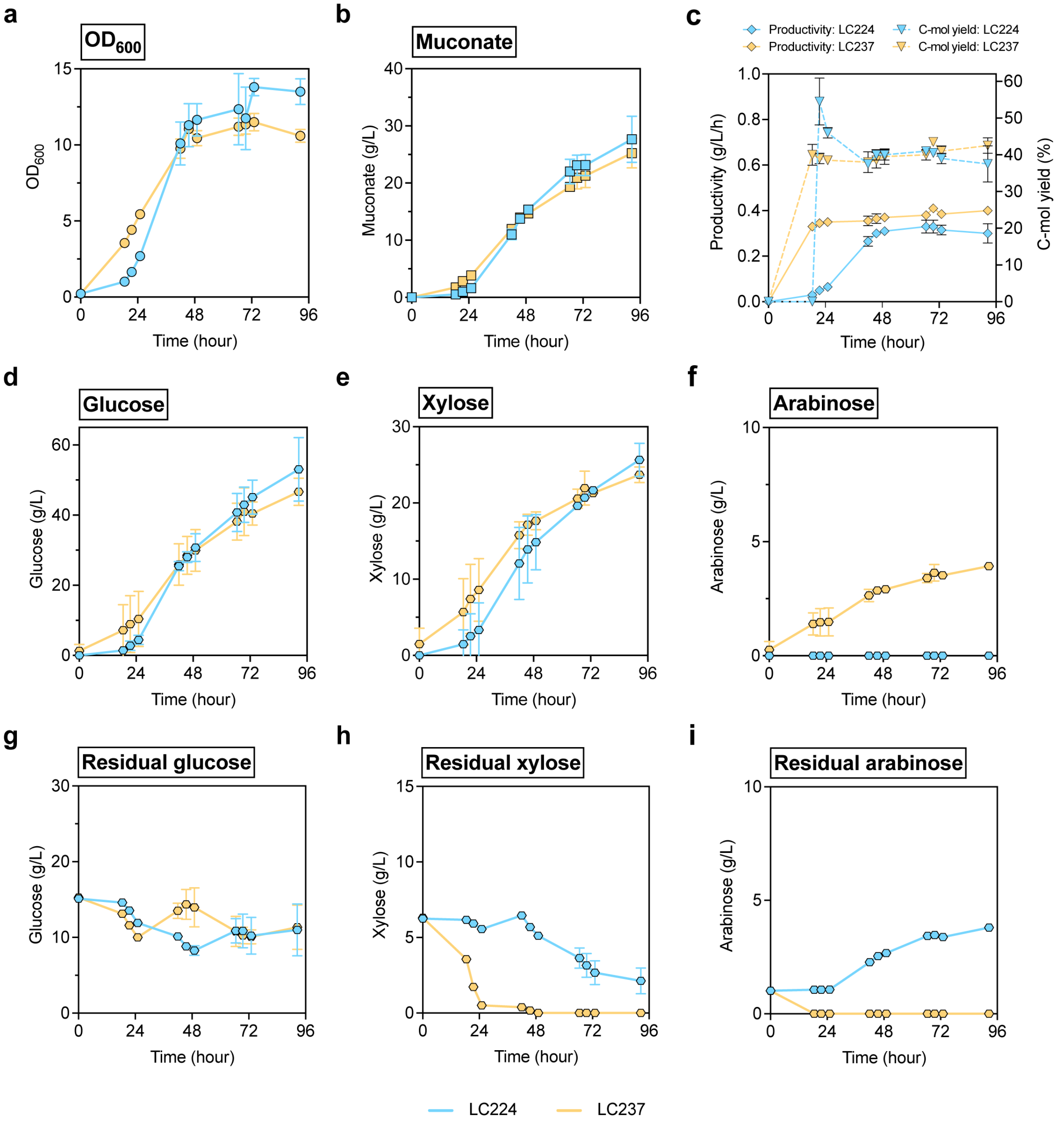


**Figure S3 | Metabolic profiles of LC224 and LC237 in fed-batch bioreactor cultivation.** Fed-batch bioreactor evaluation of LC224 and LC237 at 0.5-L scale. Profiles show the bacterial growth ($OD_{600}$) (**a**), muconate production (**b**), and comparison of muconate productivity and carbon yield (**c**). Total sugar utilization profiles are presented for glucose (**d**), xylose (**e**), and arabinose (**f**), with corresponding residual concentrations of glucose (**g**), xylose (**h**), arabinose (**i**) in the bioreactor. Rate: titer/time (g/L/h), Yield (C-mol %): [(mM muconate × 6) / mM (glucose × 6 + mM xylose × 5 + mM arabinose × 5) × 100%]. Data represent the average of $n$ = 2 biological replicates. Error bars correspond to absolute error between duplicates. Numerical data are provided in a **Source Data File**.

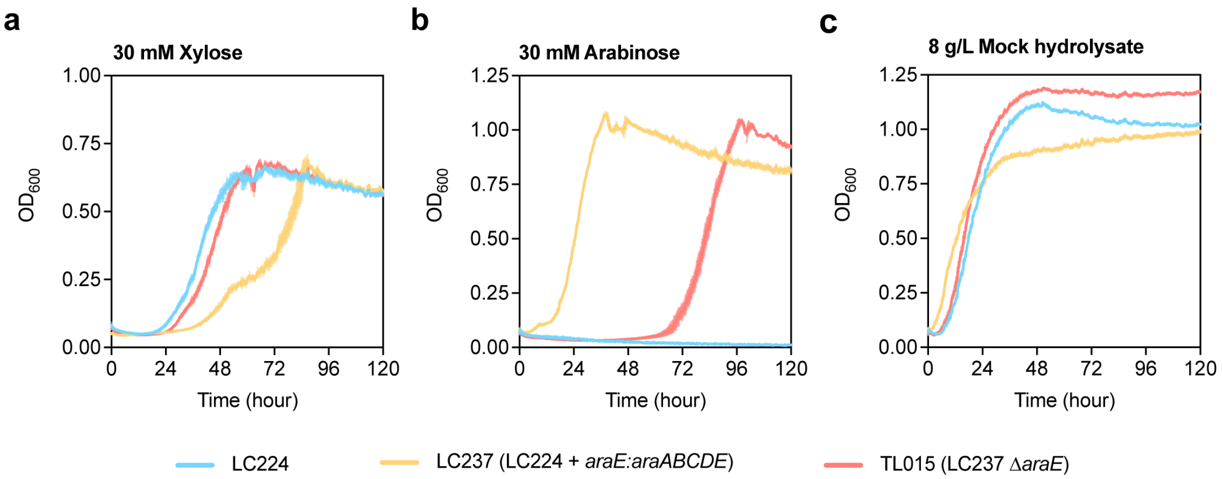


**Figure S4 | Effect of AraE deletion on LC224 – growth curves of LC224, LC237, and TL015 in plate cultivation.** Growth curves of LC224, LC237, and TL015 (LC224 Δ*araE*) in plate cultivation for comparison. Strains were grown on M9 medium supplemented with 30 mM xylose (**a**), 30 mM arabinose (**b**), and 8 g/L mock hydrolysate (**c**). Data represent the average of $n$ = 3 biological replicates. Error bars correspond to standard deviation. Numerical data are provided in a **Source Data File**.

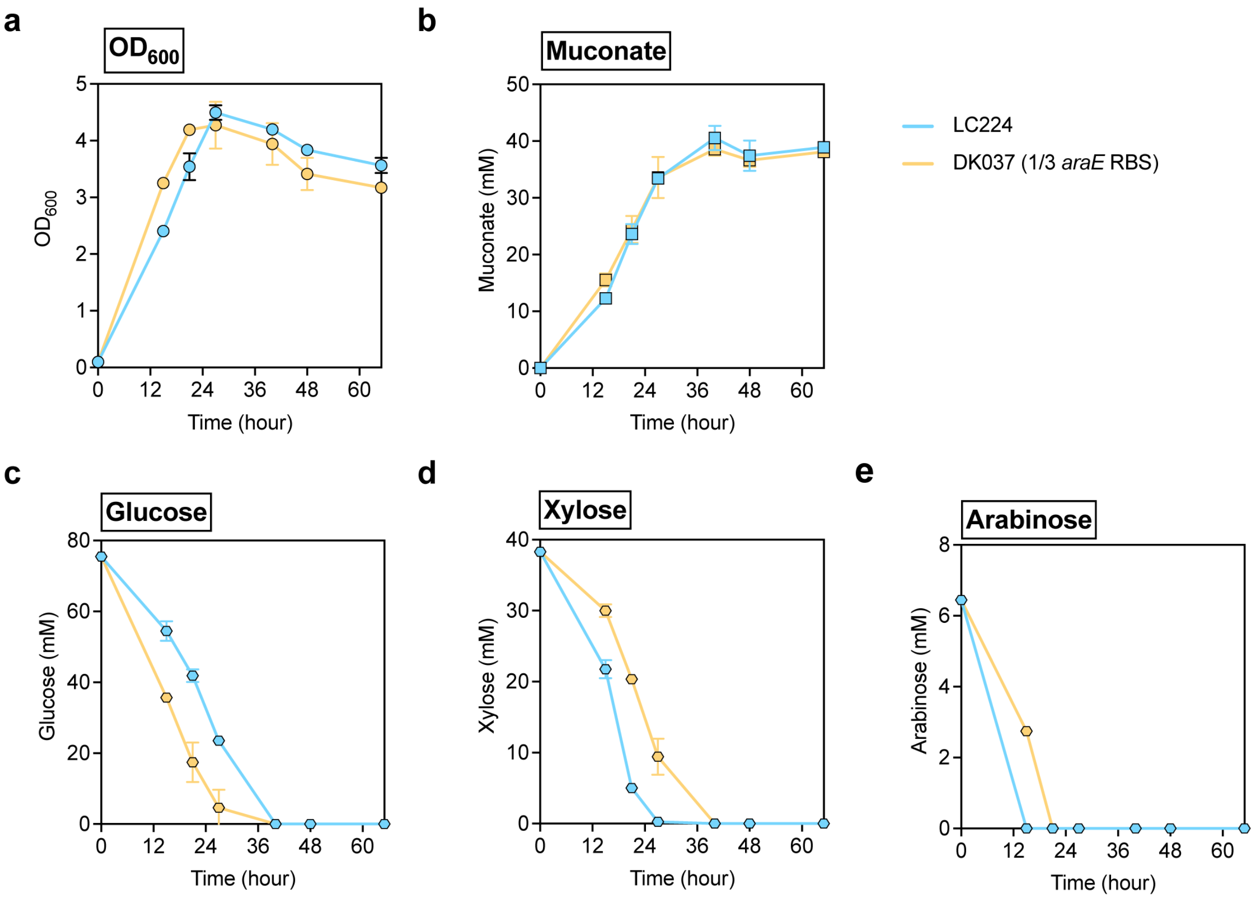


**Figure S5 | Metabolic profiles of LC237 and DK037 in shake-flask cultivation.** Metabolic profiles of LC237 and DK037 (LC237 1/3 RBS strength *araE*) in shake-flask cultivation for comparison. Profiles show the bacterial growth ($OD_{600}$) (**a**), muconate production (**b**), and residual concentrations of glucose (**c**), xylose (**d**), and arabinose (**e**) on M9 medium supplemented with 25 g/L mock hydrolysate. Data represent the average of $n$ = 3 biological replicates. Error bars correspond to standard deviation. Numerical data are provided in a **Source Data File**.

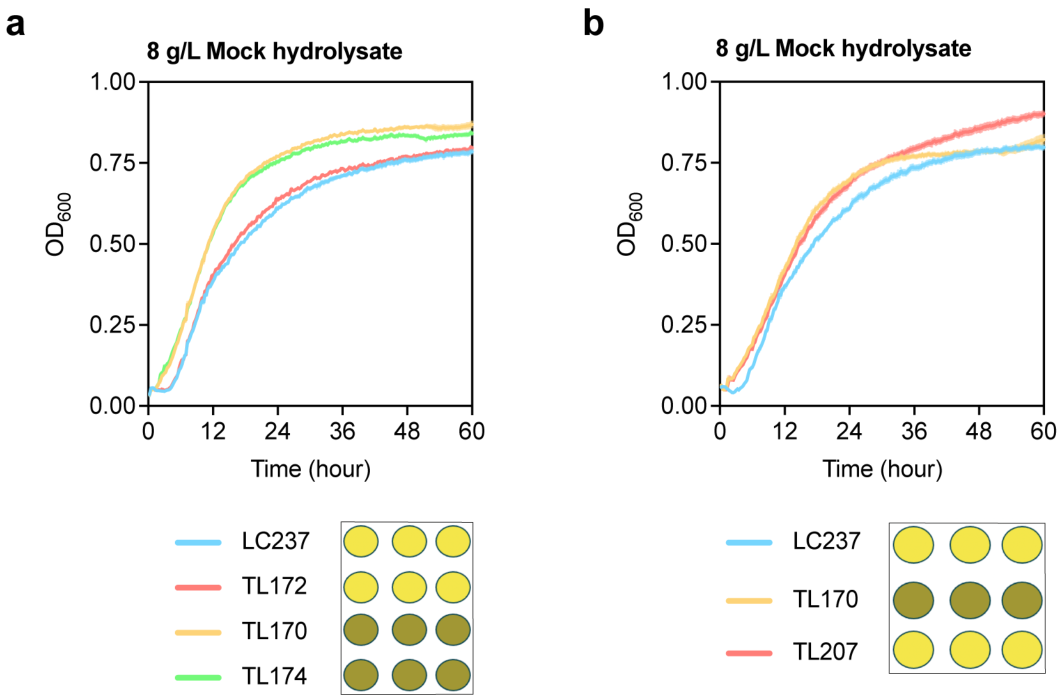


**Figure S6 | Growth curves of LC237, TL170, TL172, TL174, and TL207 in plate cultivation. a-b** Growth curves of *glf*-expressing strains. LC237, TL170, TL172, and TL174 **(a)** and LC237, TL170, and TL207 **(b)** in plate cultivation for comparison. All strains were grown on M9 medium supplemented with 8 g/L mock hydrolysate. The yellow and brown circles indicate the color of the growth medium after growth in the plate assays. Yellow indicates the normal color of the medium after *P. putida* growth, while the brown color indicates darkening of the growth medium, potentially due to catechol accumulation in TL170 and TL174 (**a**) and TL170 (**b**). Data represent the average of $n = 4$ biological replicates. Error bars correspond to standard deviation. Numerical data are provided in a **Source Data File**.

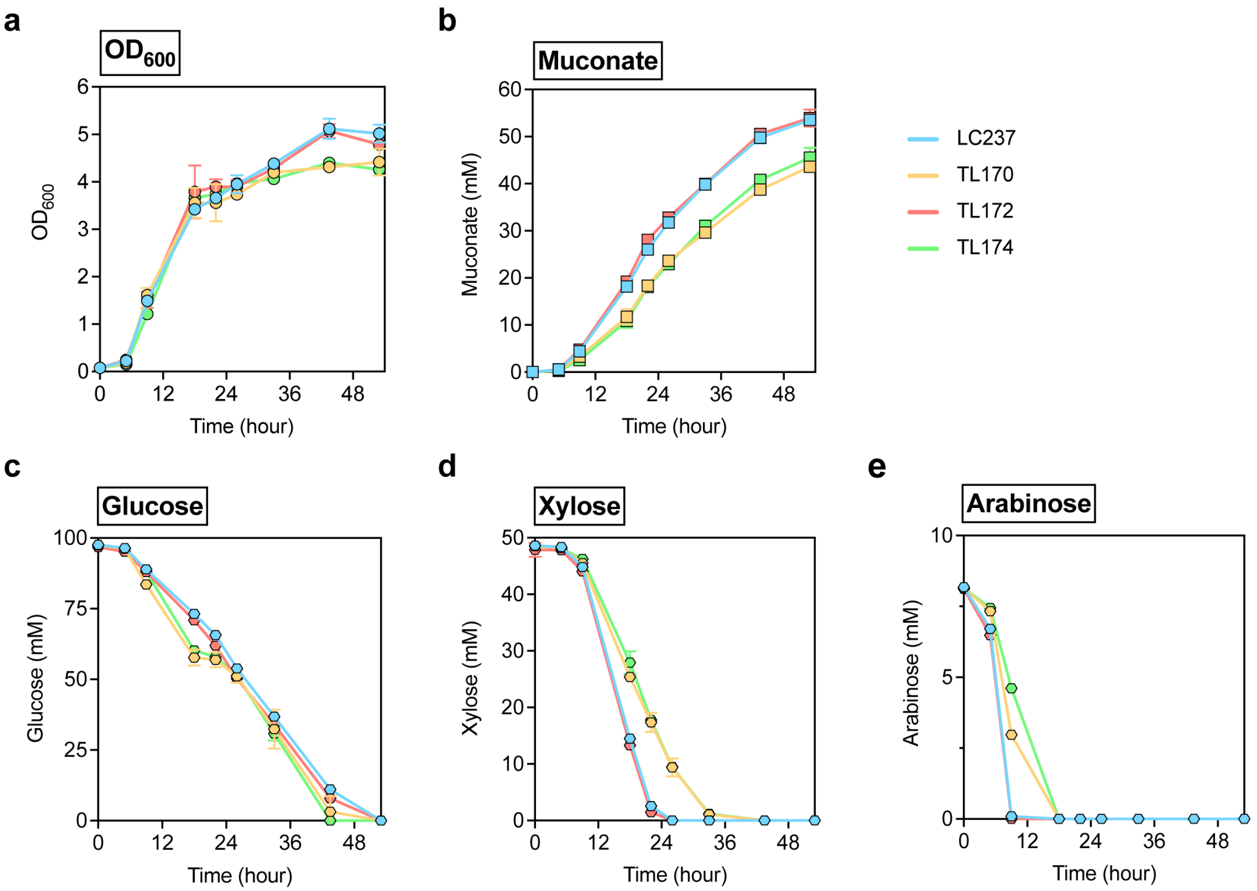


**Figure S7 | Metabolic profiles of LC237, TL170, TL172 and TL174 in shake-flask cultivation.** Metabolic profiles of LC237, TL170, TL172 and TL174 in shake-flask cultivation for comparison. Profiles show the bacterial growth ($OD_{600}$) (**a**), muconate production (**b**), and residual concentrations of glucose (**c**), xylose (**d**), and arabinose (**e**) on M9 medium supplemented with 25 g/L mock hydrolysate. Data represent the average of $n$ = 3 biological replicates. Error bars correspond to standard deviation. Numerical data are provided in a **Source Data File**.

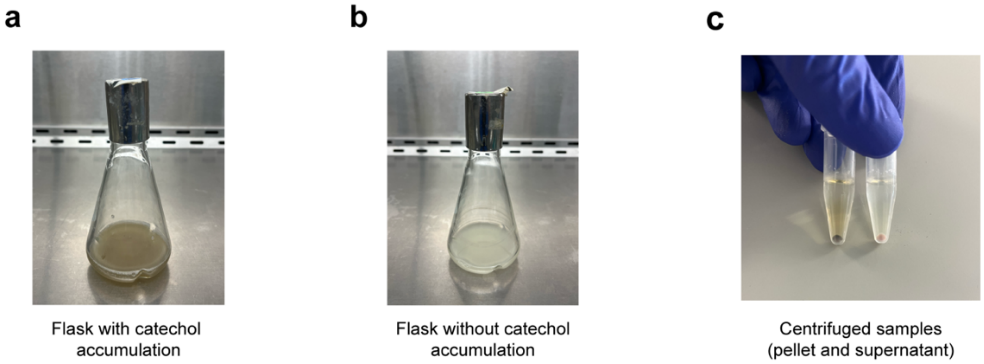


**Figure S8 | Phenotypic observation of dark brown coloration in shake-flask cultivation.** Visual comparison of color changes observed during shake-flask cultivation of T170 and LC237. **a** Dark brown colorization of the culture medium of TL170 indicates catechol accumulation. **b** The culture medium of LC237 remains yellow in the absence of catechol accumulation. **c** Centrifuged samples showing a dark brown cell pellet in catechol accumulating conditions (*left*), compared to yellow pellet in the control (*right*). **c** is adapted from **Fig. 2m**. All images were captured after 48 h of cultivation.

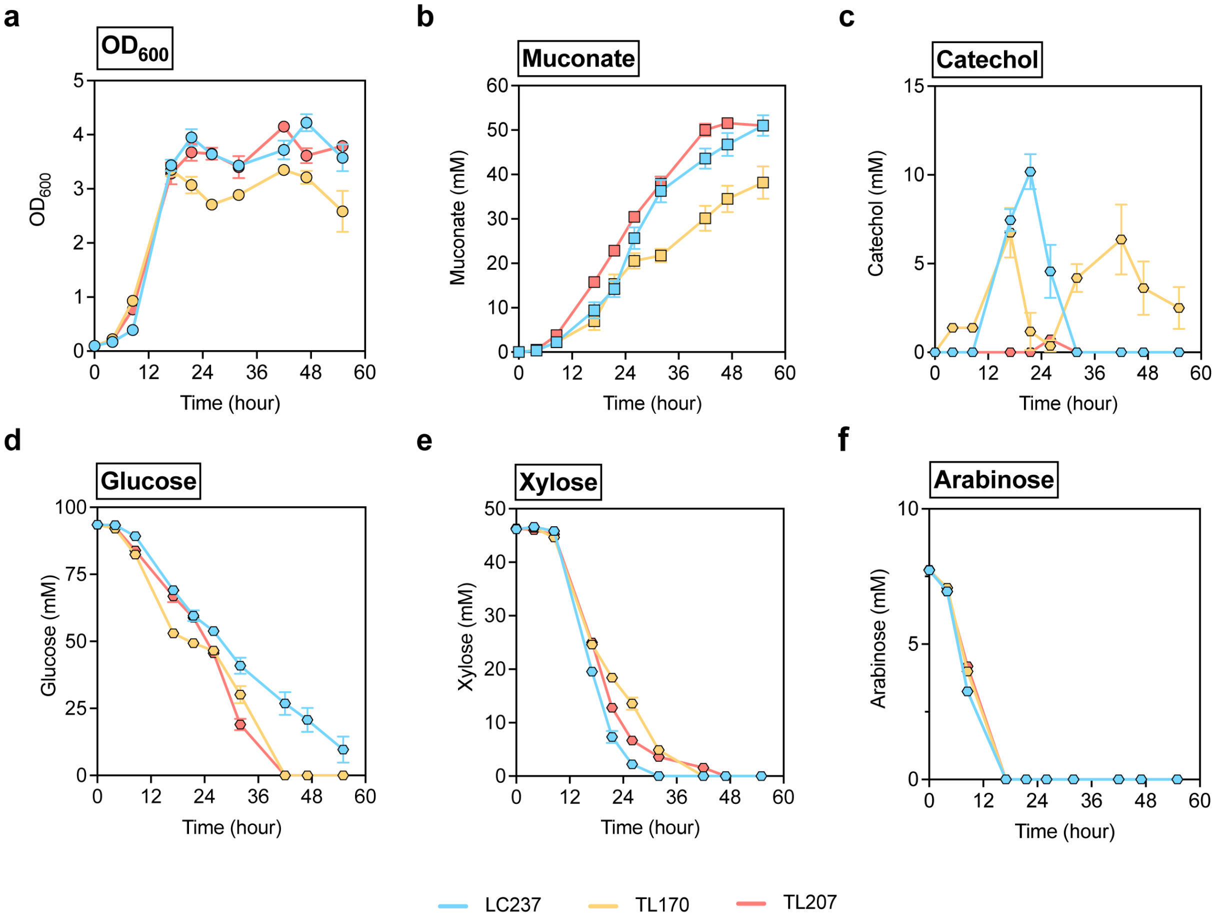


**Figure S9 | Metabolic profiles of LC237, TL170, and TL207 in shake-flask cultivation.** Metabolic profiles of LC237, TL170, and TL207 in shake-flask cultivation for comparison. Profiles show the bacterial growth ($OD_{600}$) (**a**), muconate production (**b**), catechol accumulation (**c**), and residual concentrations of glucose (**d**), xylose (**e**), and arabinose (**f**) on M9 medium supplemented with 25 g/L mock hydrolysate. Data represent the average of $n = 3$ biological replicates. Error bars correspond to standard deviation. Numerical data are provided in a **Source Data File**.

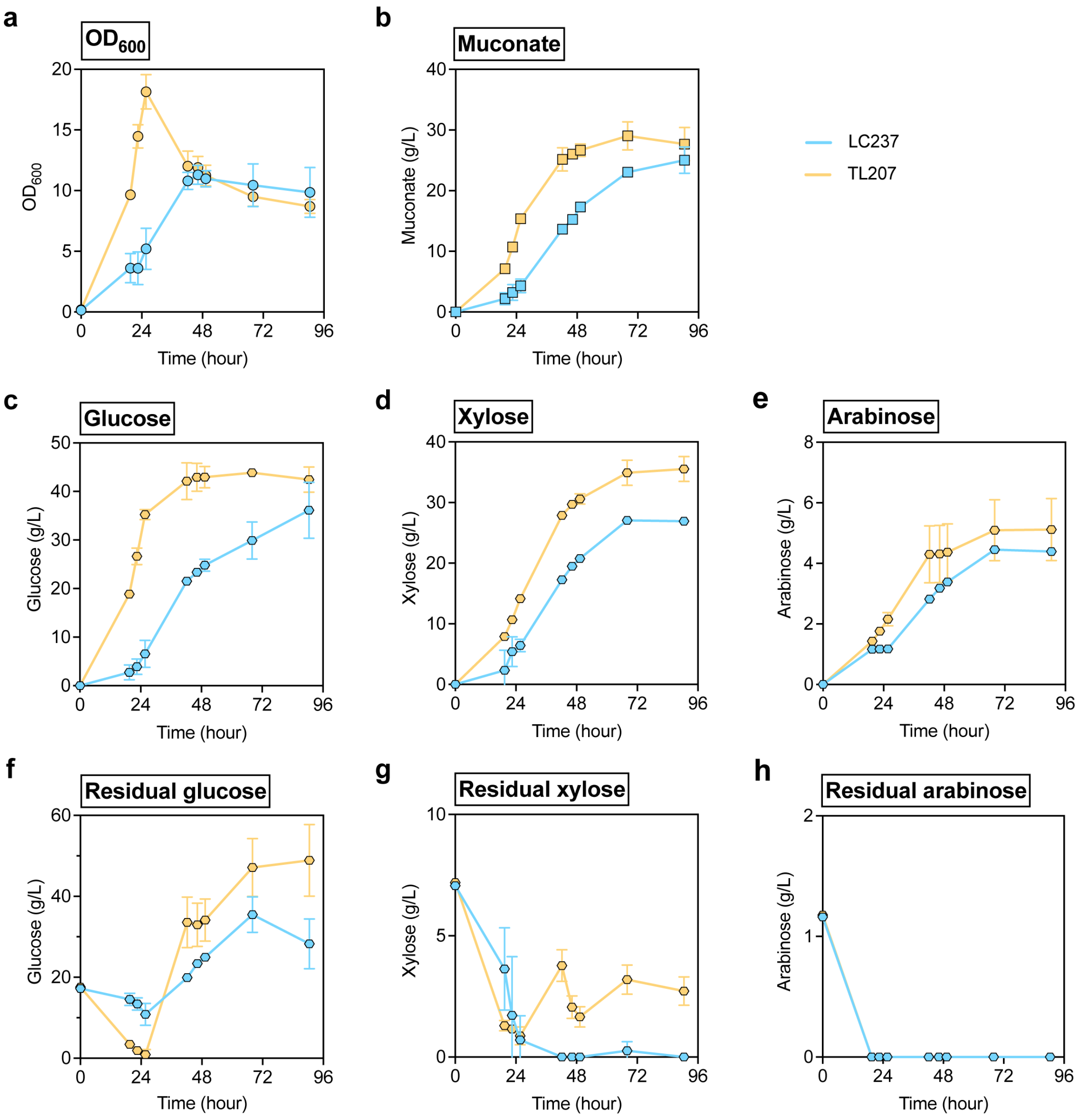


**Figure S10 | Metabolic profiles of LC237 and TL207 in fed-batch bioreactor cultivation**. Fed-batch bioreactor evaluation of LC237 and TL207 at 0.5-L scale. Profiles show the bacterial growth ($OD_{600}$) (**a**) and muconate production (**b**). Total sugar utilization profiles are presented for glucose (**c**), xylose (**d**), and arabinose (**e**), with corresponding residual concentrations of glucose (**f**), xylose (**g**), arabinose (**h**) in the bioreactor. Data represent the average of $n = 2$ biological replicates. Error bars correspond to absolute error between duplicates. Numerical data are provided in a **Source Data File**.

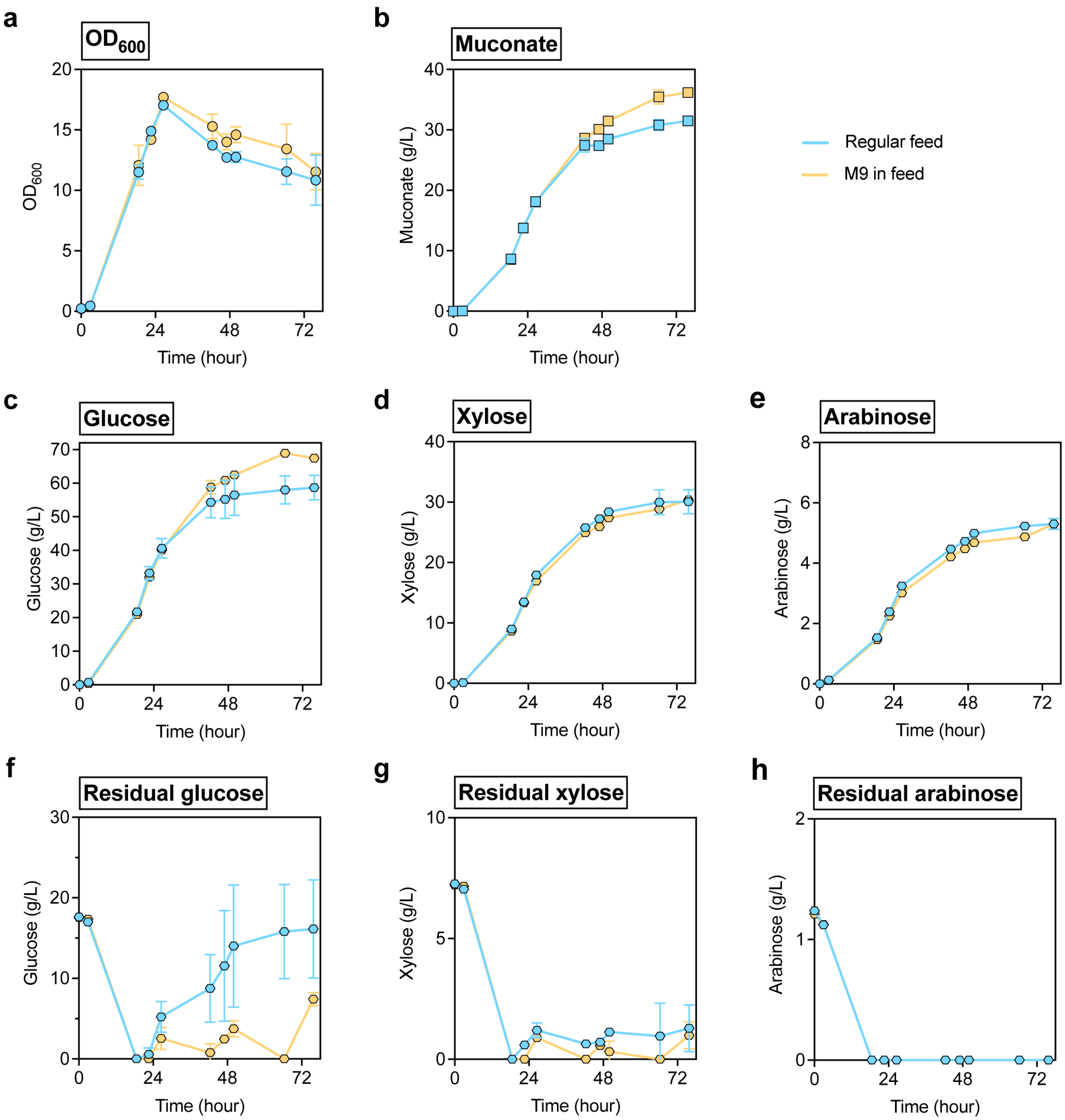


**Figure S11 | Metabolic profiles of TL207 during medium optimization in fed-batch bioreactor cultivation**. Fed-batch bioreactor evaluation of TL207 for medium optimization at 0.5-L scale, comparing regular feed and M9-supplemented feed conditions. Profiles show the bacterial growth ($OD_{600}$) (**a**) and muconate production (**b**). Total sugar utilization profiles are presented for glucose (**c**), xylose (**d**), and arabinose (**e**), with corresponding residual concentrations of glucose (**f**), xylose (**g**), arabinose (**h**) in the bioreactor. Data represent the average of $n = 2$ biological replicates. Error bars correspond to absolute error between duplicates. Numerical data are provided in a **Source Data File**.

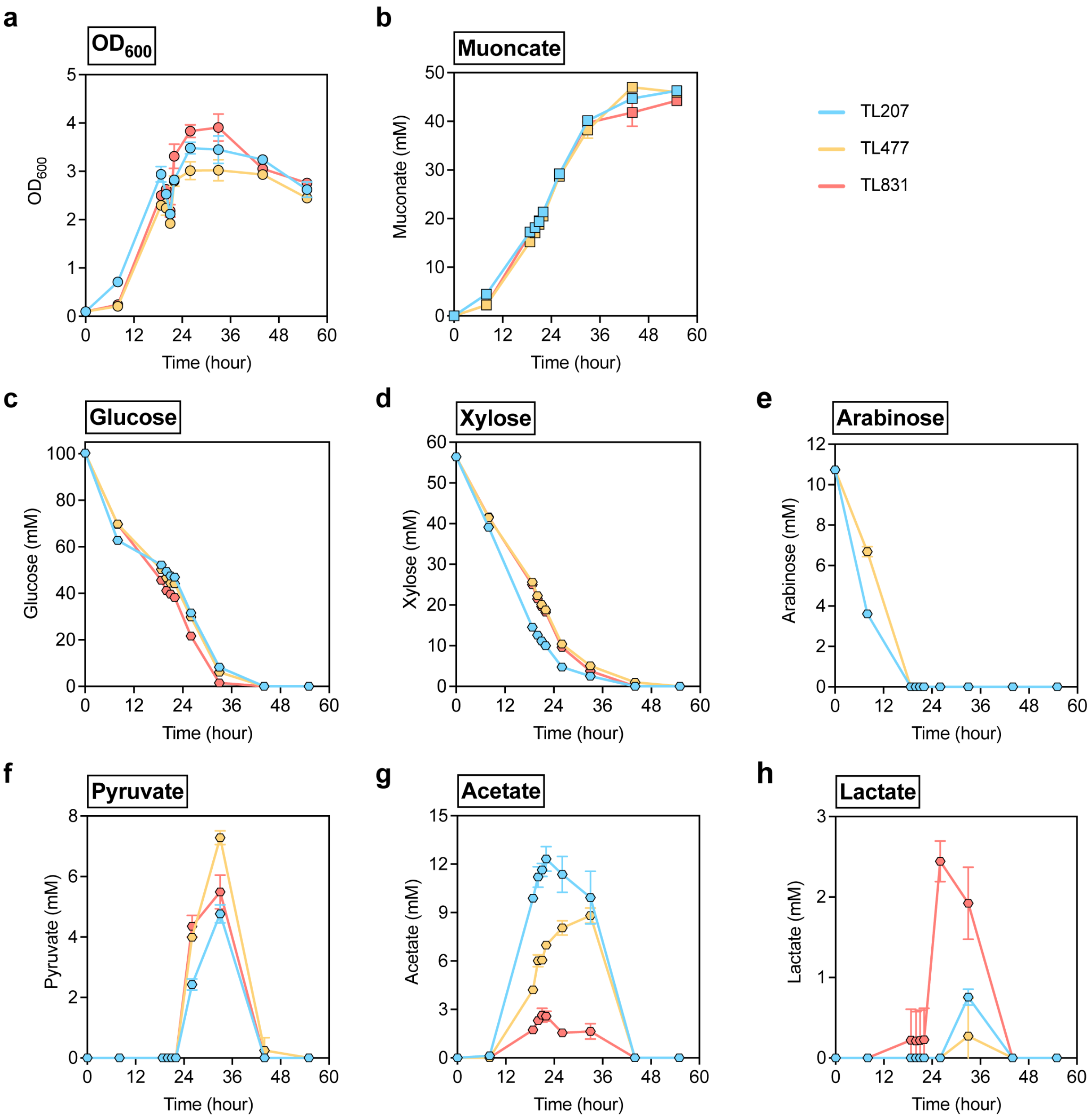


**Figure S12 | Metabolic profiles of TL207, TL477, and TL831 in shake-flask cultivation.** Metabolic profiles of TL207, TL477, and TL831 in shake-flask cultivation for comparison. Profiles show the bacterial growth ($OD_{600}$) (**a**), muconate production (**b**), residual concentrations of glucose (**c**), xylose (**d**), and arabinose (**e**), and the accumulation of pyruvate (**f**), acetate (**g**), and lactate (**h**) on M9 medium supplemented with 25 g/L mock hydrolysate. Data represent the average of $n = 3$ biological replicates. Error bars correspond to standard deviation. Numerical data are provided in a **Source Data File**.

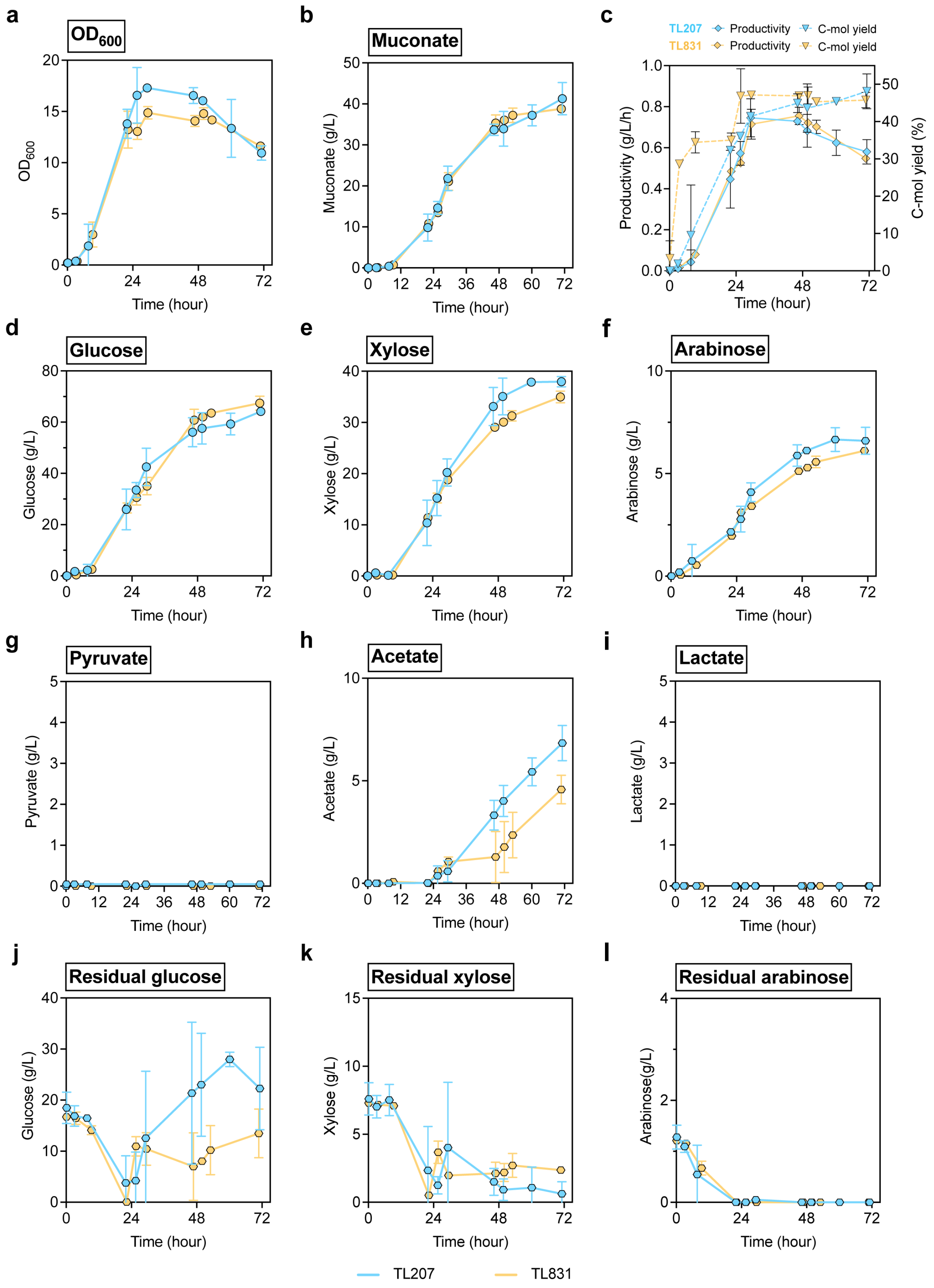

a
OD600
b
Muconate
c
TL207 Productivity C-mol yield
TL831 Productivity C-mol yield
Productivity (g/L/h)
C-mol yield (%)
d
Glucose
e
Xylose
f
Arabinose
g
Pyruvate
h
Acetate
i
Lactate
j
Residual glucose
k
Residual xylose
l
Residual arabinose
Muconate (g/L)
Glucose (g/L)
Xylose (g/L)
Arabinose (g/L)
Pyruvate (g/L)
Acetate (g/L)
Lactate (g/L)
Arabinose(g/L)
Time (hour)
TL207
TL831

**Figure S13 | Metabolic profiles of TL207 and TL831 in fed-batch bioreactor cultivation.** Fed-batch bioreactor evaluation of TL207 and TL831 at 0.5-L scale. Profiles show the bacterial growth ($OD_{600}$) (**a**), muconate production (**b**), and comparison of muconate productivity and carbon yield (**c**). Total sugar utilization profiles are presented for glucose (**d**), xylose (**e**), and arabinose (**f**). Accumulation of overflow metabolism intermediates is shown for pyruvate (**g**), acetate (**h**), while lactate (**i**) was not detected. Residual sugar concentrations in the bioreactor are provided for glucose (**j**), xylose (**k**), and arabinose (**l**). Rate: titer/time (g/L/h), Yield (C-mol %): [(mM muconate × 6) / mM (glucose × 6 + mM xylose × 5 + mM arabinose × 5) × 100%]. Data represent the average of $n = 2$ biological replicates. Error bars correspond to absolute error between duplicates. Numerical data are provided in a **Source Data File**.

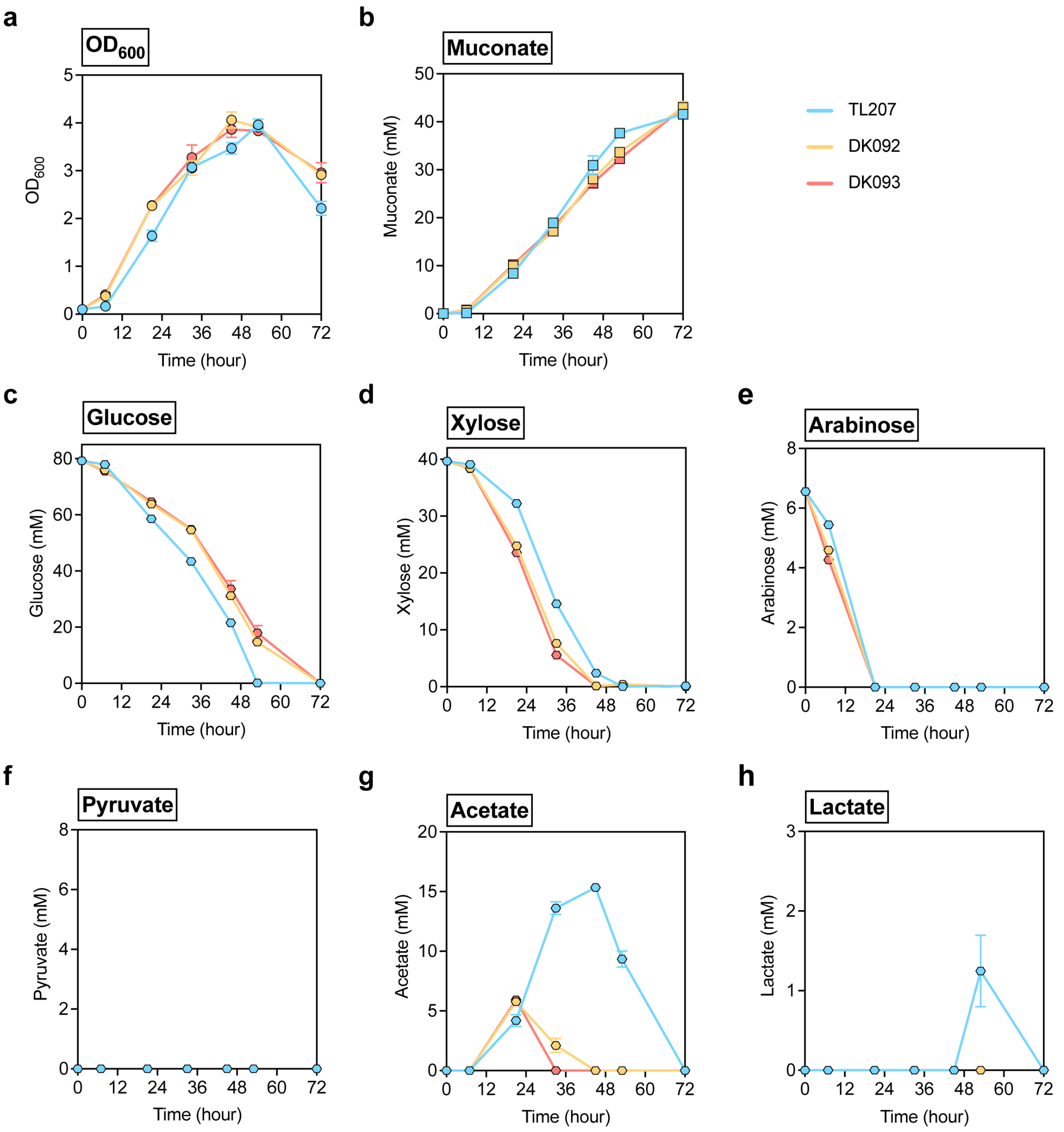


**Figure S14 | Metabolic profiles of TL207, DK092, and DK093 in shake-flask cultivation.** Metabolic profiles of TL207, DK092 (TL207 $P_{lac}$-*glf*), and DK093 (TL207 $P_{6.6}$-*glf*) in shake-flask cultivation for comparison (promoter strength: $P_{lac}$ < $P_{6.6}$ (*12*)). Profiles show the bacterial growth ($OD_{600}$) (**a**), muconate production (**b**), residual concentrations of glucose (**c**), xylose (**d**), and arabinose (**e**), and the accumulation of pyruvate (**f**), acetate (**g**), and lactate (**h**) on M9 medium supplemented with 25 g/L mock hydrolysate. Data represent the average of *n* = 3 biological replicates. Error bars correspond to standard deviation. Numerical data are provided in a **Source Data File**.

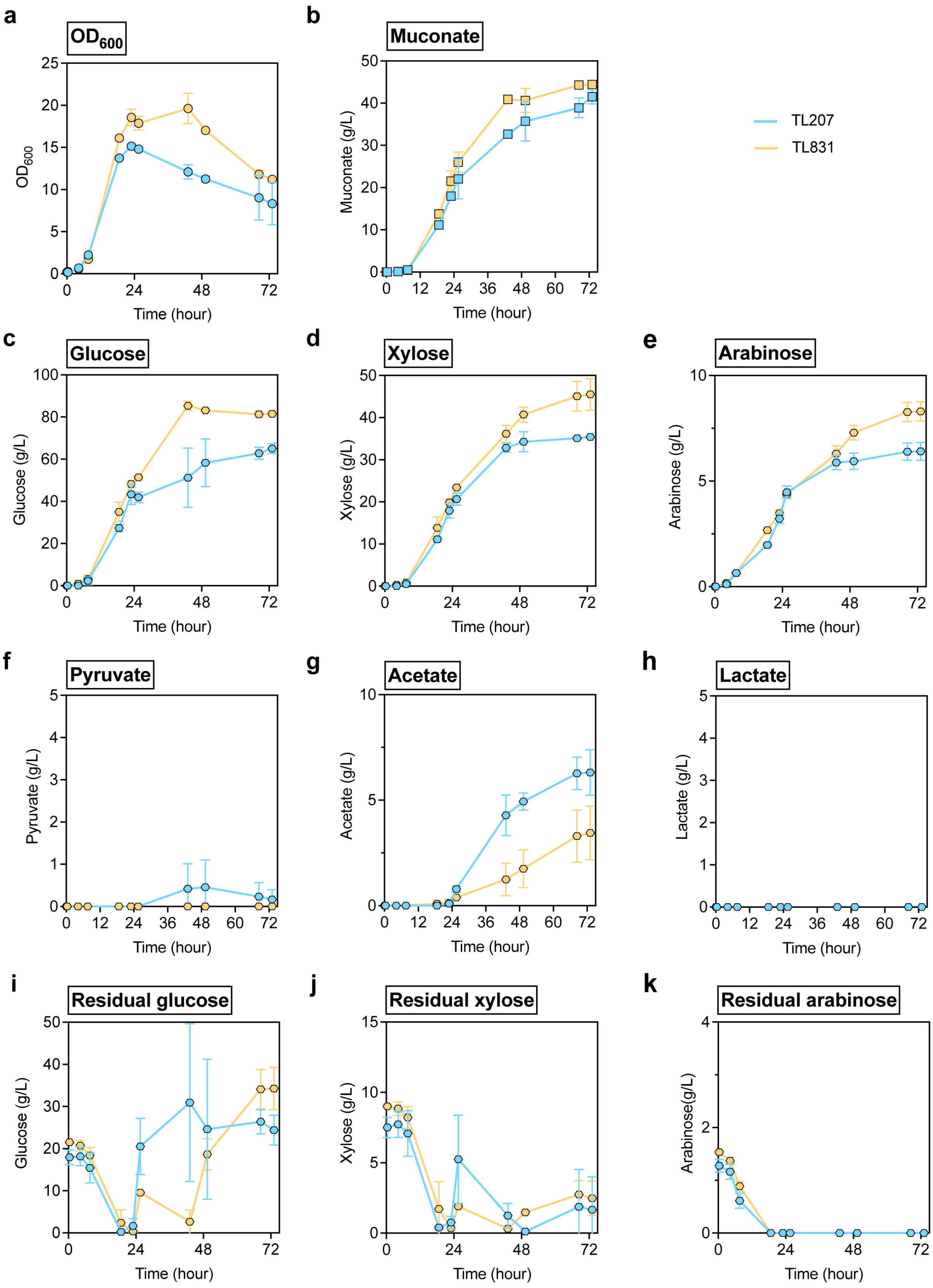

a
OD600
OD600
Time (hour)
b
Muconate
Muconate (g/L)
Time (hour)
TL207
TL831
c
Glucose
Glucose (g/L)
Time (hour)
d
Xylose
Xylose (g/L)
Time (hour)
e
Arabinose
Arabinose (g/L)
Time (hour)
f
Pyruvate
Pyruvate (g/L)
Time (hour)
g
Acetate
Acetate (g/L)
Time (hour)
h
Lactate
Lactate (g/L)
Time (hour)
i
Residual glucose
Glucose (g/L)
Time (hour)
j
Residual xylose
Xylose (g/L)
Time (hour)
k
Residual arabinose
Arabinose(g/L)
Time (hour)

**Figure S15 | Metabolic profiles of TL207 and TL565 in fed-batch bioreactor cultivation.** Fed-batch bioreactor evaluation of TL207 and TL565 at 0.5-L scale. Profiles show the bacterial growth ($OD_{600}$) (**a**) and muconate production (**b**). Total sugar utilization profiles are presented for glucose (**c**), xylose (**d**), and arabinose (**e**). Accumulation of overflow metabolism intermediates is shown for pyruvate (**f**), acetate (**g**), while lactate (**h**) was not detected. Residual sugar concentrations in the bioreactor are provided for glucose (**i**), xylose (**j**), and arabinose (**k**). Rate: titer/time (g/L/h), Yield (C-mol %): [(mM muconate × 6) / mM (glucose × 6 + mM xylose × 5 + mM arabinose × 5) × 100%]. Data represent the average of $n = 2$ biological replicates. Error bars correspond to absolute error between duplicates. Numerical data are provided in a **Source Data File**.

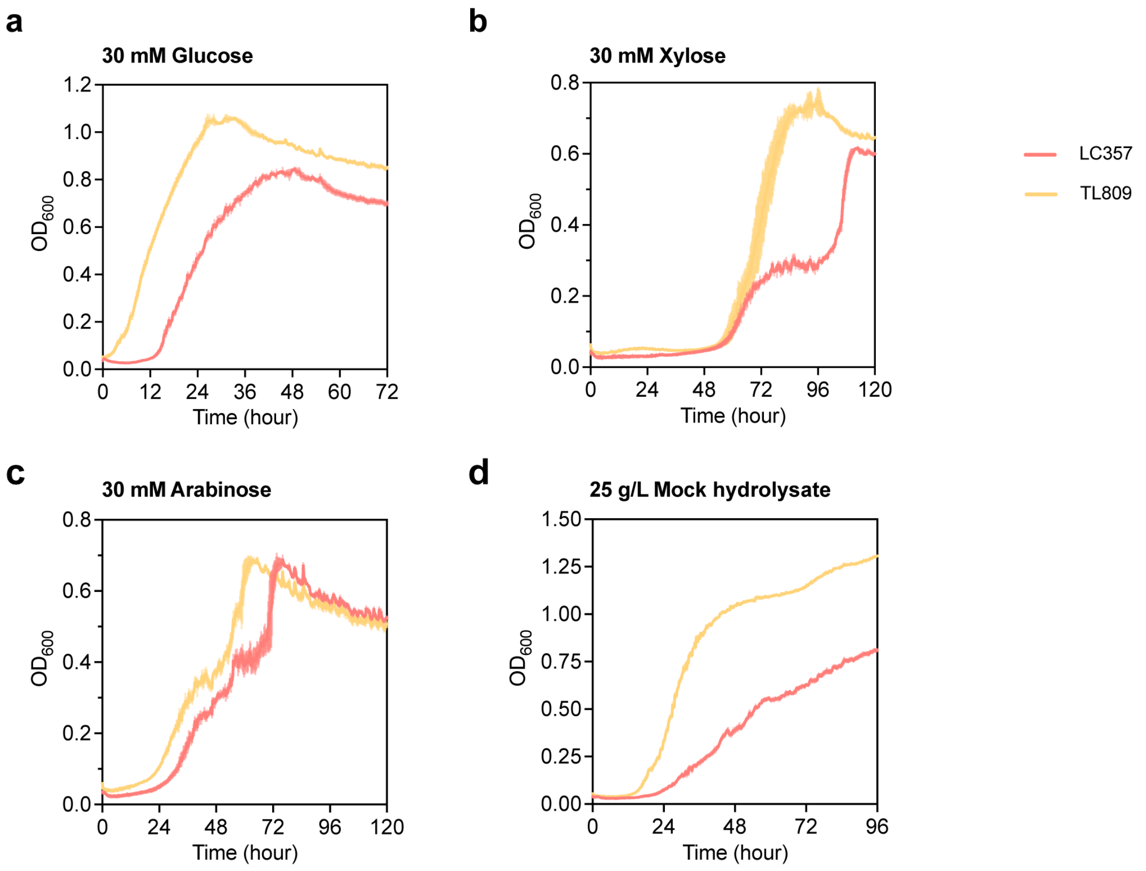


**Figure S16 | Growth curves of LC357 and TL809 in plate cultivation.** Growth curves of LC357 and TL809 in plate cultivation for comparison. Strains were grown on M9 medium supplemented with 30 mM glucose (**a**), 30 mM xylose (**b**), 30 mM arabinose (**c**), and 25 g/L mock hydrolysate (**d**). Data represent the average of $n = 3$ biological replicates except TL809 in **b** ($n = 2$). Error bars correspond to standard deviation except TL809 in **b**; Error bars correspond to absolute error between duplicates. Numerical data are provided in a **Source Data File**.

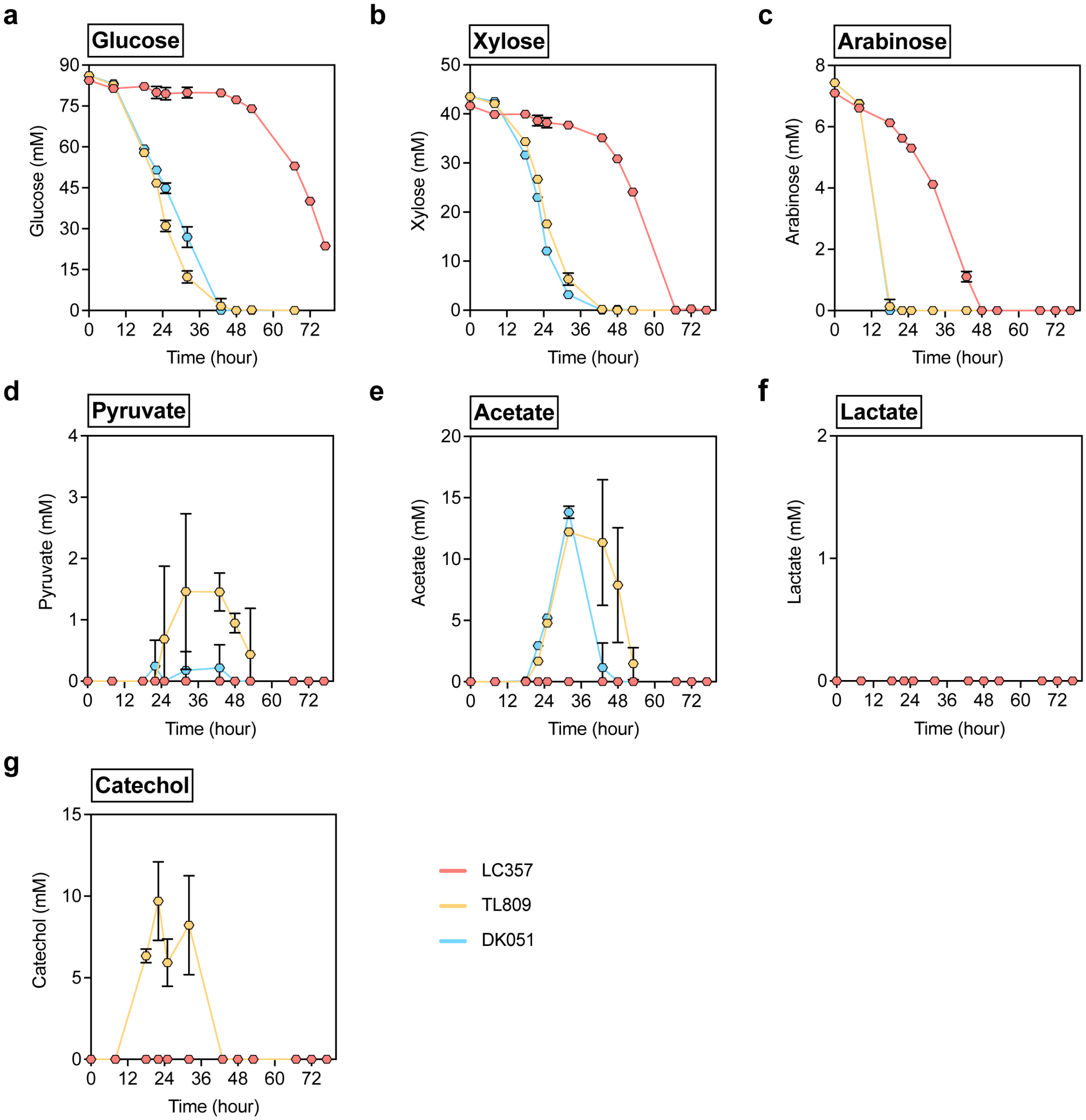


**Figure S17 | Metabolic profiles of LC357, TL809, and DK051 in shake-flask cultivation.** Metabolic profiles of LC357, TL809, and DK051 in shake-flask cultivation for comparison. Profiles show residual concentrations of glucose (**a**), xylose (**b**), and arabinose (**c**), and the accumulation of pyruvate (**d**), acetate (**e**), lactate (**f**), and catechol (**g**) on M9 medium supplemented with 25 g/L mock hydrolysate. Data represent the average of $n = 3$ biological replicates. Error bars correspond to standard deviation. Numerical data are provided in a **Source Data File**.

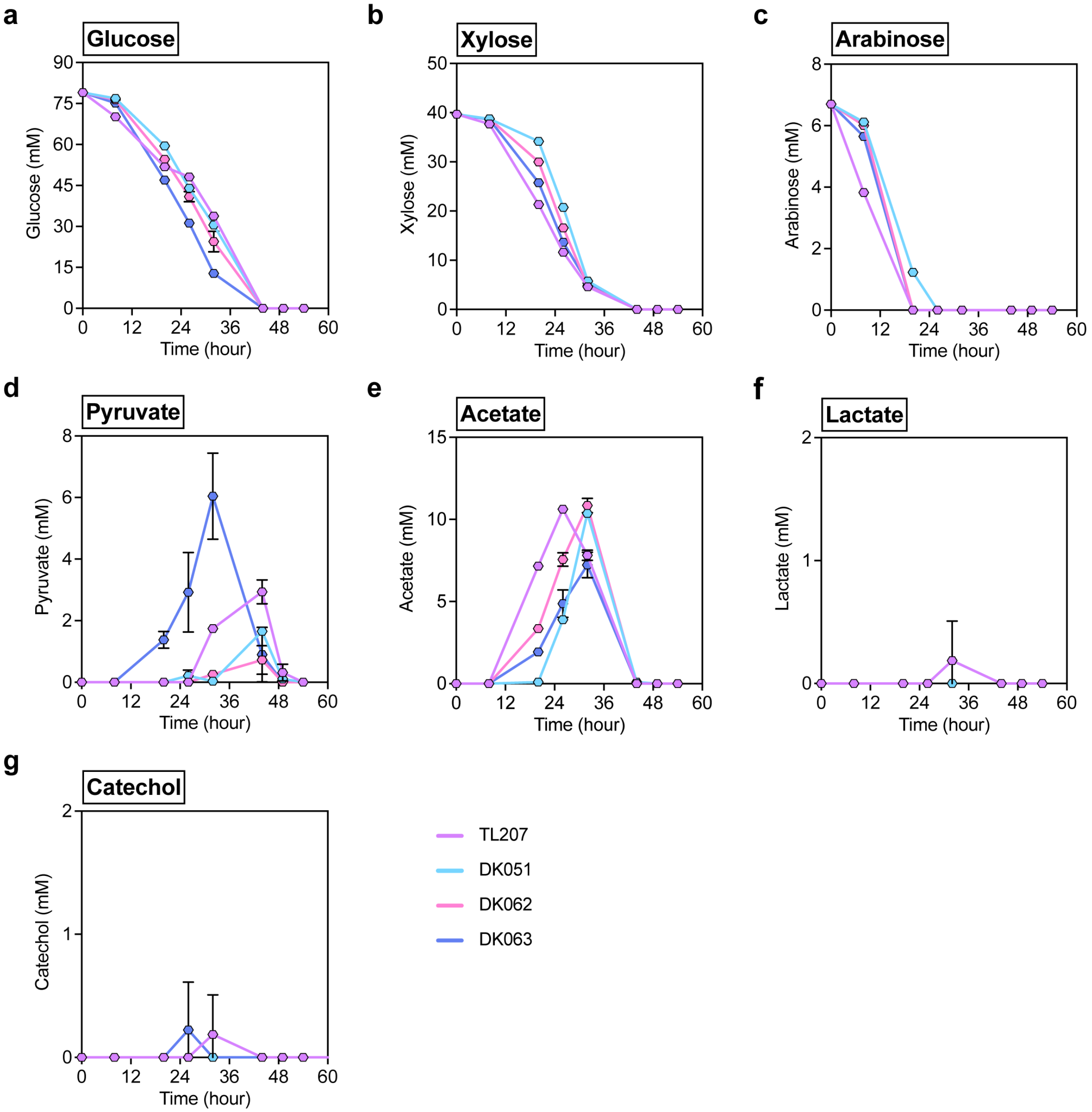


**Figure S18 | Metabolic profiles of TL207, DK051, DK062, and DK063 in shake-flask cultivation.** Metabolic profiles of TL207, DK051, DK062, and DK063 in shake-flask cultivation for comparison. Profiles show residual concentrations of glucose (**a**), xylose (**b**), and arabinose (**c**), and the accumulation of pyruvate (**d**), acetate (**e**), lactate (**f**), and catechol (**g**) on M9 medium supplemented with 25 g/L mock hydrolysate. Data represent the average of $n = 3$ biological replicates. Error bars correspond to standard deviation. Numerical data are provided in a **Source Data File**.

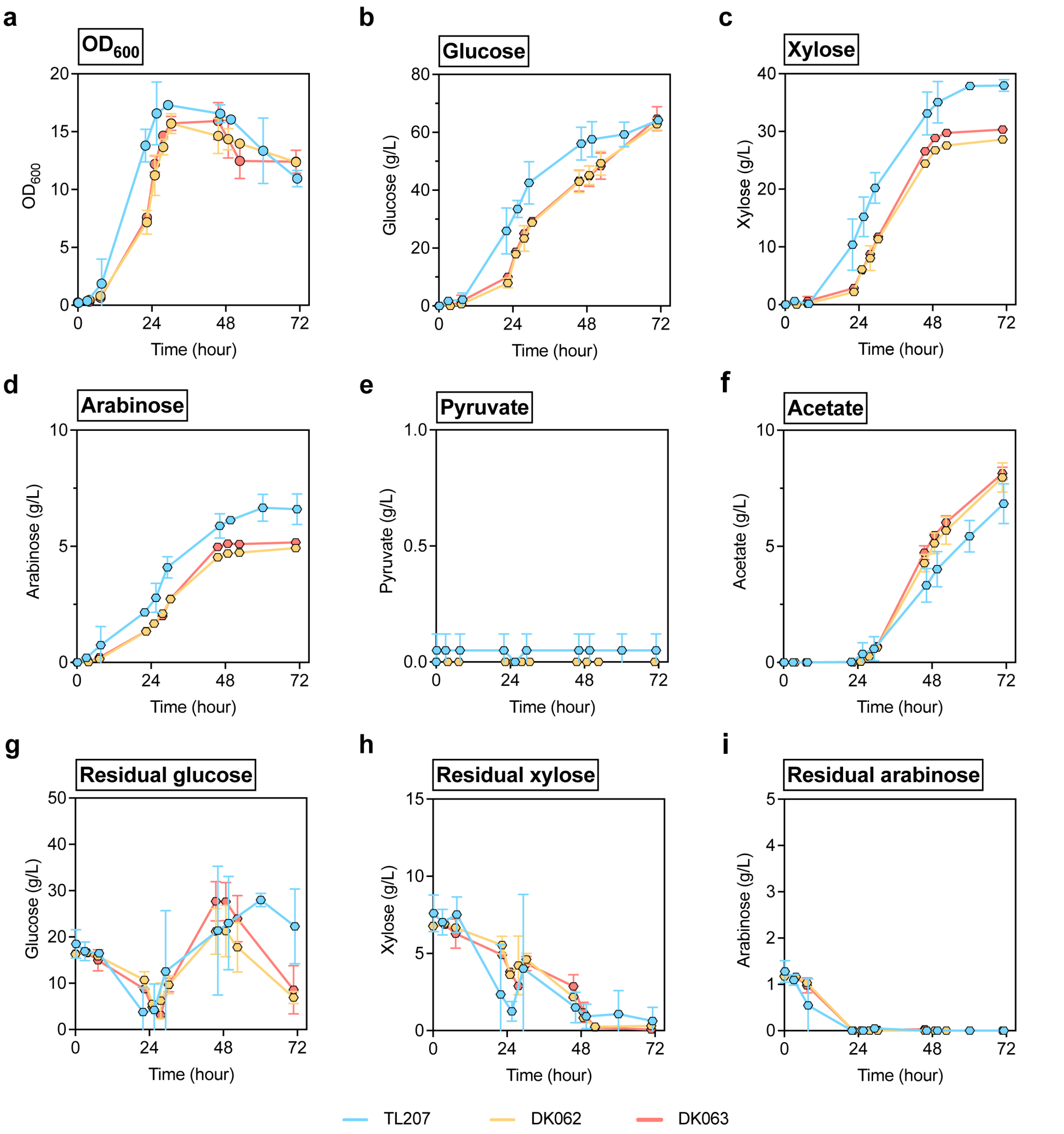


**Figure S19 | Metabolic profiles of TL207, DK062, and DK063 in fed-batch bioreactor cultivation.** Fed-batch bioreactor evaluation of TL207, DK062, and DK063 at 0.5-L scale. Profiles show the bacterial growth ($OD_{600}$) (**a**). Total sugar utilization profiles are presented for glucose (**b**), xylose (**c**), and arabinose (**d**). Accumulation of overflow metabolism intermediates is shown for pyruvate (**e**) and acetate (**f**), while lactate was not detected. Residual sugar concentrations in the bioreactor are provided for glucose (**g**), xylose (**h**), and arabinose (**i**). Data represent the average of $n = 2$ biological replicates. Error bars correspond to absolute error between duplicates. Numerical data are provided in a **Source Data File**.

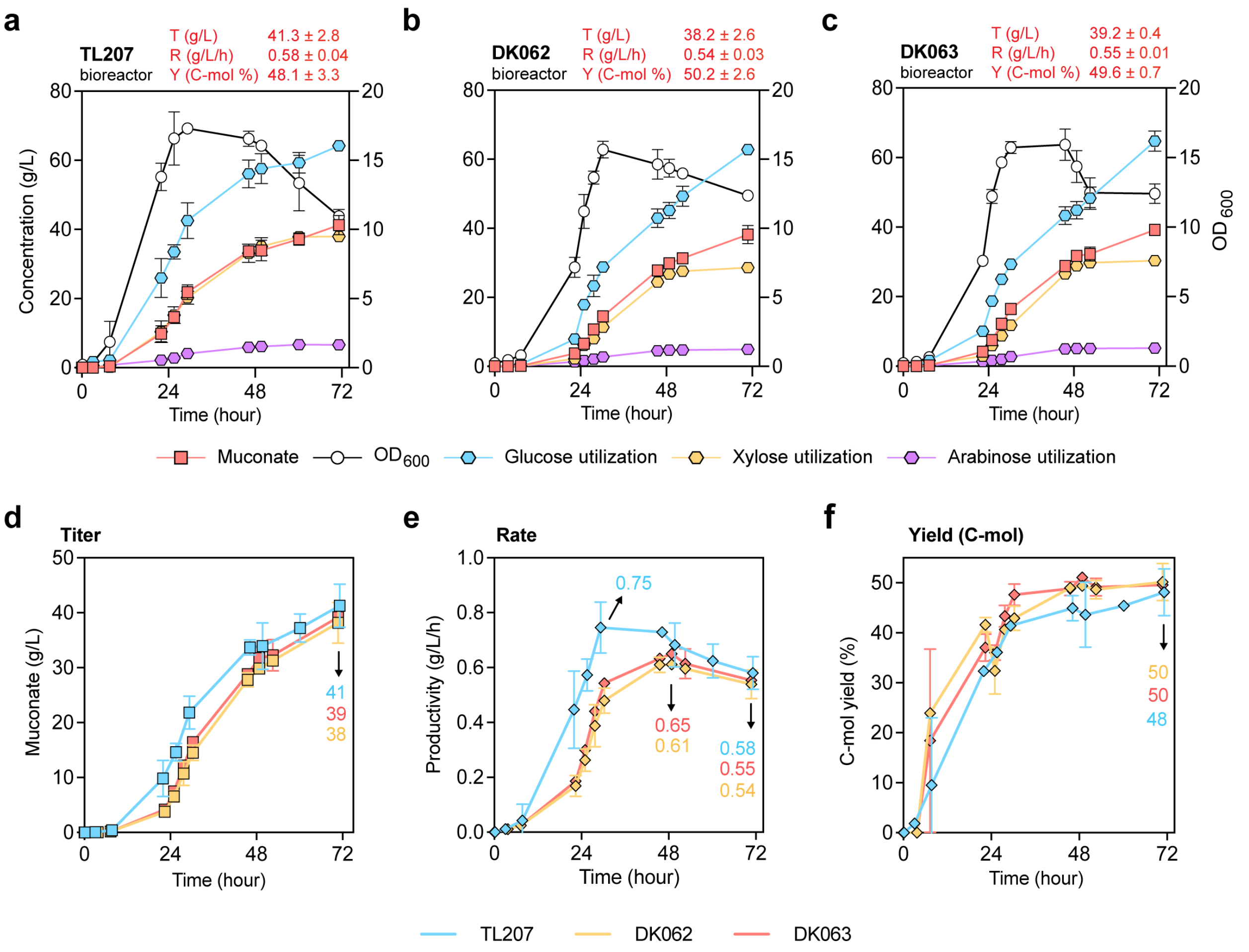


**Figure S20 | Strain performance comparison of TL207, DK062, and DK063 in fed-batch bioreactor cultivation.** **a**-**c** Evaluation of TL207 (**a**), DK062 (**b**), and DK063 (**c**) in 0.5-L bioreactors in fed-batch model. Cultivation profiles show bacterial growth ($OD_{600}$), total sugar utilization, and muconate production. **a**-**b** are adapted from **Fig. 5** for comparison. **d**-**f** Comparison of muconate titer (**d**), rate (**e**), and carbon yield (**f**) of TL207, DK062, and DK063 with their final titers, maximum and final productivities, and final C-mol yields presented. Titer: muconate concentration (g/L), Rate: titer/time (g/L/h), Yield (C-mol %): [(mM muconate × 6) / mM (glucose × 6 + mM xylose × 5 + mM arabinose × 5) × 100%]. Data represent the average of $n$ = 2 biological replicates. Error bars correspond to absolute error between duplicates. Numerical data are provided in a **Source Data File**.

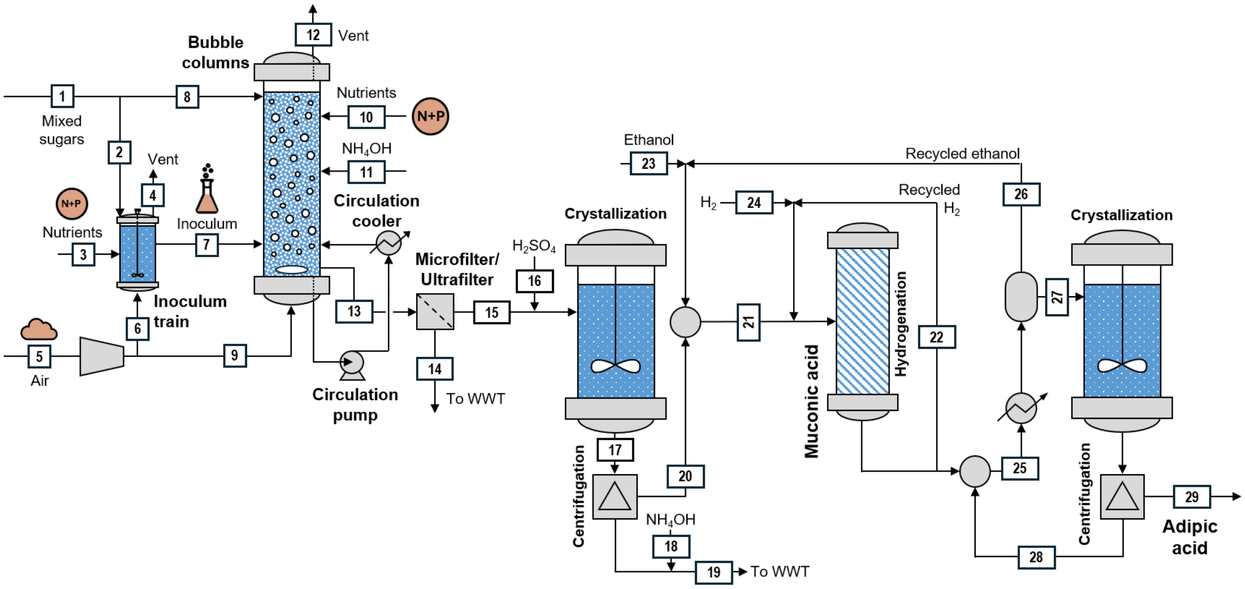


**Figure S21 | Main operations required for the bioconversion of mixed sugars to muconic acid and posterior catalytic upgrading to adipic acid.** G Detailed stream information for the process is shown in **Table S7**.

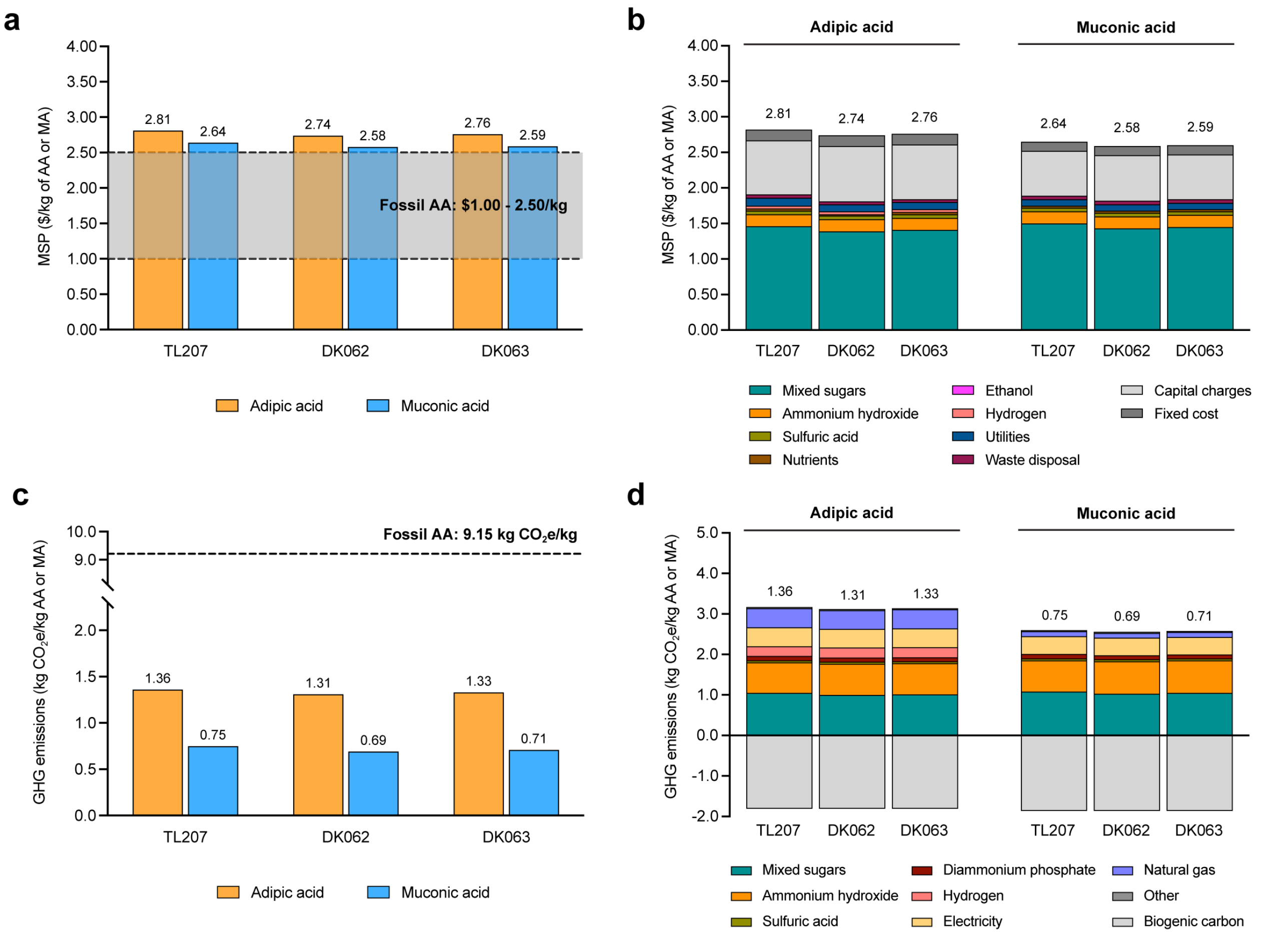


**Figure S22 | Expanded techno-economic analysis and life cycle assessment for adipic acid (AA) and muconic acid (MA) production. a** Minimum selling prices (MSPs) for AA (orange) and MA (blue) produced from mixed sugars using bioprocess metrics from TL207, DK062, and DK063. Total MSP are compared against the fossil-derived AA price range ($1.00 - $2.50/kg). Bioprocess metrics are based on the titer, rate, and yield achieved at the final timepoint of the bioreactor cultivations. **b** Detailed MSP cost breakdown into key contributors, including mixed sugars, ammonium hydroxide, sulfuric acid, nutrients, ethanol, hydrogen, utilities, waste disposal, capital charges, and fixed cost, in AA and MA production. The sum of costs from all contributors is corresponding to the final MSP value in **a**. **c** Greenhouse gas (GHG) emissions for AA and MA for TL207, DK062, and DK063. Total GHG emissions are compared against the fossil-derived AA benchmark (9.15 $kgCO_2e/kg$). **d** Detailed GHG emissions breakdown into key contributors, including mixed sugars, ammonium hydroxide, sulfuric acid, diammonium phosphate, hydrogen, electricity, natural gas, biogenic carbon, and others in AA and MA production. Negative values represent the credit from biogenic carbon. The sum of GHG emissions from all contributors is corresponding to the final GHG emissions value in **c**. Numerical data are provided in a **Source Data File**.

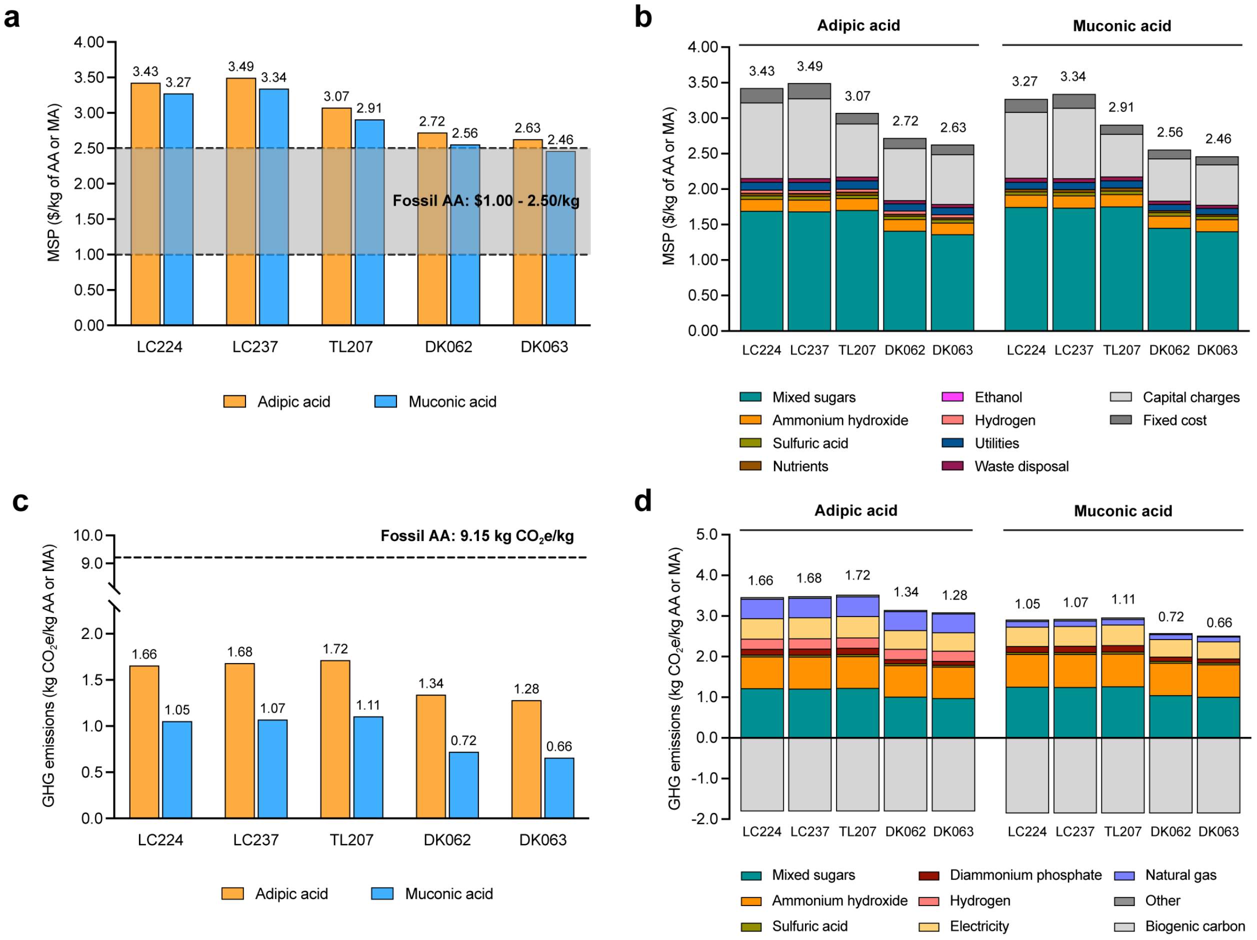


**Figure S23 | Techno-economic analysis and life cycle assessment for adipic acid (AA) and muconic acid (MA) production from TRY achieved at maximum rate. a** Minimum selling prices (MSPs) for AA (orange) and MA (blue) produced from mixed sugars using bioprocess metrics from a benchmark study by Ling *et al.* (2022) (LC224) (*2*), and those determined in this study (LC237, TL207, DK062, and DK063). Total MSP are compared against the fossil-derived AA price range ($1.00-$2.50/kg). Bioprocess metrics are based on the titer and yield achieved at the maximum rate of the bioreactor cultivations. **b** Detailed MSP cost breakdown into key contributors, including mixed sugars, ammonium hydroxide, sulfuric acid, nutrients, ethanol, hydrogen, utilities, waste disposal, capital charges, and fixed cost, in AA and MA production. The sum of costs from all contributors is corresponding to the final MSP value in **a**. **c** Greenhouse gas (GHG) emissions for AA and MA for the strains described in **a**. Total GHG emissions are compared against the fossil-derived AA benchmark (9.15 kg $CO_2$e/kg). **d** Detailed GHG emissions breakdown into key contributors, including mixed sugars, ammonium hydroxide, sulfuric acid, diammonium phosphate, hydrogen, electricity, natural gas, biogenic carbon, and others in AA and MA production. Negative values represent the credit from biogenic carbon. The sum of GHG emissions from all contributors is corresponding to the GHG emissions value in **c**. Numerical data are provided in a **Source Data File**.

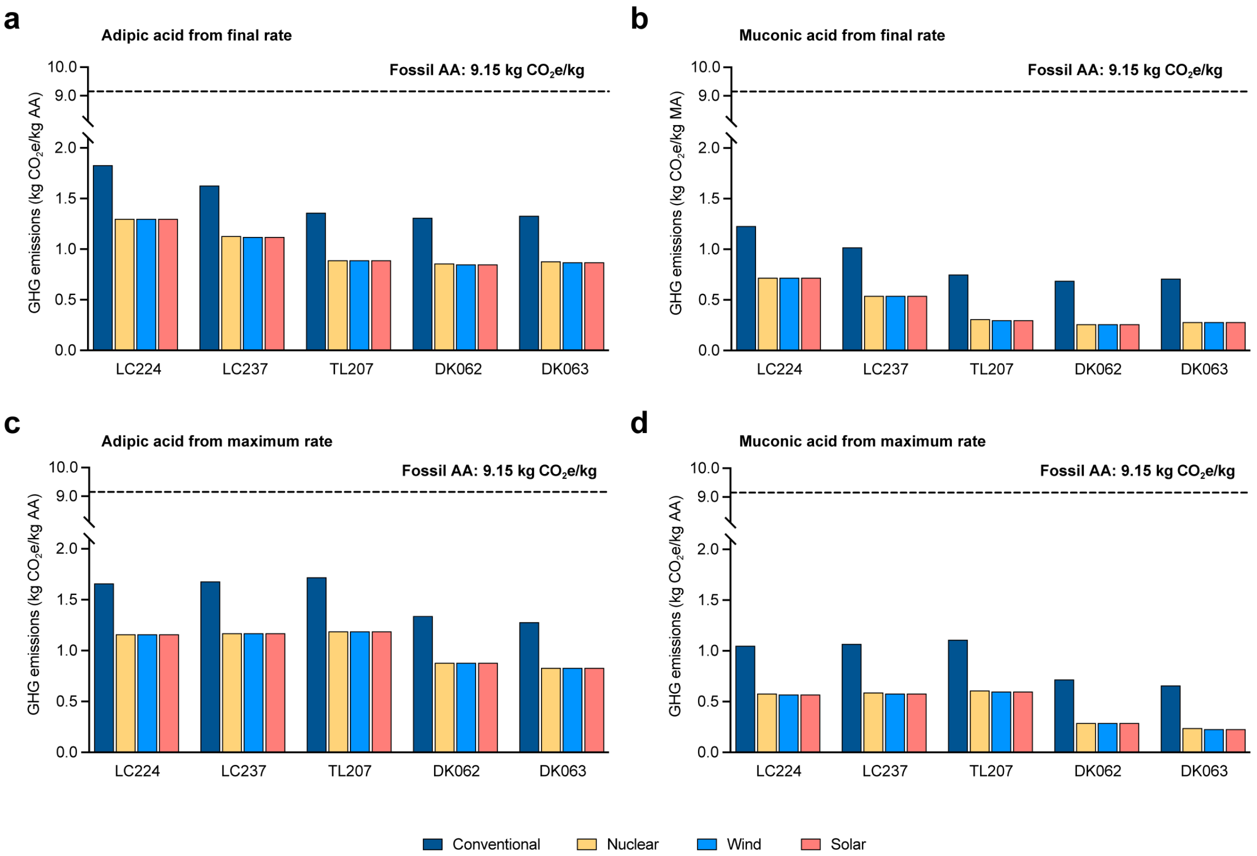


**Figure S24 | Sensitivity analysis of low-emission electricity sources on adipic acid (AA) and muconic acid (MA) production from DMR sugars.** Greenhouse gas (GHG) emissions were evaluated across different electricity sources – conventional, nuclear, wind and solar – to assess their impacts. Bioprocess metrics from a benchmark study by Ling *et al.* (2022) (LC224) (*2*) and those determined in this study (LC237, TL207, DK062, and DK063) were used to calculate GHG emissions. Total GHG emissions are compared against the fossil-derived AA benchmark (9.15 kg $CO_2$e/kg). The carbon intensity of electricity by source (g $CO_2$e/MJ) is as follows: conventional (110.21), nuclear (0.74), wind (0), and solar (0). **a**-**b** GHG emissions for AA (**a**) and MA (**b**) production. Bioprocess metrics are based on the titer, rate, and yield achieved at the final timepoint of the bioreactor cultivations. **c**-**d** GHG emissions for AA (**c**) and MA (**d**) production. Bioprocess metrics are based on the titer and yield achieved at the maximum rate of the bioreactor cultivations. Numerical data are provided in a **Source Data File**.

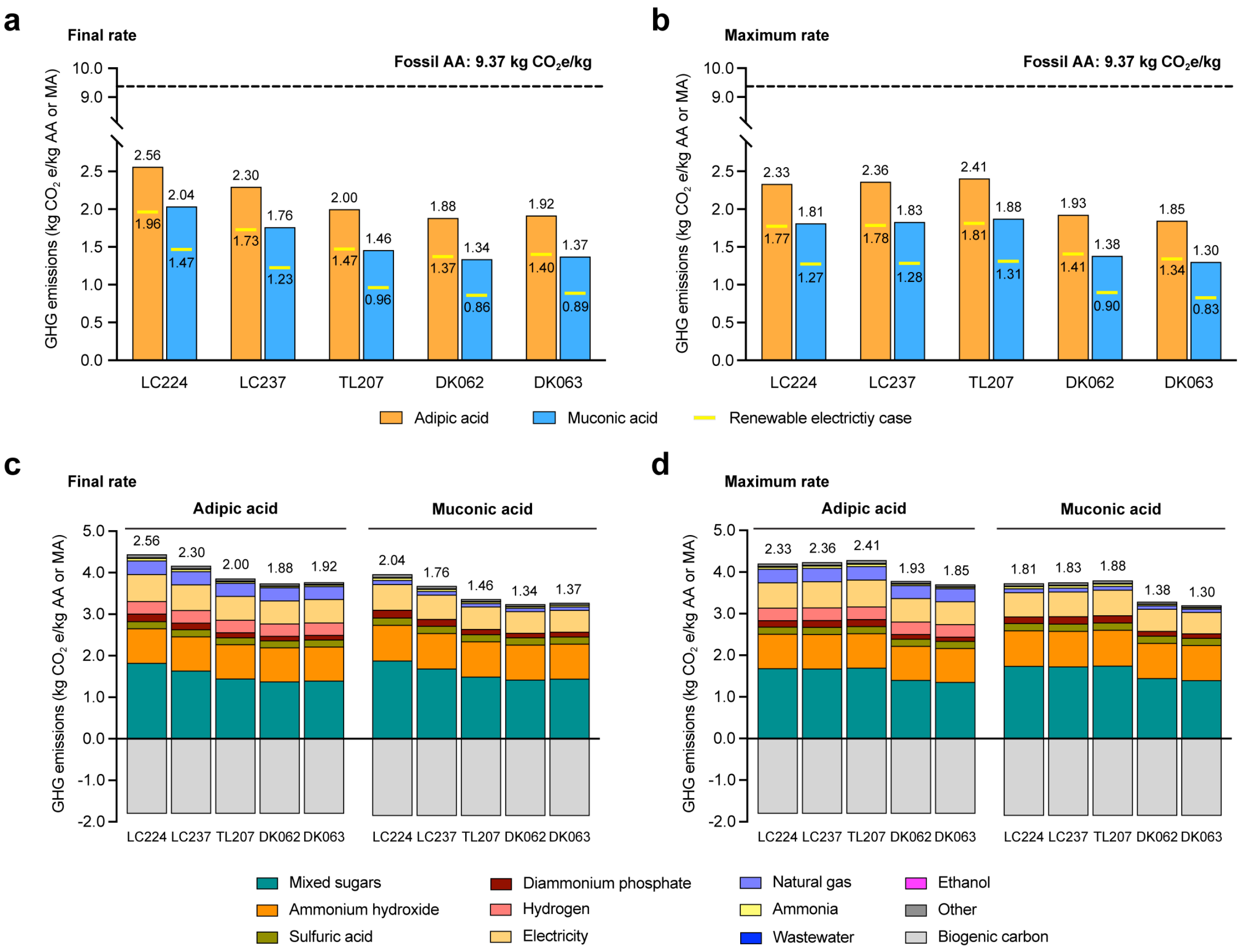


**Figure S25 | Life cycle assessment of adipic acid (AA) and muconic acid (MA) production from TRY achieved at final and maximum rate (Database: Brightway using ecoinvent v3.11). a-b** Greenhouse gas (GHG) emissions for AA (orange) and MA (blue) produced from mixed sugars using bioprocess metrics from a benchmark study by Ling *et al.* (2022) (LC224) (*2*), and those determined in this study (LC237, TL207, DK062, and DK063). Total GHG emissions are compared against the fossil-derived AA benchmark (9.37 kg $CO_2$e/kg). GHG emissions from renewable electricity case were highlighted as yellow lines. Bioprocess metrics are based on the titer, rate, and yield achieved at the final timepoint (**a**) and titer and yield achieved at the maximum rate of the bioreactor cultivations (**b**). **c-d** Detailed GHG emissions breakdown into key contributors, including mixed sugars, ammonium hydroxide, sulfuric acid, diammonium phosphate, hydrogen, electricity, natural gas, ammonia, wastewater, ethanol, biogenic carbon, and others in AA and MA production for the strains described in **a-b**. Negative values represent the credit from biogenic carbon. Bioprocess metrics are based on the titer, rate, and yield achieved at the final timepoint (**c**) and titer and yield achieved at the maximum rate of the bioreactor cultivations (**d**). The sum of GHG emissions from all contributors is corresponding to the final GHG emissions value in **a** and **b**, respectively. Numerical data are provided in a **Source Data File**.

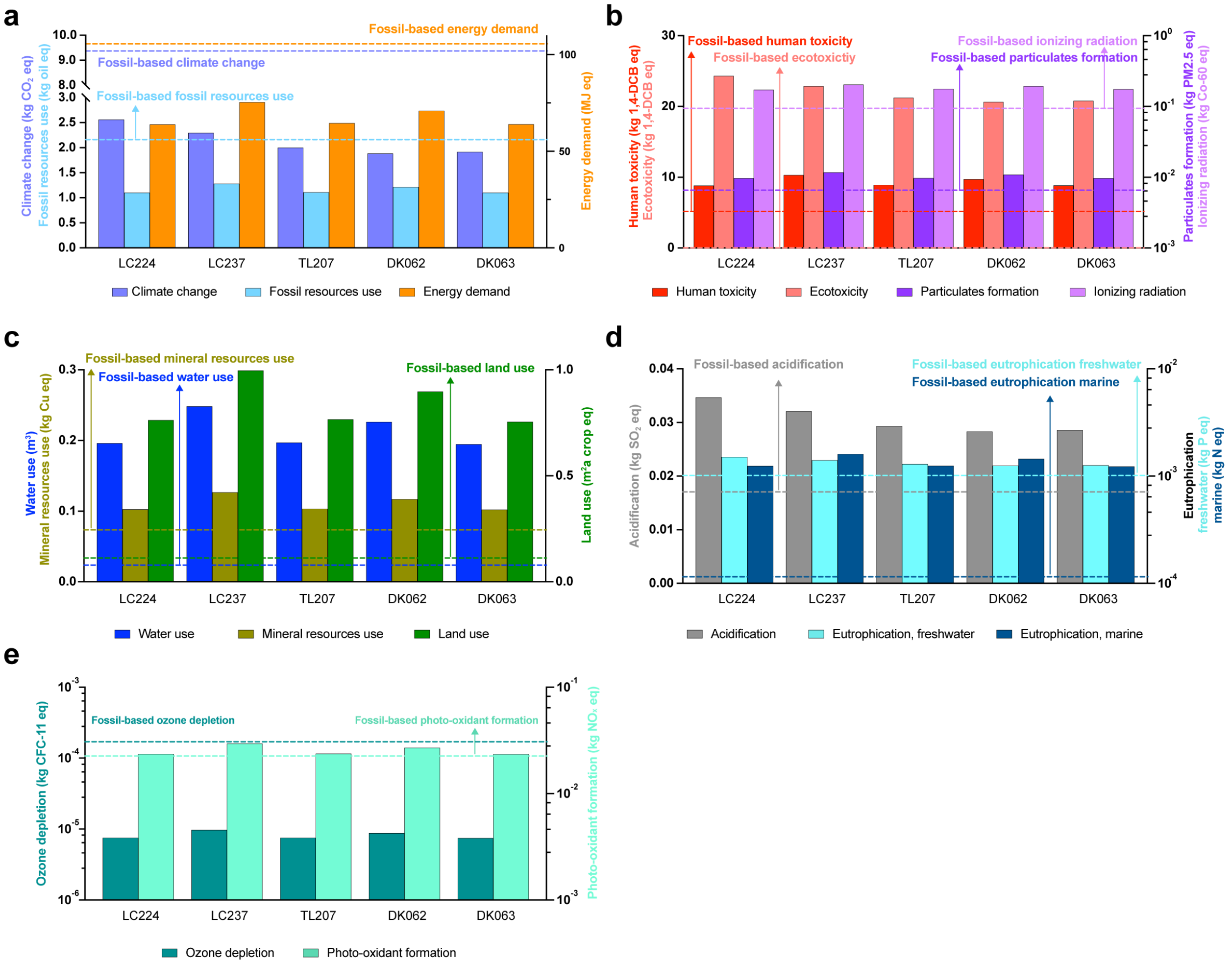


**Figure S26 | Life cycle assessment of adipic acid (AA) production across various environmental impact categories (Database: Brightway using ecoinvent v3.11).** Environmental impacts of AA production using engineered *P. putida* strains (LC237, TL207, DK062, and DK063) are compared against a benchmark study, Ling *et al.* (2022) (LC224) (*2*). Data are categorized into five major environmental themes. Impacts were calculated using bioprocess metrics (TRY) at the final timepoint. Dashed lines represent values from fossil-based adipic acid production. **a** Global warming and energy: climate change and fossil resources use (left Y-axis); energy demand (right Y-axis). **b** Toxicity and human health: human toxicity and ecotoxicity (left Y-axis); particulates formation and ionizing radiation (right Y-axis). **c** Resources and land use: water use and mineral resources use (left Y-axis); land use (right Y-axis). **d** Acidification and eutrophication: acidification (left Y-axis); eutrophication in freshwater and marine (right Y-axis). **e** Atmospheric impacts: ozone depletion (left Y-axis); photo-oxidant formation (right Y-axis). Numerical data are provided in a **Source Data File**.
