## Supplemental Table for "Pathway selection for arabinose utilization in *Pseudomonas putida* reveals a rate-yield tradeoff in muconic acid production from lignocellulosic sugars"

### Supplementary Table

**Table S1. Lists of abbreviation shown in Fig. 1.**

| Abbreviation | Full name |
|---|---|
| **GNCN** | Gluconate |
| **2-KGn** | 2-ketogluconate |
| **2-KG-6-P** | 2-ketogluconate-6-P |
| **L-KDA** | 2-keto-3-deoxy-L-arabinoate |
| **2-KGSA** | 2-ketoglutarate semialdehyde |
| **G6P** | glucose-6-P |
| **6PG** | 6-phosphogluconate |
| **KDPG** | 2-keto-3-deoxy-6-phosphogluconate |
| **G3P** | glyceraldehyde-3-P |
| **DHAP** | dihydroxyacetone phosphate |
| **FBP** | fructose-1,6-P2 |
| **F6P** | fructose-6-P |
| **E4P** | erythrose-4-phosphate |
| **S7P** | sedoheptulose-7-P |
| **R5P** | ribose-5-P |
| **Ri5P** | ribulose-5-P |
| **3PG** | 3-phosphoglycerate |
| **PEP** | phosphoenoylpyruvate |
| **DAHP** | 3-deoxy-D-arabino-heptulosonic acid 7-phosphate |
| **3DHQ** | 3-dehydroquinate acid |
| **3DHS** | 3-dehydroshikimate |
| **SA** | shikimate |
| **S3P** | shikimate-3-phosphate |
| **CSA** | chorismite |
| **4HB** | 4-hydroxybenzoate |
| **PCA** | protocatechuate |
| **CAT** | catechol |
| **ICIT** | isocitrate |
| **CIT** | citrate |
| **aKG** | alpha-ketoglutarate |
| **SUCC** | succinate |
| **FUM** | fumarate |
| **MAL** | malate |
| **GLX** | glyoxylate |
| **OAA** | oxaloacetate |
| **Ac-CoA** | acetyl-Coenzyme A |
| **PYR** | pyruvate |

**Table S2. Extended genotypes of strains used in this study.**

| Strains | Genotype | Sources |
|---|---|---|
| **LC224** | ***P. putida*** **KT2440** Δ*catRBC*::$P_{tac}$:*catA* Δ*pcaHG*::$P_{tac}$:*aroY*:*ecdB*:*asbF* Δ*pykA*::*aroG-D146N*:*aroY*:*ecdB*:*asbF* Δ*pykF* Δ*ppc* Δ*pgi-2* Δ*gcd* Δ*hexR* Δ*ampC*::$P_{xylE*}$:*xylE*:$P_{tac}$:*xylAB*:*talB*:*tktA* PP_1736-1737(intergenic)::$P_{lac}$:*ubiC*-C22 *xylE-A62V, A455V* $P_{PP\_2569}$ G→A Δ*pykF*::$P_{tac}$:*aroB* | (*1*) |
| **LC237** | **LC224** Δ*gcd*::$P_{xylE}$:$araE_1$:$P_{tac}$:$araC_2D_2A_2B_2E_2$ | This study |
| **LC357** | **LC224** Δ*gcd*::$P_{xylE}$:$araE_1$:$P_{tac}$:$araB_1A_1D_1$ | This study |
| **TL015** | **LC237** Δ*araE* | This study |
| **TL170** | **LC237** Δ*ppc*::$P_{lac}$:RBSv1-*glf* | This study |
| **TL172** | **LC237** Δ*ppc*::$P_{lac}$:RBSv2- *glf* | This study |
| **TL174** | **LC237** Δ*ppc*::$P_{lac}$:RBSv3- *glf* | This study |
| **TL207** | **LC237** Δ*ppc*::$P_{lac}$:RBSv1-*glf* Δ*catRBC*::$P_{tac}$:3,574 TIR RBS-*catA* | This study |
| **TL248** | **LC237** Δ*pgi-2*::$P_{tac}$:*catA2* | This study |
| **TL250** | **LC237** Δ*ppc*::$P_{lac}$:RBSv1-*glf* Δ*pgi-2*::$P_{tac}$:*catA2* | This study |
| **TL252** | **LC237** Δ*ppc*::$P_{lac}$:RBSv3- *glf* Δ*pgi-2*::$P_{tac}$:*catA2* | This study |
| **TL254** | **LC237** Δ*ppc*::$P_{lac}$:RBSv1-*glf* Δ*catRBC*::$P_{tac}$:3,574 TIR RBS-*catA* Δ*pgi-2*::$P_{tac}$:*catA2* | This study |
| **TL477** | **LC237** Δ*ppc*::$P_{lac}$:RBSv1-*glf* Δ*catRBC*::$P_{tac}$:3,574 TIR RBS-*catA* $P_{6.6}$:$araC_2D_2A_2B_2E_2$ *gltA*-R164L | This study |
| **TL565** | **LC237** Δ*ppc*::$P_{lac}$:RBSv1-*glf* Δ*catRBC*::$P_{tac}$:3,574 TIR RBS-*catA* Δ*xylX*::$P_{tac}$:*aroG* D146N | This study |
| **TL809** | **LC357** Δppc::$P_{lac}$:RBSv1-*glf* | This study |
| **TL831** | **LC237** Δ*ppc*::$P_{lac}$:RBSv1-*glf* Δ*catRBC*::$P_{tac}$:3,574 TIR RBS-*catA* $P_{6.6}$:$araC_2D_2A_2B_2E_2$ *gltA*-R164L | This study |
| **JE3288** | ***P. putida*** **KT2440** Δ*hsdR*::Bxb1int-*attB* Δ*gcd*::$P_{xylE}$-*araE*:$P_{tac}$-$araB_1A_1D_1$ Δ*ampC*::$P_{xylE*}$:*xylE*:$P_{tac}$:*xylAB*:*talB*:*tktA* | (*2*) |
| **TL683** | **JE3288** Δ*araE* | This study |
| **DK037** | **LC237** 2,424 TIR RBS *araE* | |
| **DK051** | **LC357** Δ*ppc*::$P_{lac}$:RBSv1-*glf* Δ*catRBC*::$P_{tac}$:RBS C>G-*catA* | This study |
| **DK062** | **LC357** Δ*ppc*::$P_{lac}$:RBSv1-*glf* Δ*catRBC*::$P_{tac}$:RBS C>G-*catA* Δ*xylX*::$P_{tac}$:*aroG*-D146N | This study |
| **DK063** | **LC357** Δ*ppc*::$P_{lac}$:RBSv1-*glf* Δ*catRBC*::$P_{tac}$:RBS C>G-*catA* Δ*xylX*::$P_{tac}$:*aroG*-D146N *gltA*-R164L | This study |
| **DK092** | **LC237** Δ*ppc*::$P_{lac}$:RBSv1-*glf* Δ*catRBC*::$P_{tac}$:3,574 TIR RBS-*catA* $P_{lac}$:8,332 TIR RBS-*glf* | This study |
| **DK093** | **LC237** Δ*ppc*::$P_{lac}$:RBSv1-*glf* Δ*catRBC*::$P_{tac}$:3,574 TIR RBS-*catA* $P_{3.3}$:8,332 TIR RBS-*glf* | This study |

**Table S3. Construction details for strains used in this study.**

| Strains | Genotype | Construction details |
|---|---|---|
| **LC237** | **LC224** Δ*gcd*::$P_{xylE}$:*araE*$_1$:$P_{tac}$:*araC*$_2$*D*$_2$*A*$_2$*B*$_2$*E*$_2$ | The arabinose ozidative pathway was integrated into the Δ*gcd* locus of strain **LC224** using plasmid pJE1345 via conjugation. |
| **LC357** | **LC224** Δ*gcd*::$P_{xylE}$:*araE*$_1$:$P_{tac}$:*araB*$_1$*A*$_1$*D*$_1$ | The arabinose isomerase pathway was integrated into the Δ*gcd* locus of strain **LC224** using plasmid pJE1176 via conjugation. |
| **TL015** | **LC237** Δ*araE* | The *araE* was deleted from **LC237** using plasmid pTL012 via conjugation. |
| **TL170** | **LC237** Δ*ppc*::$P_{lac}$:RBSv1-*glf* | The *glf* with RBSv1 was integrated into the Δ*ppc* locus of strain **LC237** using plasmid pTL039 via conjugation. |
| **TL172** | **LC237** Δ*ppc*::$P_{lac}$:RBSv2-*glf* | The *glf* with RBSv2 was integrated into the Δ*ppc* locus of strain **LC237** using plasmid pTL040 via conjugation. |
| **TL174** | **LC237** Δ*ppc*::$P_{lac}$:RBSv3-*glf* | The *glf* with RBSv3 was integrated into the Δ*ppc* locus of strain **LC237** using plasmid pTL041 via conjugation. |
| **TL207** | **LC237** Δ*ppc*::$P_{lac}$:RBSv1-*glf* Δ*catRBC*::$P_{tac}$:3,574 TIR RBS-*catA* | The native *catA* RBS was replaced with RBSv4 in strain **TL170** using plasmid pTL048 via conjugation. |
| **TL248** | **LC237** Δ*pgi-2*::$P_{tac}$:*catA2* | The *catA2* was integrated into the Δ*pgi-2* locus of strain **LC237** using plasmid pTL051 via conjugation. |
| **TL250** | **LC237** Δ*ppc*::$P_{lac}$:RBSv1-*glf* Δ*pgi-2*::$P_{tac}$:*catA2* | The *catA2* was integrated into the Δ*pgi-2* locus of strain **TL170** using plasmid pTL051 via conjugation. |
| **TL252** | **LC237** Δ*ppc*::$P_{lac}$:RBSv3-*glf* Δ*pgi-2*::$P_{tac}$:*catA2* | The *catA2* was integrated into the Δ*pgi-2* locus of strain **TL174** using plasmid pTL051 via conjugation. |
| **TL254** | **LC237** Δ*ppc*::$P_{lac}$:RBSv1-*glf* Δ*catRBC*::$P_{tac}$:3,574 TIR RBS-*catA* Δ*pgi-2*::$P_{tac}$:*catA2* | The *catA2* was integrated into the Δ*pgi-2* locus of strain **TL207** using plasmid pTL051 via conjugation. |
| **TL477** | **LC237** Δ*ppc*::$P_{lac}$:RBSv1-*glf* Δ*catRBC*::$P_{tac}$:3,574 TIR RBS-*catA* Δ$P_{tac}$::$P_{6.6}$:*araC*$_2$*D*$_2$*A*$_2$*B*$_2$*E*$_2$ | The $P_{tac}$ promoter was replaced with the $P_{6.6}$ promoter (*3*) in the *araC*$_2$*D*$_2$*A*$_2$*B*$_2$*E*$_2$ operon in strain **TL207** using plasmid pTL104 via electroporation. |
| **TL565** | **LC237** Δ*ppc*::$P_{lac}$:RBSv1-*glf* Δ*catRBC*::$P_{tac}$:3,574 TIR RBS-*catA* Δ*xylX*::$P_{tac}$:*aroG* D146N | The second copy of $P_{tac}$:*aroG*-D146N was integrated into strain **TL207** using plasmid pTL084 via electroporation. |
| **TL683** | **JE3288** Δ*araE* | The arabinose transporter was deleted from **JE3288** using plasmid pTL012 via electroporation. |
| **TL809** | **LC357** Δ*ppc*::$P_{lac}$:RBSv1-*glf* | The *glf* with RBSv1 was integrated into the Δ*ppc* locus of strain **TL794** using plasmid pTL039 via electroporation. |
| **TL831** | **LC237** Δ*ppc*::$P_{lac}$:RBSv1-*glf* Δ*catRBC*::$P_{tac}$:3,574 TIR RBS-*catA* Δ$P_{tac}$::$P_{6.6}$:*araC*$_2$*D*$_2$*A*$_2$*B*$_2$*E*$_2$ *gltA*-R164L | The GltA-R164L mutation was introduced in strain **TL477** using plasmid pTL158 via electroporation. |
| **DK037** | **LC237** 2,424 TIR RBS *araE* | The 2,424 TIR RBS of *araE* mutation was introduced in strain **LC237** using plasmid pDK066 via electroporation. |
| **DK051** | **LC357** Δ*ppc*::$P_{lac}$:RBSv1-*glf* Δ*catRBC*::$P_{tac}$:RBS C>G-*catA* | The cassette of $P_{tac}$:RBS C>G *catA* was integrated into strain **TL809** using plasmid pTL131 via electroporation. |
| **DK062** | **LC357** Δ*ppc*::$P_{lac}$:RBSv1-*glf* Δ*catRBC*::$P_{tac}$:RBS C>G-*catA* Δ*xylX*::$P_{tac}$:*aroG* D146N | The second copy of $P_{tac}$:*aroG*-D146N was integrated into strain **DK051** using plasmid pTL084 via electroporation. |
| **DK063** | **LC357** Δ*ppc*::$P_{lac}$:RBSv1-*glf* Δ*catRBC*::$P_{tac}$:RBS C>G-*catA* Δ*xylX*::$P_{tac}$:*aroG* D146N *gltA* R164L | The GltA-R164L mutation was introduced in strain **DK062** using plasmid pTL158 via electroporation. |
| **DK092** | **LC237** Δ*ppc*::$P_{lac}$:RBSv1-*glf* Δ*catRBC*::$P_{tac}$:3,574 TIR RBS-*catA* $P_{lac}$:8,332 TIR RBS-*glf* | The 8,332 TIR RBS of *glf* mutation was introduced in strain **TL207** using plasmid pDK057 via electroporation. |
| **DK093** | **LC237** Δ*ppc*::$P_{lac}$:RBSv1-*glf* Δ*catRBC*::$P_{tac}$:3,574 TIR RBS-*catA* $P_{3.3}$:8,332 TIR RBS-*glf* | The $P_{lac}$ promoter was replaced with the $P_{3.3}$ promoter (*3*) in the *glf* expression cassette in strain **DK092** using plasmid pDK055 via electroporation. |

**Table S4. Plasmids used in this study.**

| Plasmids | Description | Source |
|---|---|---|
| **pJE1345** | Insertion of arabinose oxidative pathway consisting of the codon-optimized AraE transporter from *E. coli* and the codon-optimized $araB_2C_2D_2A_2B_2E_2$ genes from *Burkholderia ambifaria* AMMD driven by the $P_{tac}$ promoter | (*2*) |
| **pJE1176** | Insertion of the arabinose isomerase pathway consisting of the codon-optimized AraE transporter from *E. coli* and the codon-optimized $araB_1A_1D_1$ genes from *E. coli* driven by the $P_{tac}$ promoter | (*2*) |
| **pK18msB** | Sucrose counter-selection allelic exchange vector for *P. putida* KT2440; $Km^R$ | (*1*) |
| **pK18msBI** | Sucrose counter-selection allelic exchange vector for *P. putida* KT2440; $Km^R$ ;LacI | This study |
| **pTL012** | pK18msBI-derived plasmid for deletion of *araE* in arabinose-oxidative strain | This study |
| **pTL039** | pK18msBI-derived plasmid for insertion of the codon-optimized *glf* driven by $P_{lac}$ promoter to the Δ*ppc* locus with RBSv1 | This study |
| **pTL040** | pK18msBI-derived plasmid for insertion of the codon-optimized *glf* driven by $P_{lac}$ promoter to the Δ*ppc* locus with RBSv2 | This study |
| **pTL041** | pK18msBI-derived plasmid for insertion of the codon-optimized *glf* driven by $P_{lac}$ promoter to the Δ*ppc* locus with RBSv3 | This study |
| **pTL048** | pK18msBI-derived plasmid for replacement of native *catA* RBS with RBSv4 | This study |
| **pTL051** | pK18msBI-derived plasmid for insertion of *catA2* driven by $P_{tac}$ promoter to the Δ*pgi-2* locus | This study |
| **pTL084** | pK18msBI-derived plasmid for introduction of the codon-optimized $P_{tac}$:*aroG-D146N* to the Δ*xylX* locus | This study |
| **pTL104** | pK18msBI-derived plasmid for replacement of $P_{tac}$ with $P_{6.6}$ to drive $araC_2D_2A_2B_2E_2$ expression | This study |
| **pTL158** | pK18msBI-derived plasmid for introduction of *gltA-R164L* mutation | This study |
| **pDK055** | pK18msBI-derived plasmid for introduction of 2,424 TIR RBS to drive *araE* expression | This study |
| **pDK057** | pK18msBI-derived plasmid for introduction of 8,332 TIR RBS to drive *glf* expression | This study |
| **pDK066** | pK18msBI-derived plasmid for replacement of $P_{lac}$ with $P_{3.3}$ to drive *glf* expression | This study |

**Table S5. Different sequences of each RBS and their predicted TIR of the chemically synthesized oligonucleotide. TIRs are predicted by the Salis RBS Calculator version 2.0 (*4*).**

| RBS Identifier | Sequence (5'-3') | Predicted TIR | Details |
|---|---|---|---|
| **RBSv1** | tgtgtggaagcatgaagacaacttaagcataagga gatttctt | 25419 | Strong RBS for *glf* |
| **RBSv2** | attcagccattcctgttcgctgggagcggattcagg tttttcc | 74 | Weak RBS for *glf* |
| **RBSv3** | attcagccattcaggaaccatacgcacgaggaggt aaagtatt | 618 | Native RBS for *glf* |
| **3,574 RBS *catA*** | tcaggacagaaggaaggtattta | 3574 | Modified RBS for *catA* |
| **2,424 RBS *araE*** | ctttaacatattaagaaggtctaact | 2424 | Modified RBS for *araE* |
| **8,332 RBS *glf*** | tcaatacattgacccaggaatcaca | 8332 | Medium RBS for *glf* |

**Table S6. Main reactions considered in the bioconversion of glucose, xylose, and arabinose to muconic acid and *P. putida* biomass.**

| Reaction | Equation |
|---|---|
| Glucose to product | **1** Glu + **1.94** $O_2$ → **0.74** Muconic + **1.57** $CO_2$ + **3.78** $H_2O$ |
| Xylose/arabinose to product | **1** Xyl/Ara + **1.57** $O_2$ → **0.62** Muconic + **1.26** $CO_2$ + **3.13** $H_2O$ |
| Glucose to biomass | **1** Glu + **0.28** $NH_3$ + **1.17** $O_2$ → **4.8** *P. putida* + **1.2** $CO_2$ + **1.98** $H_2O$ |
| Xylose/arabinose to biomass | **1** Xyl/Ara + **0.23** $NH_3$ + **0.98** $O_2$ → **4** *P. putida* + **1** $CO_2$ + **1.65** $H_2O$ |
| Glucose loss to contamination | **1** Glu + **6** $O_2$ → **6** $CO_2$ + **6** $H_2O$ |
| Xylose/arabinose loss to contamination | **1** Xyl/Ara + **5** $O_2$ → **5** $CO_2$ + **5** $H_2O$ |

**Table S7. Detailed stream information for the process shown in Figure S21**. The process corresponds to the production of adipic acid following the TRY conditions obtained for the DK062 strain (MSP of $2.74/kg).

| | Units | Stream number | | | | | | | |
|---|---|---|---|---|---|---|---|---|---|
| | | 1 | 2 | 3 | 4 | 5 | 6 | 7 | 8 |
| Temperature | **°C** | **38.0** | **38.0** | **25.0** | **32.0** | **25.0** | **40.0** | **32.0** | **38.0** |
| Pressure | atm | **1.0** | **1.0** | **1.0** | **1.0** | **1.0** | **5.9** | **1.0** | **1.0** |
| Mass Flows | kg/h | **419,501** | **41,950** | **287** | **4,221** | **65,409** | **3,874** | **41,706** | **377,551** |
| Water | kg/h | 373,798 | 37,380 | 0 | 117 | 1,281 | 76 | 38,082 | 336,418 |
| Ethanol | kg/h | 0.0 | 0.0 | 0.0 | 0.0 | 0.0 | 0.0 | 0.0 | 0.0 |
| Glucose | kg/h | 28,017 | 2,802 | 0.0 | 0.0 | 0.0 | 0.0 | 280.2 | 25,215 |
| Xylose | kg/h | 11,674 | 1,167 | 0.0 | 0.0 | 0.0 | 0.0 | 116.7 | 10,506 |
| Arabinose | kg/h | 2,001 | 200.1 | 0.0 | 0.0 | 0.0 | 0.0 | 20.0 | 1,801 |
| Cellobiose | kg/h | 341.2 | 34.1 | 0.0 | 0.0 | 0.0 | 0.0 | 34.1 | 307.1 |
| Glucose oligomers | kg/h | 1,832 | 183.2 | 0.0 | 0.0 | 0.0 | 0.0 | 183.2 | 1,649 |
| Xylose oligomers | kg/h | 961.5 | 96.2 | 0.0 | 0.0 | 0.0 | 0.0 | 96.2 | 865.4 |
| Arabinose oligomers | kg/h | 83.9 | 8.4 | 0.0 | 0.0 | 0.0 | 0.0 | 8.4 | 75.5 |
| Soluble lignin | kg/h | 501.5 | 50.1 | 0.0 | 0.0 | 0.0 | 0.0 | 50.1 | 451.3 |
| HMF | kg/h | 112.2 | 11.2 | 0.0 | 0.0 | 0.0 | 0.0 | 11.2 | 101.0 |
| Lactic acid | kg/h | 21.0 | 2.1 | 0.0 | 0.0 | 0.0 | 0.0 | 2.1 | 18.9 |
| Ammonia | kg/h | 0.1 | 0.0 | 40.2 | 0.3 | 0.0 | 0.0 | 4.7 | 0.1 |
| Sulfuric acid | kg/h | 0.0 | 0.0 | 0.0 | 0.0 | 0.0 | 0.0 | 0.0 | 0.0 |
| Ammonium acetate | kg/h | 69.2 | 6.9 | 0.0 | 0.0 | 0.0 | 0.0 | 6.9 | 62.3 |
| DAP | kg/h | 0.0 | 0.0 | 246.9 | 0.0 | 0.0 | 0.0 | 0.0 | 0.0 |
| Oil (lipids) | kg/h | 6.9 | 0.7 | 0.0 | 0.0 | 0.0 | 0.0 | 0.7 | 6.2 |
| $O_2$ | kg/h | 0.0 | 0.0 | 0.0 | 100.0 | 14,937 | 884.6 | 0.0 | 0.0 |
| $N_2$ | kg/h | 0.1 | 0.0 | 0.0 | 2,913 | 49,192 | 2,913 | 0.5 | 0.1 |
| $CO_2$ | kg/h | 0.4 | 0.0 | 0.0 | 1,092 | 0.0 | 0.0 | 8.4 | 0.4 |
| Cellulose | kg/h | 6.2 | 0.6 | 0.0 | 0.0 | 0.0 | 0.0 | 0.6 | 5.6 |
| Galactan | kg/h | 5.0 | 0.5 | 0.0 | 0.0 | 0.0 | 0.0 | 0.5 | 4.5 |
| Mannan | kg/h | 2.5 | 0.2 | 0.0 | 0.0 | 0.0 | 0.0 | 0.2 | 2.2 |
| Lignin | kg/h | 50.2 | 5.0 | 0.0 | 0.0 | 0.0 | 0.0 | 5.0 | 45.1 |
| Protein | kg/h | 13.0 | 1.3 | 0.0 | 0.0 | 0.0 | 0.0 | 1.3 | 11.7 |
| Ash | kg/h | 2.3 | 0.2 | 0.0 | 0.0 | 0.0 | 0.0 | 0.2 | 2.0 |
| Enzyme | kg/h | 1.4 | 0.1 | 0.0 | 0.0 | 0.0 | 0.0 | 0.1 | 1.3 |
| $H_2$ | kg/h | 0.0 | 0.0 | 0.0 | 0.0 | 0.0 | 0.0 | 0.0 | 0.0 |
| *P. putida* biomass | kg/h | 0.0 | 0.0 | 0.0 | 0.0 | 0.0 | 0.0 | 2,793 | 0.0 |
| Muconic acid | kg/h | 0.0 | 0.0 | 0.0 | 0.0 | 0.0 | 0.0 | 0.0 | 0.0 |
| Adipic acid | kg/h | 0.0 | 0.0 | 0.0 | 0.0 | 0.0 | 0.0 | 0.0 | 0.0 |
| Ammonium hydroxide | kg/h | 0.0 | 0.0 | 0.0 | 0.0 | 0.0 | 0.0 | 0.0 | 0.0 |
| Ammonium sulfate | kg/h | 0.0 | 0.0 | 0.0 | 0.0 | 0.0 | 0.0 | 0.0 | 0.0 |

*Abbreviations: DAP: diammonium phosphate; HMF: hydroxymethylfurfural.*

**Table S7** (continued)

| | | Stream number | | | | | | | |
|---|---|---|---|---|---|---|---|---|---|
| | Units | 9 | 10 | 11 | 12 | 13 | 14 | 15 | 16 |
| **Temperature** | **°C** | **37.6** | **25.0** | **25.0** | **35.0** | **38.1** | **38.1** | **38.1** | **25.0** |
| **Pressure** | **atm** | **4.6** | **1.0** | **1.0** | **1.3** | **1.0** | **1.0** | **1.0** | **1.0** |
| **Mass Flows** | **kg/h** | **61,536** | **812** | **7,413** | **64,174** | **424,329** | **22,168** | **402,162** | **14,693** |
| Water | kg/h | 1,205 | 0 | 0 | 1,578 | 386,711 | 10,374 | 376,337 | 1,029 |
| Ethanol | kg/h | 0.0 | 0.0 | 0.0 | 0.0 | 0.0 | 0.0 | 0.0 | 0.0 |
| Glucose | kg/h | 0.0 | 0.0 | 0.0 | 0.0 | 0.0 | 0.0 | 0.0 | 0.0 |
| Xylose | kg/h | 0.0 | 0.0 | 0.0 | 0.0 | 0.0 | 0.0 | 0.0 | 0.0 |
| Arabinose | kg/h | 0.0 | 0.0 | 0.0 | 0.0 | 0.0 | 0.0 | 0.0 | 0.0 |
| Cellobiose | kg/h | 0.0 | 0.0 | 0.0 | 0.0 | 341.2 | 9.2 | 332.1 | 0.0 |
| Glucose oligomers | kg/h | 0.0 | 0.0 | 0.0 | 0.0 | 1,832 | 49.1 | 1,783 | 0.0 |
| Xylose oligomers | kg/h | 0.0 | 0.0 | 0.0 | 0.0 | 961.5 | 25.8 | 935.7 | 0.0 |
| Arabinose oligomers | kg/h | 0.0 | 0.0 | 0.0 | 0.0 | 83.9 | 2.3 | 81.7 | 0.0 |
| Soluble lignin | kg/h | 0.0 | 0.0 | 0.0 | 0.0 | 501.5 | 13.5 | 488.0 | 0.0 |
| HMF | kg/h | 0.0 | 0.0 | 0.0 | 0.0 | 112.2 | 3.0 | 109.2 | 0.0 |
| Lactic acid | kg/h | 0.0 | 0.0 | 0.0 | 0.0 | 21.0 | 0.6 | 20.4 | 0.0 |
| Ammonia | kg/h | 0.0 | 113.8 | 0.0 | 0.4 | 4.4 | 0.1 | 4.3 | 0.0 |
| Sulfuric acid | kg/h | 0.0 | 0.0 | 0.0 | 0.0 | 0.0 | 0.0 | 0.0 | 13,665 |
| Ammonium acetate | kg/h | 0.0 | 0.0 | 0.0 | 0.0 | 69.2 | 1.9 | 67.4 | 0.0 |
| DAP | kg/h | 0.0 | 698.1 | 0.0 | 0.0 | 0.0 | 0.0 | 0.0 | 0.0 |
| Oil (lipids) | kg/h | 0.0 | 0.0 | 0.0 | 0.0 | 6.9 | 0.2 | 6.7 | 0.0 |
| $O_2$ | kg/h | 14,052 | 0.0 | 0.0 | 1,754 | 0.5 | 0.0 | 0.5 | 0.0 |
| $N_2$ | kg/h | 46,278 | 0.0 | 0.0 | 46,272 | 6.6 | 0.2 | 6.4 | 0.0 |
| $CO_2$ | kg/h | 0.0 | 0.0 | 0.0 | 14,562 | 94.5 | 2.5 | 92.0 | 0.0 |
| Cellulose | kg/h | 0.0 | 0.0 | 0.0 | 0.0 | 6.2 | 6.2 | 0.0 | 0.0 |
| Galactan | kg/h | 0.0 | 0.0 | 0.0 | 0.0 | 5.0 | 5.0 | 0.0 | 0.0 |
| Mannan | kg/h | 0.0 | 0.0 | 0.0 | 0.0 | 2.5 | 2.5 | 0.0 | 0.0 |
| Lignin | kg/h | 0.0 | 0.0 | 0.0 | 0.0 | 50.2 | 49.9 | 0.3 | 0.0 |
| Protein | kg/h | 0.0 | 0.0 | 0.0 | 0.0 | 13.0 | 13.0 | 0.1 | 0.0 |
| Ash | kg/h | 0.0 | 0.0 | 0.0 | 0.0 | 2.3 | 2.2 | 0.0 | 0.0 |
| Enzyme | kg/h | 0.0 | 0.0 | 0.0 | 0.0 | 1.4 | 1.4 | 0.0 | 0.0 |
| $H_2$ | kg/h | 0.0 | 0.0 | 0.0 | 0.0 | 0.0 | 0.0 | 0.0 | 0.0 |
| *P. putida* biomass | kg/h | 0.0 | 0.0 | 0.0 | 0.0 | 11,059 | 11,003 | 55.3 | 0.0 |
| Muconic acid | kg/h | 0.0 | 0.0 | 0.0 | 7.6 | 15,030 | 403.2 | 14,627 | 0.0 |
| Adipic acid | kg/h | 0.0 | 0.0 | 0.0 | 0.0 | 0.0 | 0.0 | 0.0 | 0.0 |
| Ammonium hydroxide | kg/h | 0.0 | 0.0 | 7,413 | 0.0 | 7,413 | 198.9 | 7,214 | 0.0 |
| Ammonium sulfate | kg/h | 0.0 | 0.0 | 0.0 | 0.0 | 0.0 | 0.0 | 0.0 | 0.0 |

**Table S7** (continued)

| | Units | Stream number | | | | | | | |
|---|---|---|---|---|---|---|---|---|---|
| | | 17 | 18 | 19 | 20 | 21 | 22 | 23 | 24 |
| **Temperature** | **°C** | **15.0** | **25.0** | **33.8** | **15.0** | **64.4** | **38.8** | **25.0** | **43.3** |
| **Pressure** | **atm** | **1.0** | **1.0** | **1.0** | **1.0** | **1.0** | **30.0** | **1.0** | **27.2** |
| **Mass Flows** | **kg/h** | **416,874** | **2,551** | **405,425** | **14,000** | **70,007** | **107** | **52** | **410** |
| Water | kg/h | 381,091 | 0 | 382,403 | 0 | 0 | 0 | 0 | 0 |
| Ethanol | kg/h | 0.0 | 0.0 | 0.0 | 0.0 | 56,000 | 0.0 | 51.7 | 0.0 |
| Glucose | kg/h | 0.0 | 0.0 | 0.0 | 0.0 | 0.0 | 0.0 | 0.0 | 0.0 |
| Xylose | kg/h | 0.0 | 0.0 | 0.0 | 0.0 | 0.0 | 0.0 | 0.0 | 0.0 |
| Arabinose | kg/h | 0.0 | 0.0 | 0.0 | 0.0 | 0.0 | 0.0 | 0.0 | 0.0 |
| Cellobiose | kg/h | 332.1 | 0.0 | 332.1 | 0.0 | 0.0 | 0.0 | 0.0 | 0.0 |
| Glucose oligomers | kg/h | 1,783 | 0.0 | 1,783 | 0.0 | 0.0 | 0.0 | 0.0 | 0.0 |
| Xylose oligomers | kg/h | 935.8 | 0.0 | 935.8 | 0.0 | 0.0 | 0.0 | 0.0 | 0.0 |
| Arabinose oligomers | kg/h | 81.7 | 0.0 | 81.7 | 0.0 | 0.0 | 0.0 | 0.0 | 0.0 |
| Soluble lignin | kg/h | 488.1 | 0.0 | 488.1 | 0.0 | 0.0 | 0.0 | 0.0 | 0.0 |
| HMF | kg/h | 109.2 | 0.0 | 109.2 | 0.0 | 0.0 | 0.0 | 0.0 | 0.0 |
| Lactic acid | kg/h | 20.4 | 0.0 | 20.4 | 0.0 | 0.0 | 0.0 | 0.0 | 0.0 |
| Ammonia | kg/h | 4.3 | 0.0 | 4.3 | 0.0 | 0.0 | 0.0 | 0.0 | 0.0 |
| Sulfuric acid | kg/h | 3,570 | 0.0 | 0.0 | 0.0 | 0.0 | 0.0 | 0.0 | 0.0 |
| Ammonium acetate | kg/h | 67.4 | 0.0 | 67.4 | 0.0 | 0.0 | 0.0 | 0.0 | 0.0 |
| DAP | kg/h | 0.0 | 0.0 | 0.0 | 0.0 | 0.0 | 0.0 | 0.0 | 0.0 |
| Oil (lipids) | kg/h | 6.7 | 0.0 | 6.7 | 0.0 | 0.0 | 0.0 | 0.0 | 0.0 |
| $O_2$ | kg/h | 0.5 | 0.0 | 0.5 | 0.0 | 0.0 | 0.0 | 0.0 | 0.0 |
| $N_2$ | kg/h | 6.4 | 0.0 | 6.4 | 0.0 | 0.0 | 0.0 | 0.0 | 0.0 |
| $CO_2$ | kg/h | 92.0 | 0.0 | 92.0 | 0.0 | 0.0 | 0.0 | 0.0 | 0.0 |
| Cellulose | kg/h | 0.0 | 0.0 | 0.0 | 0.0 | 0.0 | 0.0 | 0.0 | 0.0 |
| Galactan | kg/h | 0.0 | 0.0 | 0.0 | 0.0 | 0.0 | 0.0 | 0.0 | 0.0 |
| Mannan | kg/h | 0.0 | 0.0 | 0.0 | 0.0 | 0.0 | 0.0 | 0.0 | 0.0 |
| Lignin | kg/h | 0.3 | 0.0 | 0.3 | 0.0 | 0.0 | 0.0 | 0.0 | 0.0 |
| Protein | kg/h | 0.1 | 0.0 | 0.1 | 0.0 | 0.0 | 0.0 | 0.0 | 0.0 |
| Ash | kg/h | 0.0 | 0.0 | 0.0 | 0.0 | 0.0 | 0.0 | 0.0 | 0.0 |
| Enzyme | kg/h | 0.0 | 0.0 | 0.0 | 0.0 | 0.0 | 0.0 | 0.0 | 0.0 |
| $H_2$ | kg/h | 0.0 | 0.0 | 0.0 | 0.0 | 7.3 | 106.6 | 0.0 | 410.1 |
| *P. putida* biomass | kg/h | 55.3 | 0.0 | 55.3 | 0.0 | 0.0 | 0.0 | 0.0 | 0.0 |
| Muconic acid | kg/h | 14,628 | 0.0 | 627.7 | 14,000 | 14,000 | 0.0 | 0.0 | 0.0 |
| Adipic acid | kg/h | 0.0 | 0.0 | 0.0 | 0.0 | 0.0 | 0.0 | 0.0 | 0.0 |
| Ammonium hydroxide | kg/h | 0.0 | 2,551 | 0.0 | 0.0 | 0.0 | 0.0 | 0.0 | 0.0 |
| Ammonium sulfate | kg/h | 13,601 | 0.0 | 18,411 | 0.0 | 0.0 | 0.0 | 0.0 | 0.0 |

**Table S7** (continued)

| | Units | Stream number | | | | |
|---|---|---|---|---|---|---|
| | | 25 | 26 | 27 | 28 | 29 |
| **Temperature** | **°C** | **51.8** | **81.3** | **15.0** | **15.0** | **15.0** |
| **Pressure** | **atm** | **1.0** | **1.0** | **1.0** | **1.0** | **1.0** |
| **Mass Flows** | **kg/h** | **123,857** | **55,236** | **68,621** | **720** | **14,435** |
| Water | kg/h | 0 | 0 | 0 | 0 | 0 |
| Ethanol | kg/h | 104,243 | 55,228 | 49,015 | 719.9 | 37.9 |
| Glucose | kg/h | 0.0 | 0.0 | 0.0 | 0.0 | 0.0 |
| Xylose | kg/h | 0.0 | 0.0 | 0.0 | 0.0 | 0.0 |
| Arabinose | kg/h | 0.0 | 0.0 | 0.0 | 0.0 | 0.0 |
| Cellobiose | kg/h | 0.0 | 0.0 | 0.0 | 0.0 | 0.0 |
| Glucose oligomers | kg/h | 0.0 | 0.0 | 0.0 | 0.0 | 0.0 |
| Xylose oligomers | kg/h | 0.0 | 0.0 | 0.0 | 0.0 | 0.0 |
| Arabinose oligomers | kg/h | 0.0 | 0.0 | 0.0 | 0.0 | 0.0 |
| Soluble lignin | kg/h | 0.0 | 0.0 | 0.0 | 0.0 | 0.0 |
| HMF | kg/h | 0.0 | 0.0 | 0.0 | 0.0 | 0.0 |
| Lactic acid | kg/h | 0.0 | 0.0 | 0.0 | 0.0 | 0.0 |
| Ammonia | kg/h | 0.0 | 0.0 | 0.0 | 0.0 | 0.0 |
| Sulfuric acid | kg/h | 0.0 | 0.0 | 0.0 | 0.0 | 0.0 |
| Ammonium acetate | kg/h | 0.0 | 0.0 | 0.0 | 0.0 | 0.0 |
| DAP | kg/h | 0.0 | 0.0 | 0.0 | 0.0 | 0.0 |
| Oil (lipids) | kg/h | 0.0 | 0.0 | 0.0 | 0.0 | 0.0 |
| $O_2$ | kg/h | 0.0 | 0.0 | 0.0 | 0.0 | 0.0 |
| $N_2$ | kg/h | 0.0 | 0.0 | 0.0 | 0.0 | 0.0 |
| $CO_2$ | kg/h | 0.0 | 0.0 | 0.0 | 0.0 | 0.0 |
| Cellulose | kg/h | 0.0 | 0.0 | 0.0 | 0.0 | 0.0 |
| Galactan | kg/h | 0.0 | 0.0 | 0.0 | 0.0 | 0.0 |
| Mannan | kg/h | 0.0 | 0.0 | 0.0 | 0.0 | 0.0 |
| Lignin | kg/h | 0.0 | 0.0 | 0.0 | 0.0 | 0.0 |
| Protein | kg/h | 0.0 | 0.0 | 0.0 | 0.0 | 0.0 |
| Ash | kg/h | 0.0 | 0.0 | 0.0 | 0.0 | 0.0 |
| Enzyme | kg/h | 0.0 | 0.0 | 0.0 | 0.0 | 0.0 |
| $H_2$ | kg/h | 7.3 | 7.3 | 0.0 | 0.0 | 0.0 |
| *P. putida* biomass | kg/h | 0.0 | 0.0 | 0.0 | 0.0 | 0.0 |
| Muconic acid | kg/h | 0.0 | 0.0 | 0.0 | 0.0 | 0.0 |
| Adipic acid | kg/h | 19,606 | 0.0 | 19,606 | 0.0 | 14,397 |
| Ammonium hydroxide | kg/h | 0.0 | 0.0 | 0.0 | 0.0 | 0.0 |
| Ammonium sulfate | kg/h | 0.0 | 0.0 | 0.0 | 0.0 | 0.0 |

**Table S8. Main financial assumptions used in the TEA, based on an $n^{th}$-plant design.**

| **Financial Assumptions** | **Value** |
|---|---|
| Plant life | 30 years |
| Cost year dollar | 2020$ |
| Capacity Factor | 90% |
| Discount rate | 10% |
| General plant depreciation | MACRS |
| General plant recovery period | 7 years |
| Steam plant depreciation | MACRS |
| Steam plant recovery period | 20 years |
| Federal tax rate | 21% |
| Financing | 40% equity |
| Loan terms | 10-year loan at 8% APR |
| Construction period | 3 years |
| *First 12 months' expenditures* | 8% |
| *Next 12 months' expenditures* | 60% |
| *Last 12 months' expenditures* | 32% |
| Working capital | 5% of fixed capital investment |
| Start-up time | 6 months |
| *Revenues during start-up* | 50% |
| *Variable costs during start-up* | 75% |

APR: annual percentage rate; MACRS: Modified Accelerated Cost Recovery System

**Table S9. Typical inputs and outputs of the mixed sugar bioconversion process.** The list includes the associated purchase/selling prices (given in 2020$) for the case with the lowest minimum selling price (MSP) of adipic acid shown in the main text (strain DK062).

| **Input** | **Value** | **Price** |
|---|---|---|
| Mixed sugars* | 41,692 kg/h | $0.480/kg |
| Ammonium hydroxide ($NH_4OH$) | 9,964 kg/h | $0.239/kg |
| Sulfuric acid ($H_2SO_4$), 93 wt% | 14,693 kg/h | $0.046/kg |
| Ammonia ($NH_3$) | 154 kg/h | $0.56/kg |
| Diammonium phosphate (DAP) | 945 kg/h | $0.29/kg |
| Hydrogen ($H_2$) | 410 kg/h | $1.54/kg |
| Ethanol | 52 kg/h | $0.79/kg |
| Natural gas | 97.5 MMBTU/h | $1.81/MMBTU |
| Cooling water | 5.276,558 kg/h | $0.02/1,000 kg |
| Chiller water, 40ºF | 51 MMkcal/h | $5.09/GJ |
| Electricity | 5,962 kW | $0.082/kWh |
| **Output** | **Value** | **Price** |
| Adipic acid | 14,435 kg/h | *MSP of $2.74/kg* |

* Price obtained for the production of mixed sugars via a Deacetylation and Mechanical Refining (DMR) pretreatment of corn stover, further detailed in Mokwatlo *et al.* (2024) (*5*).

**Table S10. Breakdown of capital expenditures (CAPEX)** for the case with the lowest MSP of adipic acid: M9 medium, fermentation time of 96h, and use of DMR-based mixed sugars.

| | **Total cost (MM$)** |
|---|---|
| **Total Installed Costs** | **344.7** |
| *Fermentation* | *196.0* |
| *Muconic acid recovery* | *84.4* |
| *Upgrading to adipic acid* | *31.8* |
| *Adipic acid recovery* | *30.7* |
| *Other equipment* | *2.8* |
| **Other Direct Costs** | **60.3** |
| **Total Indirect Costs** | **243.0** |
| **Fixed Capital Investment** | **648.0** |

**Table S11. Life-cycle inventory for bio-based adipic acid production using final TRY metrics timepoint.**

| Strain | LC224* | LC224 | LC237 | DK062 | DK063 | TL207 |
|---|---|---|---|---|---|---|
| **Main product** | **kg/hr** | **kg/hr** | **kg/hr** | **kg/hr** | **kg/hr** | **kg/hr** |
| Adipic acid | 14,247 | 10,917 | 12,166 | 14,435 | 14,232 | 13,733 |
| **Resource Consumption** | **kg/hr** | **kg/hr** | **kg/hr** | **kg/hr** | **kg/hr** | **kg/hr** |
| DMR mixed sugars | 41,692 | 41,692 | 41,692 | 41,692 | 41,692 | 41,692 |
| Ammonia | 140 | 196 | 192 | 154 | 158 | 166 |
| Diammonium Phosphate | 857 | 1,212 | 1,187 | 945 | 966 | 1,019 |
| Sulfuric acid (93 wt%) | 14,512 | 11,249 | 12,470 | 14,693 | 14,494 | 14,005 |
| Ammonium hydroxide | 9,822 | 7,674 | 8,502 | 9,964 | 9,834 | 9,513 |
| Hydrogen | 404.7 | 310.3 | 346.2 | 410.1 | 404.3 | 390.2 |
| Ethanol | 51.0 | 39.5 | 45.1 | 51.7 | 51.0 | 49.3 |
| **Energy Use** | | | | | | |
| Electricity (kW) | 16,058 | 14,703 | 15,514 | 16,584 | 16,506 | 16,320 |
| Natural gas (MMkcal/h) | 24.3 | 19.6 | 21.3 | 24.6 | 24.3 | 23.6 |
| **Waste Generation** | **kg/hr** | **kg/hr** | **kg/hr** | **kg/hr** | **kg/hr** | **kg/hr** |
| Aqueous stream | 426,836 | 424,713 | 425,422 | 426,856 | 426,726 | 426,405 |

*LC224 values obtained from Ling *et al.* (2022) (*1*)

**Table S12. Life-cycle inventory for bio-based adipic acid production using maximum rate timepoint.**

| Strain | LC224* | LC224 | LC237 | DK062 | DK063 | TL207 |
|---|---|---|---|---|---|---|
| **Main product** | **kg/hr** | **kg/hr** | **kg/hr** | **kg/hr** | **kg/hr** | **kg/hr** |
| Adipic acid | 12,405 | 11,805 | 11,870 | 14,176 | 14,685 | 11,738 |
| **Resource Consumption** | **kg/hr** | **kg/hr** | **kg/hr** | **kg/hr** | **kg/hr** | **kg/hr** |
| DMR mixed sugars | 41,692 | 41,692 | 41,692 | 41,692 | 41,692 | 41,692 |
| Ammonia | 171 | 181 | 197 | 159 | 150 | 200 |
| Diammonium Phosphate | 1,053 | 1,117 | 1,218 | 972 | 918 | 1,232 |
| Sulfuric acid (93 wt%) | 12,707 | 12,119 | 12,179 | 14,440 | 14,939 | 12,050 |
| Ammonium hydroxide | 8,638 | 8,250 | 8,310 | 9,798 | 10,125 | 8,225 |
| Hydrogen | 353.0 | 335.4 | 337.8 | 402.7 | 417.2 | 334.1 |
| Ethanol | 45.9 | 41.4 | 44.1 | 50.8 | 52.6 | 43.7 |
| **Energy Use** | | | | | | |
| Electricity (kW) | 15,323 | 14,860 | 15,408 | 16,513 | 16,738 | 15,676 |
| Natural gas (MMkcal/h) | 21.7 | 20.8 | 20.9 | 24.2 | 24.9 | 20.7 |
| **Waste Generation** | **kg/hr** | **kg/hr** | **kg/hr** | **kg/hr** | **kg/hr** | **kg/hr** |
| Aqueous stream | 425,663 | 425,277 | 425,229 | 426,687 | 427,010 | 425,110 |

*LC224 values obtained from Ling *et al.* (2022) (*1*)

**Table S13. Life-cycle inventory for bio-based muconic acid production using final TRY metrics timepoint.**

| Strain | LC224* | LC224 | LC237 | DK062 | DK063 | TL207 |
|---|---|---|---|---|---|---|
| **Main product** | **kg/hr** | **kg/hr** | **kg/hr** | **kg/hr** | **kg/hr** | **kg/hr** |
| Adipic acid | 13,817 | 10,588 | 11,800 | 14,000 | 13,803 | 13,319 |
| **Resource Consumption** | **kg/hr** | **kg/hr** | **kg/hr** | **kg/hr** | **kg/hr** | **kg/hr** |
| DMR mixed sugars | 41,692 | 41,692 | 41,692 | 41,692 | 41,692 | 41,692 |
| Ammonia | 140 | 196 | 192 | 154 | 158 | 166 |
| Diammonium Phosphate | 857 | 1,212 | 1,187 | 945 | 966 | 1,019 |
| Sulfuric acid (93 wt%) | 14,512 | 11,249 | 12,470 | 14,693 | 14,494 | 14,005 |
| Ammonium hydroxide | 9,822 | 7,674 | 8,502 | 9,964 | 9,834 | 9,513 |
| **Energy Use** | | | | | | |
| Electricity (kW) | 14,569 | 13,543 | 14,229 | 15,077 | 15,019 | 14,882 |
| Natural gas (MMkcal/h) | 6.1 | 5.6 | 5.8 | 6.1 | 6.1 | 6.0 |
| **Waste Generation** | **kg/hr** | **kg/hr** | **kg/hr** | **kg/hr** | **kg/hr** | **kg/hr** |
| Aqueous stream | 426,836 | 424,713 | 425,422 | 426,856 | 426,726 | 426,405 |

*LC224 values obtained from Ling *et al.* (2022) (*1*)

**Table S14. Life-cycle inventory for bio-based muconic acid production using maximum rate timepoint.**

| Strain | LC224* | LC224 | LC237 | DK062 | DK063 | TL207 |
|---|---|---|---|---|---|---|
| **Main product** | **kg/hr** | **kg/hr** | **kg/hr** | **kg/hr** | **kg/hr** | **kg/hr** |
| Adipic acid | 12,031 | 11,449 | 11,512 | 13,749 | 14,243 | 11,385 |
| **Resource Consumption** | **kg/hr** | **kg/hr** | **kg/hr** | **kg/hr** | **kg/hr** | **kg/hr** |
| DMR mixed sugars | 41,692 | 41,692 | 41,692 | 41,692 | 41,692 | 41,692 |
| Ammonia | 171 | 181 | 197 | 159 | 150 | 200 |
| Diammonium Phosphate | 1,053 | 1,117 | 1,218 | 972 | 918 | 1,232 |
| Sulfuric acid (93 wt%) | 12,707 | 12,119 | 12,179 | 14,440 | 14,939 | 12,050 |
| Ammonium hydroxide | 8,638 | 8,250 | 8,310 | 9,798 | 10,125 | 8,225 |
| **Energy Use** | | | | | | |
| Electricity (kW) | 14,015 | 13,838 | 14,153 | 15,031 | 15,206 | 14,433 |
| Natural gas (MMkcal/h) | 5.9 | 5.8 | 5.7 | 6.1 | 6.2 | 5.7 |
| **Waste Generation** | **kg/hr** | **kg/hr** | **kg/hr** | **kg/hr** | **kg/hr** | **kg/hr** |
| Aqueous stream | 425,663 | 425,277 | 425,229 | 426,687 | 427,010 | 425,110 |

*LC224 values obtained from Ling et al. (2022) (*1*)
